# The Influence of Multivalent Charge and PEGylation on Shape Transitions in Fluid Lipid Assemblies: From Vesicles to Discs, Rods, and Spheres

**DOI:** 10.1101/2023.08.09.552538

**Authors:** Victoria M. Steffes, Zhening Zhang, Kai K. Ewert, Cyrus R. Safinya

## Abstract

Lipids, and cationic lipids in particular, are of interest as delivery vectors for hydrophobic drugs such as the cancer therapeutic paclitaxel, and the structures of lipid assemblies affect their efficacy. We investigated the effect of incorporating the multivalent cationic lipid MVL5 (+5*e*) and poly(ethylene glycol)-lipids (PEG-lipids), alone and in combination, on the structure of fluid-phase lipid assemblies of the charge-neutral lipid 1,2-dioleoyl-*sn*-glycero-phosphocholine (DOPC). This allowed us to elucidate lipid–liposome structure correlations in sonicated formulations with high charge density, which are not accessible with univalent lipids such as the well-studied DOTAP (+1*e*). Cryogenic TEM allowed us to determine the structure of the lipid assemblies, revealing diverse combinations of vesicles and disc-shaped, worm-like, and spherical micelles. Remarkably, MVL5 forms an essentially pure phase of disc micelles at 50 mol% MVL5. At higher (75 mol%) content of MVL5, short and intermediate-length worm-like micellar rods were observed and, in ternary mixtures with PEG-lipid, longer and highly flexible worm-like micelles formed. Independent of their length, the worm-like micelles coexisted with spherical micelles. In stark contrast, DOTAP forms mixtures of vesicles, disc micelles and spherical micelles at all studied compositions, even when combined with PEG-lipids. The observed similarities and differences in the effects of charge (multivalent versus univalent) and high curvature (multivalent charge versus PEG-lipid) on assembly structure provide insights into parameters that control the size of fluid lipid nanodiscs, relevant for future applications.

## Introduction

Liposomes are assemblies of amphiphilic molecules (lipids and surfactants) with a wide variety of applications.^1,2^ Cationic liposomes (CLs; consisting of mixtures of cationic and neutral lipids) have emerged as alternatives to engineered viral vectors for the delivery of therapeutic nucleic acids *in vitro* and *in vivo,*^3–5^ including in the clinic.^6–10^ In particular, this includes the successful deployment of ionizable (cationic) lipids in lipid nanoparticles (LNPs) as vectors for therapeutic siRNA in patisiran^9^ and for mRNA in the vaccines against SARS-CoV-2 in the worldwide pandemic in 2020.^11–14^

More broadly, liposomes are of great interest as carriers for therapeutics because they can solubilize small molecule hydrophobic drugs,^15^ target therapeutics to specific sites, and protect cargos that would otherwise be rapidly degraded or removed by the immune system.^16–19^ Hydrophobic cancer drugs, such as paclitaxel, require a carrier to overcome the problem of their low water solubility. These drugs are loaded in the nonpolar tail region of lipid carriers and can therefore be delivered by both vesicles and micelles.^20–23^ The carrier’s loading capacity is strongly affected by the state of the hydrophobic tails; membranes in the fluid or chain-melted state allow much greater drug loading than membranes in the chain-ordered state, because the solubility of PTX in the tail region is significantly higher.^24–30^

The structure, i.e., the size and shape, of liposomes affects their efficacy as drug carriers.^31^ For example, the very small size (4–5 nm) of micelles may facilitate passage through the proteoglycan layer to the cell membrane and thus enhance intracellular delivery.^32,33^ Ultimately, the structure of the self-assembled liposomes (vesicles and/or micelles) is dictated by the structure and physicochemical properties of the constituting lipids. However, the a priori prediction of assembly structure from lipid structure remains elusive, especially for lipid mixtures and less abundant assembly structures such as disc micelles. Therefore, elucidation of key physical and chemical properties of lipids that enable formation of stable disc micelles, whether kinetically or due to thermal equilibrium, is of high importance for hydrophobic drug delivery *in vitro* and *in vivo*.^34–39^

The steric size and the charge of lipid headgroups are two important parameters that correlate lipid molecular geometry (i.e. shape) and assembly structure. Lipids with a headgroup consisting of a water-soluble polymer chain, such as poly(ethylene glycol) lipids (PEG-lipids), provide an example of lipids forming high-curvature assemblies due to the steric size of their headgroup (see Figure 1). Meanwhile, cationic lipids tend to form high-curvature assemblies because the electrostatic repulsion between their headgroups increases their effective size. The greater the lipid’s headgroup charge, the greater the curvature of the lipid assembly; for instance, the highly charged multivalent cationic lipid MVLBG2 (+16*e*)^40^ forms pure phases of spherical micelles at 75−100 mol%.^41^

**Figure 1.**
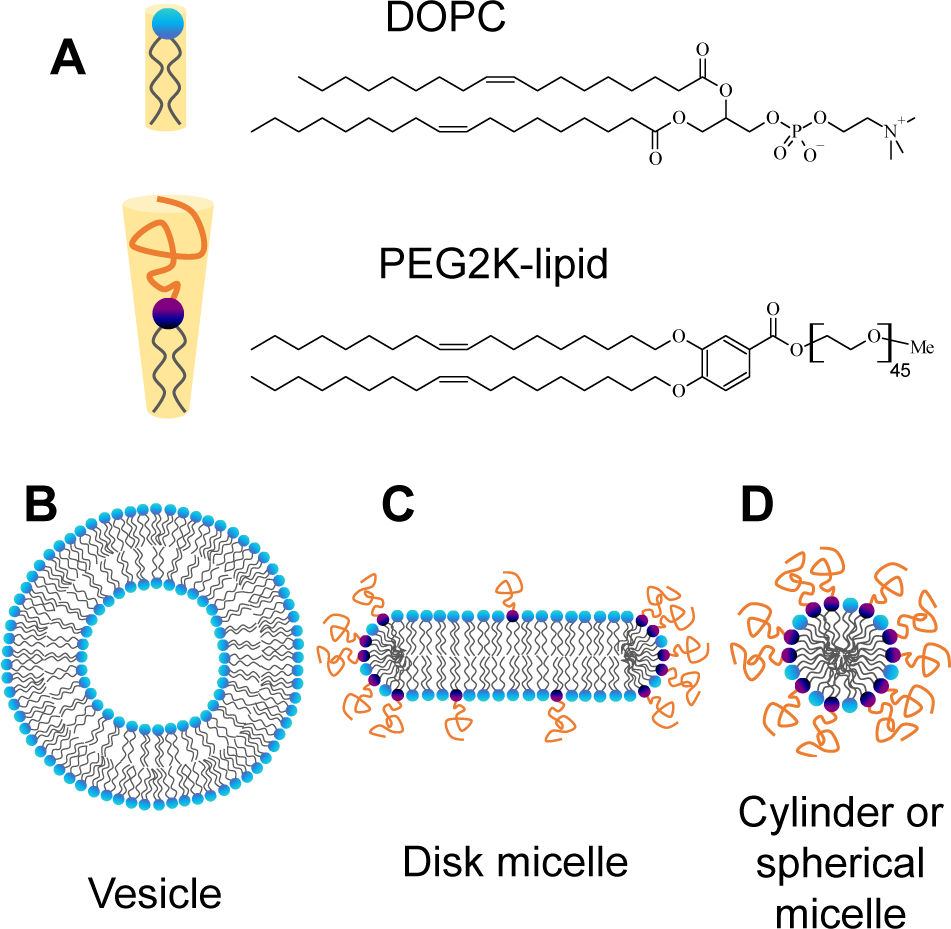
Relationship of lipid structure, molecular shape, and self-assembly according to relative spontaneous curvature. (A) Chemical structures of DOPC and PEG2K-lipid with schematic depictions of their molecular shape. (B–D) Schematic cross-sections of a vesicle, disc micelle and spherical/cylindrical micelle, illustrating the effect and distribution of high-curvature lipids in the self-assemblies.

Cationic charge and steric stabilization by PEGylation are also two important properties of lipid assemblies for drug delivery. For example, cationic lipid vectors have been observed to target tumor neovasculature, which has a greater negative charge than other tissues in the body.^42–47^ The addition of a hydrophilic PEG corona by incorporation of PEG-lipids into liposomes, in turn, is widely used in drug delivery applications to increase circulation time.^4,15,48^ Both the amount of PEG-lipid and the PEG chain length are known to effect stability, structure, and *in vivo* circulation lifetime of lipid-based carriers.^49,50^

In a previous cryo-TEM study, the structures of fluid lipid assemblies of the charge-neutral (zwitterionic) lipid 1,2-dioleoyl-*sn*-glycero-3-phosphocholine (DOPC) mixed with univalent cationic lipid (DOTAP(+1*e*)) (2,3-dioleyloxypropyltrimethylammonium chloride, headgroup charge +1*e*; see Figure S1 in the Supporting Information) or PEG2K-lipid (Figure 1, PEG molecular weight ≈2000 g/mol, n≈45), were investigated (Figure 2).^34^ DOTAP has been used for delivery of nucleic acids and hydrophobic cancer drugs,^4^ and a formulation of 50 mol% DOTAP, 47 mol% DOPC, and 3 mol% PTX (EndoTAG-1) has been investigated in clinical trials of patients with advanced triple-negative breast cancer.^51^ Cryo-TEM revealed that addition of 50 mol% DOTAP or 10 mol% PEG2K-lipid (with PEG chains in the brush conformation) to neutral vesicles of DOPC led to the unexpected emergence of new fluid lipid assembly structures. Small vesicles with diameters down to 10–20 nm coexisted with micellar discs (Figure 2) and occasional speres.^34^

**Figure 2.**
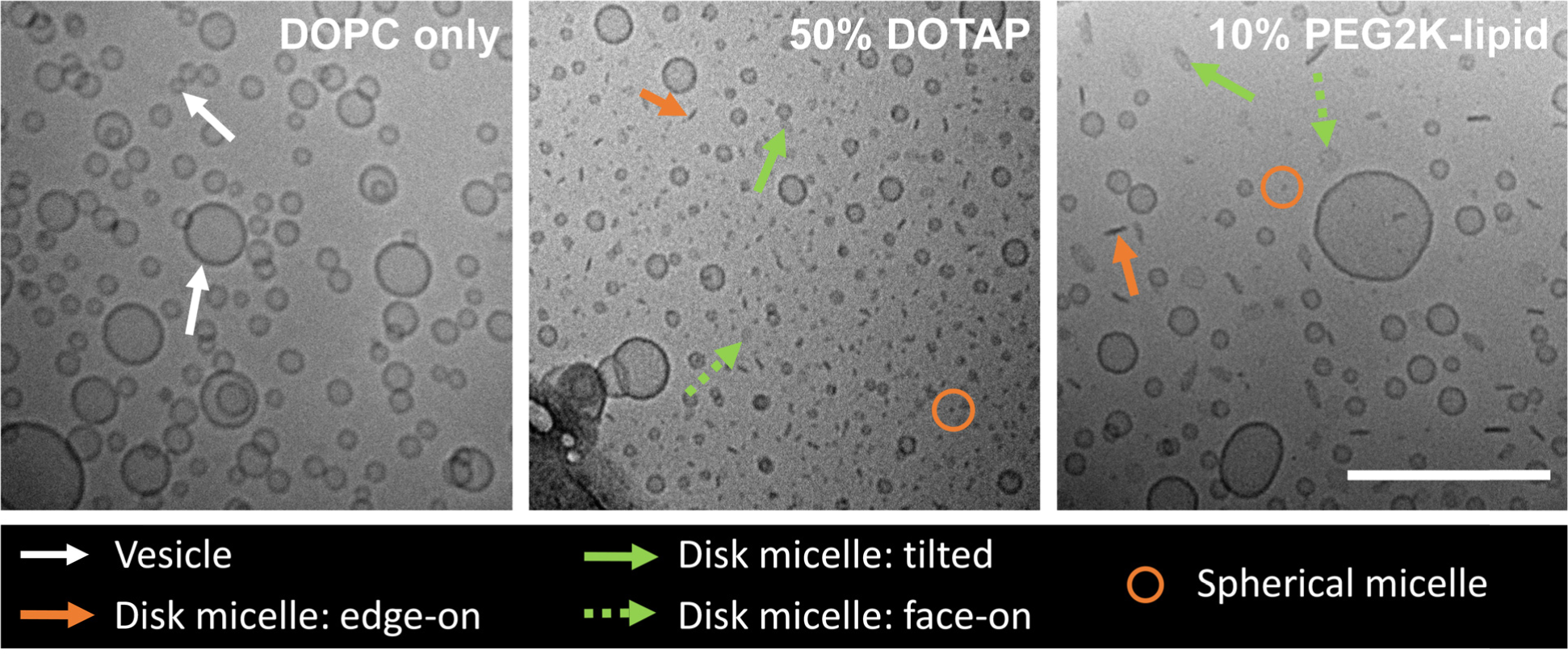
Effect of incorporation of univalent cati nic lipid or PEG2K-lipid on lipid assembly structure. Cryo-TEM reveals the structures present in formulations containing neutral DOPC alone (left), DOPC and 50 mol% DOTAP(+) (center), and DOPC and 10 mol% PEG2K-lipid (right); all formulations also contained 3 mol% PTX. Arrows and circles point out representative vesicle and micelle structures as designated in the legend. The majo ity of the nonvesicular objects are disc micelles, seen in tilted (green arrow), face-on (green dashed arrow) and edge-on (red arrow) orientation. Scale bar: 200 nm. Images adapted with permission from ref. ^34^

To further the understanding of lipid–liposome structure correlations in extremely high charge-density lipid assemblies inaccessible with univalent lipids, we employed MVL5 (+5*e*, see Figure S1 for the chemical structure)^52^ as a model multivalent cationic lipid in the present study. The high local concentration of charges in membranes containing multivalent lipids can lead to unique phenomena.^53^ MVL5 is commercially available and has been studied as a component of various lipid and lipid–nucleic acid nanostructures.^53–57^ As additional high-curvature lipids, we studied PEG-lipids (with PEG molecular weight of 2000 and 5000 g/mol), including in ternary mixtures with cationic lipid. This allowed differentiation between effects due to charge and steric size of lipid headgroups on the shape transitions in lipid assemblies. All lipids had oleyl tails (with one *cis* double bond), which exclusively form the chain-melted phase with high chain mobility and PTX solubility at room temperature.

We used cryogenic TEM (cryo-TEM) to determine the structures, shapes, and sizes of the lipid assemblies. Most of the studied formulations exhibited coexistence of multiple types of structures, forming diverse combinations of vesicles with discs, intermediate-length rods, worm-like and spherical micelles. These findings are consistent with all studied lipids (aside from DOPC) possessing a positive spontaneous curvature. Remarkably, MVL5 forms an essentially pure phase of disc micelles at 50 mol%. At higher content (75 mol%) of MVL5, spherical micelles coexist with short and intermediate-length worm-like micellar rods and, in ternary mixtures including PEG-lipid, with longer and more flexible worm-like micelles. Univalent DOTAP, in contrast, forms mixtures of vesicles, disc micelles and spherical micelles at all compositions studied (including when combined with PEG-lipid). While both PEG-lipids and cationic lipids form structures expected for lipids with high spontaneous curvature, they also exert distinctly different effects, in particular on the size of the fluid disc micelles. The resulting insights into parameters that control the size of lipid nanodiscs are relevant for future applications.

## Results

This study was, in part, designed to investigate lipid-based carriers of the hydrophobic drug paclitaxel (PTX). For this reason, several of the samples included 3 mol% PTX. However, control samples without PTX revealed that the PTX has no effect on lipid assembly structure. We therefore do not discuss the presence of PTX in the following section, but have included the exact lipid compositions in the figure captions and the Supporting Information (Table S1).

We used cryogenic TEM (cryo-TEM) to determine the structures of the lipid assemblies. Cryo-TEM is a powerful tool for this purpose, particularly when multiple types of structures coexist. However, other than for bulk techniques such as light and x-ray scattering, sampling bias toward small structures can be a concern with cryo-TEM. In this work, all lipid formulations were suspended in deionized water and sonicated 24 h prior to vitrification. Sonication of the samples ensured that large particles are rare. In addition, we analyzed both low- and high-magnification images for all samples.

### Binary Lipid Mixtures Containing a Multivalent Cationic Lipid

We investigated the effect of higher headgroup charge on lipid assembly structure by studying mixtures of DOPC and the multivalent cationic lipid MVL5^52^ (maximum headgroup charge +5*e*, see Figure S1 in the Supporting Information). Multivalent lipids enable access to very high membrane charge densities.^58^ In addition, the high local concentration of charge in the headgroup of multivalent lipids gives rise to unique effects.^53^ MVL5 is expected to be a cone-shaped lipid (with a headgroup area that is larger than that of the tails) giving rise to a large spontaneous curvature. This high spontaneous curvature is due to the increased steric size of the headgroup (compared with DOTAP or DOPC) and the intermolecular electrostatic repulsion between the highly charged headgroups.

We studied lipid formulations containing a low, intermediate, and high amount of MVL5 (10, 50, and 75 mol%, respectively) together with DOPC. Representative cryo-TEM images are shown in Figure 3, and unlabeled and additional images are provided in Figures S2–S7. For the control formulation of DOPC alone (Figure 2), cryo-TEM images showed an abundance of spherical unilamellar vesicles with diameters ranging from 12−200 nm, with the majority of vesicles 18–60 nm in diameter; some structures as large as 500 nm are visible in low-magnification images (Figures S8, S9).

**Figure 3.**
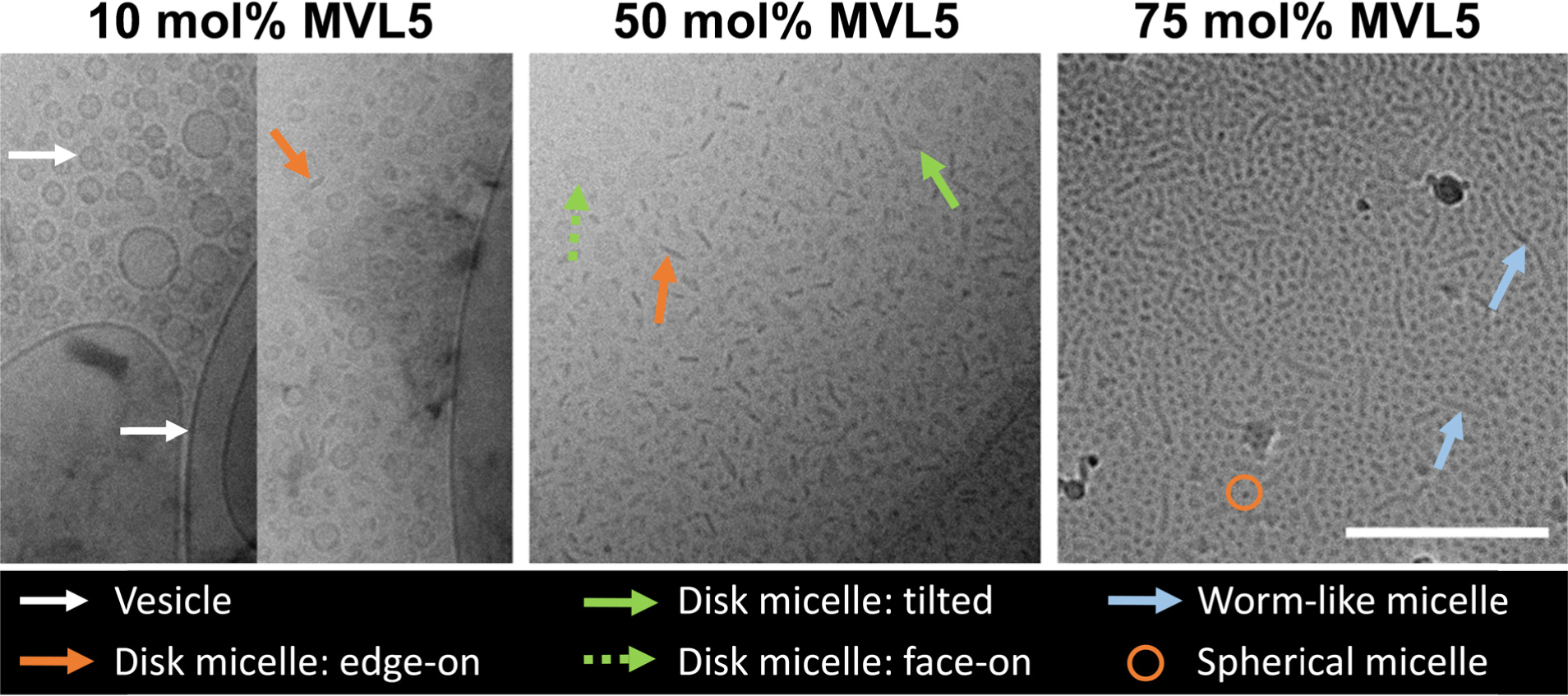
Structures of lipid assemblies containing multivalent MVL5. The cryo-TEM images show how the high curvature cationic lipid MVL5 changes lipid assembly structure as a function of composition (mol% of MVL5 in the formulation) from vesicles to discs, to short/intermediate length worm-like micellar rods coexisting with spheres. Arrows and circles identify vesicle and micelle structures according to the legend. The formulation with 10 mol% MVL5 only showed presence of discs coexisting with vesicles *in a single* out of 38 micrographs (right rectangular panel), with all other micrographs showing solely vesicles. Lipid composition: MVL5 content as specified; remainder: DOPC. Scale bar: 200 nm. For additional and unlabeled micrographs, see Figures S2–S7 in the Supporting Information.

In the formulation with 10 mol% MVL5, vesicles of a wide range of sizes were observed (12–220 nm, with the majority between 13 and 85 nm; up to >1 μm in low-magnification images; see Figures S2, S3). Disc micelles (17–25 nm diameter) were only observed in a *single* micrograph out of 38 taken (Figure 3, 10 mol% MVL5, right panel). This is in contrast to formulations containing 50 mol% univalent cationic lipid DOTAP(+1*e*) with similar charge density, where mixtures of vesicles and discs were observed in most micrographs (Figure 2).^34^

A remarkable change in structure is observed in the formulation with 50 mol% MVL5 (Figures 3 and S5). No vesicles were visible in high-resolution cryo-EM images, and the only structures observed in the micrographs are disc micelles in various orientations: tilted (elliptical objects, solid green arrow), edge-on (rod-like objects, orange arrow), and face-on (dashed green arrow). This proves unambiguously that the micelles are discs rather than rods. Most of the discs have diameters between 10 and 30 nm. However, discs as small as 7.5 nm are present, and larger discs, including large objects with irregular shape, can be found; unlike the smaller discs, these occasionally are not flat. The only vesicles observed were a few large (150–200 nm) ones in low-magnification images, see Figure S4.

When the charge density of the lipid membranes is increased even further, at 75 mol% MVL5, the observed assembly structures again change entirely (Figure 3, right panel). No disc micelles or vesicles are observed (including in low-magnification images, see Figure S6). Instead, the cryo-EM micrographs reveal a mixture of short and intermediate length worm-like micellar rods and spherical micelles. While most of the worm-like micelles are 20–100 nm in length, occasional examples are very long (>300 nm) (Figures 3 and S7).

### Binary Lipid Mixtures Containing PEG-Lipids

PEG-lipids with PEG chains of molecular weight 2000 and 5000 g/mol are important in biomedical applications. Because of the steric bulk of the PEG chain in their headgroup, PEG-lipids have a large headgroup. The size of the headgroup increases with the length of the PEG chain (due to an increase in the radius of gyration of PEG and thus the headgroup area). Previous work has revealed that PEG-lipid is able to promote the formation of disc-shaped micelles in dispersions of lipids that form membranes in the chain-ordered (“gel phase”) state. Discs were observed over a range of PEG-lipid composition (5–30 mol%),^59^ but in particular at high levels of PEG-lipid in the membrane (>20 mol%).^60^ Separately, worm-like micelles were observed under certain conditions in PEGylated cationic liposomes complexed with DNA.^61^

To gain further insight on how PEG length (2000 and 5000 g/mol molecular weight) affects the structure of *fluid-phase* lipid assemblies, we expanded on earlier studies of mixtures of DOPC with 10 mol% PEG2K-lipid. These prior studies revealed mixtures of smaller liposomes coexisting with disc micelles in the fluid membrane state (Figure 2) and showed that incorporation of 10 mol% PEG2K-lipid into DOTAP-based vectors of PTX increased their efficacy.^34^ In the present work, we prepared formulations of DOPC at both lower (2 mol%) and higher (25 mol%) content of PEG-lipids. The higher PEG-lipid contents, in particular, are comparable to those used in solid lipid nanodiscs.

At 2 mol% PEG2K-lipid, only vesicles are observed (Figures 4 and S10, S11). The only difference to vesicles of pure DOPC is the near-absence of vesicles larger than 300 nm in size, even in low-magnification images. For the previously reported formulation with 10 mol% PEG2K-lipid (Figures 2, S12, S13), vesicles (14 to ∼110 nm diameter, mostly <60 nm; up to ∼350 nm diameter in low-magnification images) constituted the majority of the objects. However, these vesicles coexisted with disc micelles having diameters between ≈12 and ≈30 nm (i.e., larger than the discs observed at 50 and 80 mol% DOTAP; see Figures 2 and S15-17) and a few objects compatible with being short rods and spheres.

**Figure 4.**
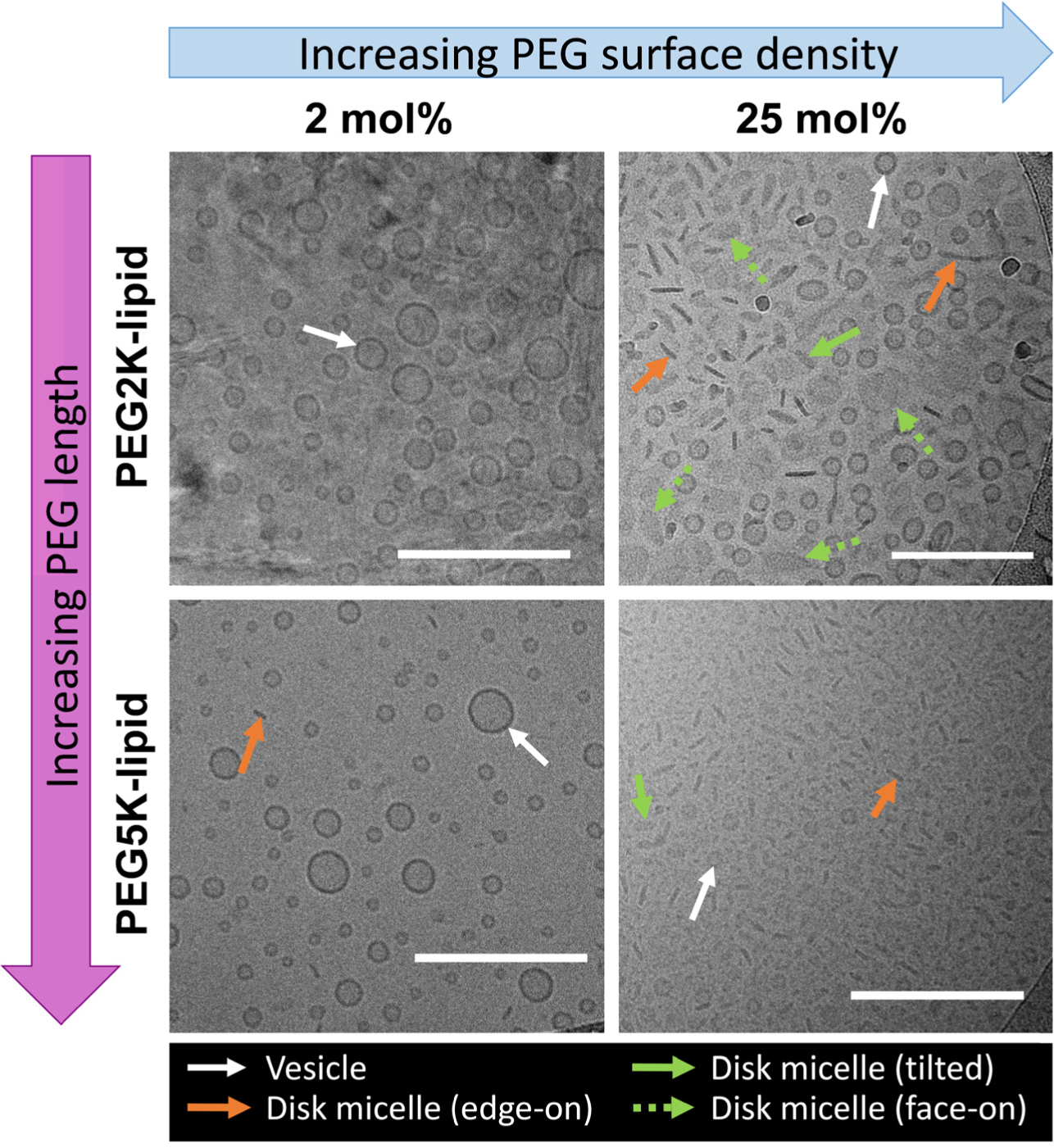
Effect of PEG-lipid content and PEG chain length on lipid assembly structure. Cryo-TEM images reveal the structures of formulations containing 2 and 25 mol% PEG-lipid with two PEG chain lengths (2000 and 5000 g/mol, see Figure S1). Arrows identify representative vesicle and micelle structures according to the legend. Lipid compositions: PEG-lipid content as specified along the image axes, remainder DOPC. Scale bars: 200 nm. For additional and unlabeled micrographs, see Figures S10, S11, and S18–S23.

At 25 mol% PEG2K-lipid (Figures 4, S18, S19), the population of disc micelles relative to vesicles has increased when compared to 10 mol% PEG2K-lipid. Discs of a wide range of sizes (diameters from <10 nm to a few >100 nm) are visible, and a number of these have irregular (noncircular) shape. The micrographs only show a few objects that may be spherical micelles. There are more and larger discs and fewer very small vesicles and spherical micelles compared to formulations containing 50 mol% univalent DOTAP (Figures 2, S15). The diameter of the vesicles varies mostly between ≈15 nm and ≈65 nm (Figures 4, S19), and even in low-magnification images (Figure S18), no vesicles of diameter >100 nm are seen.

For the two formulations (2 and 25 mol%) with a PEG-lipid with a longer PEG chain (PEG5K-lipid, PEG molecular weight 5000 g/mol), we used a commercially available lipid that also carries a negative charge at the phosphate moiety of the headgroup (see Figure S1 in the Supporting Information). Even at 2 mol% of the PEG5K-lipid (Figures 4, S20, S21), occasional disc (or rod) micelles (15–20 nm in diameter) coexisted with spherical vesicles of a variety of sizes up to 350 nm in diameter. However, no spherical micelles are observed. The observation of discs at 2 mol% of PEG5K-lipid but not PEG2K-lipid is consistent with the fact that less PEG5K-lipid is required to reach monolayer coverage of the membrane by the PEG chains.^62,63^ Repulsive interactions between PEG chains, near the mushroom to brush conformation transition on the planar segment of the disc, likely are the force driving lateral phase separation of PEG-lipid. These interactions eventually lead to formation of disc micelles with PEG-lipid primarily present on their edges (Figure 1) where a greater volume of space is available to each chain.

At 25 mol% PEG5K-lipid, the relative abundance of vesicles and discs (or rods) is reversed: only a few small vesicles are present while the majority of structures are discs (rods). The size of the vesicles (20–45 nm diameter in high magnification images, see Figures 4 and S22; even less abundant but with sizes up to 250 nm diameter in low-magnification images, see Figure S23) has decreased compared to 2 mol% PEG5K-lipid. A wider distribution of micelle sizes is present, from spherical micelle size to discs of up to 35 nm diameter, with the majority between ≈15 and ≈30 nm in size. Thus, the discs are larger at 25 mol% than at 2 mol% PEG5K-lipid, which is consistent with the trend observed for the PEG2K-lipid in going from 10 mol% (Figure 2) to 25 mol% (Figure 4). Compared to the formulation with 25 mol% PEG2K-lipid, fewer vesicles are observed at 25 mol% PEG5K-lipid, and the size of the discs is smaller.

### Ternary Lipid Mixtures: Combining Charge and PEGylation-Mediated Steric Repulsion

To examine the combined effects of multivalent charge and PEGylation on the shape of lipid assemblies, we studied ternary mixtures of 10 mol% PEG2K-lipid, multivalent MVL5, and DOPC. As evident in Figure 5 (and Figures S24, S25), no drastic effects on disc and vesicle size were evident upon addition of 10 mol% MVL5 (top left panel) to a formulation with 10 mol% PEG2K-lipid (bottom left panel). This finding is consistent with the fact that the formulation containing 10 mol% MVL5 and 90 mol% DOPC near-exclusively formed vesicles (Figure 3, left panel).

**Figure 5.**
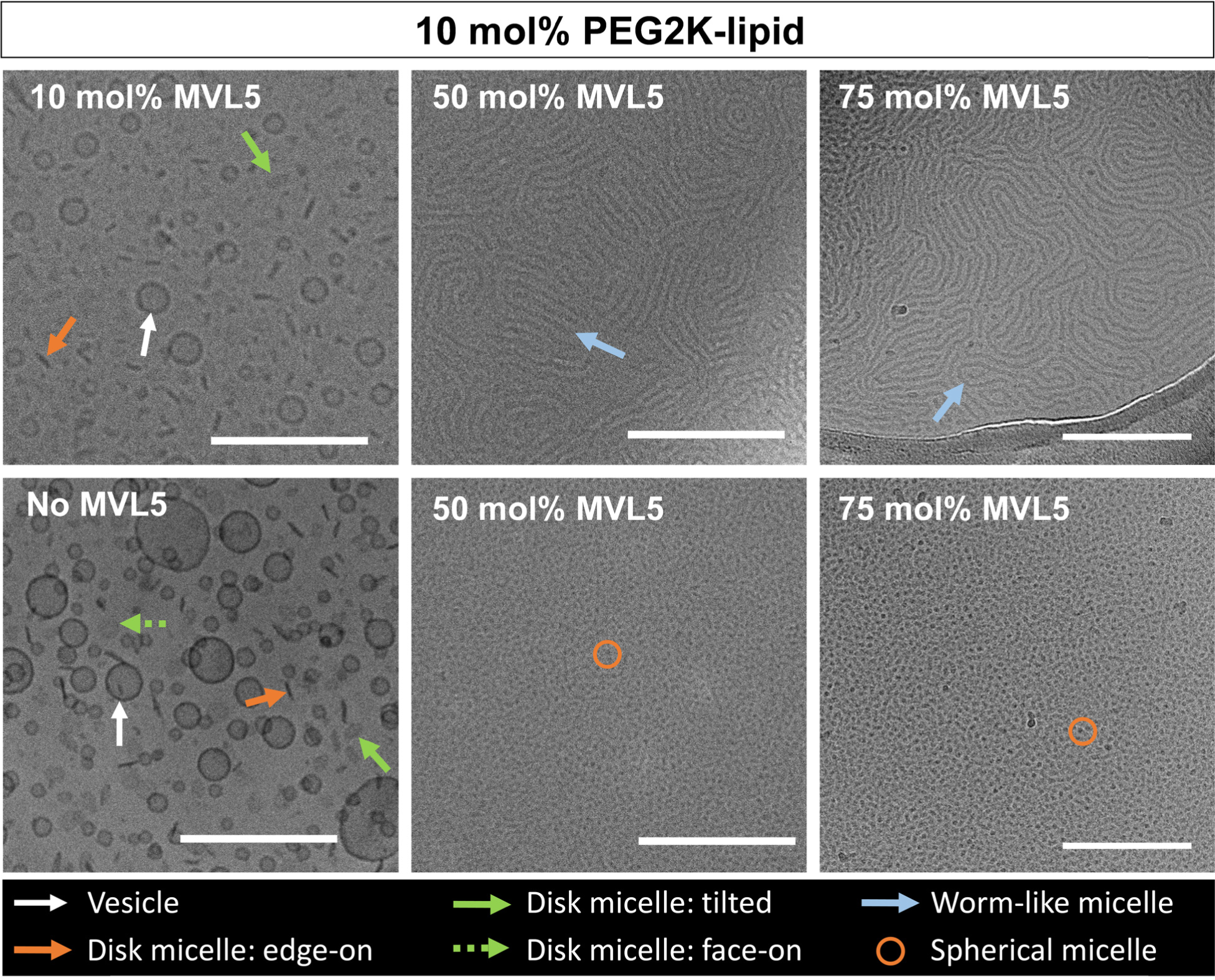
Structures of lipid assemblies in ternary formulations containing multivalent MVL5, PEG2K-lipid, and DOPC. The cryo-TEM images show the dramatic change in assembly structure when larger mol fractions of MVL5 are combined with 10 mol% PEG2K-lipid. For comparison, see the binary formulations (10 mol% PEG2K-lipid: bottom left panel; MVL5: see Figure 3). No other formulations in this study showed the relatively long and more flexible cylindrical micelles or abundance of spherical micelles observed in the samples containing 75 mol% MVL5 and 10 mol% PEG2K-lipid. Arrows and circles identify vesicle and micelle structures according to the legend. Lipid composition: 10 mol% PEG2K-lipid, MVL5 content as specified on the images, remainder DOPC. Scale bars: 200 nm. For additional and unlabeled micrographs, see Figures S12, S13, and S24–S30 in the Supporting Information.

As observed in the samples without PEG2K-lipid, incorporating larger amounts of multivalent MVL5 has a drastic effect on lipid assembly structure. Remarkably, rather than the discs observed at 50 mol% MVL5 without PEG-lipid, medium-length cylindrical micelles are the dominant structure observed in the formulation containing 10 mol% PEG2K-lipid and 50 mol% MVL5. These flexible worm-like micelles coexist with spherical micelles (Figures 5, S26, and S27).

At even higher charge density at 75 mol% MVL5 (and 10 mol% PEG2K-lipid), spherical micelles are more abundant and coexist with cylindrical micelles (Figure 5, S28–S30). While a wide range of lengths is observed for the cylindrical micelles, they are significantly longer than those found in the sample without PEG2K-lipid (Figure 3). No other formulations in this study showed the flexible cylindrical micelles or abundance of spherical micelles shown in this sample.

Interestingly, the micrographs show a spatial separation of assembly structures in the PEGylated samples with high (50 and 75 mol%) MVL5 content, i.e., only spherical micelles are observed in some images, and only cylindrical micelles in others. This may occur because similar shapes pack together more easily at high concentration, forming regularly spaced secondary structures (e.g., the worm-like micelles align parallel to one another). Alternatively, in the vitrified samples for cryo-TEM, ice thickness may determine where particles are observed as a function of their size. This is most convincingly evident in micrographs of the formulation containing 50 mol% DOTAP, shown in Figure S31.

A remarkable finding is that the effect of adding charged lipids to neutral PEGylated liposomes strongly depended on the valence. For example, the inclusion of 10 mol% MVL5 had little effect on the shapes and dimensions of observed lipid assemblies with 10 mol% PEG2K-lipid in DOPC. This is in strong contrast to the effect of addition of DOTAP on lipid assembly size as observed in earlier studies (see Figures S32–S34 and ref. ^34^). Incorporation of 50 mol% DOTAP (with a similar average charge density to 10 mol% MVL5) increased the fraction of disc micelles relative to vesicles, and their size (6–19 nm) **d**ecreased, as did the size of the vesicles (7–85 nm with the majority 9–42 nm, see Figure S33; up to 630 nm in lower-magnification images, see Figure S34). This trend continued for a formulation with 80 mol% DOTAP and 10 mol% PEG2K-lipid (Figures S32, S35, and S36), in particular with respect to the size of vesicles and disc micelles. In this sample, very small discs down to the size of spherical micelles are quite abundan. The effect of incorporating 50 mol% DOTAP was similar at a PEG2K-lipid content of 25 mol% (Figure 6 and Figures S22, S23, S37, and S38): larger disc micelles all but disappeared in favor of small and very small discs, and smaller vesicles and spherical micelles became abundan.

**Figure 6.**
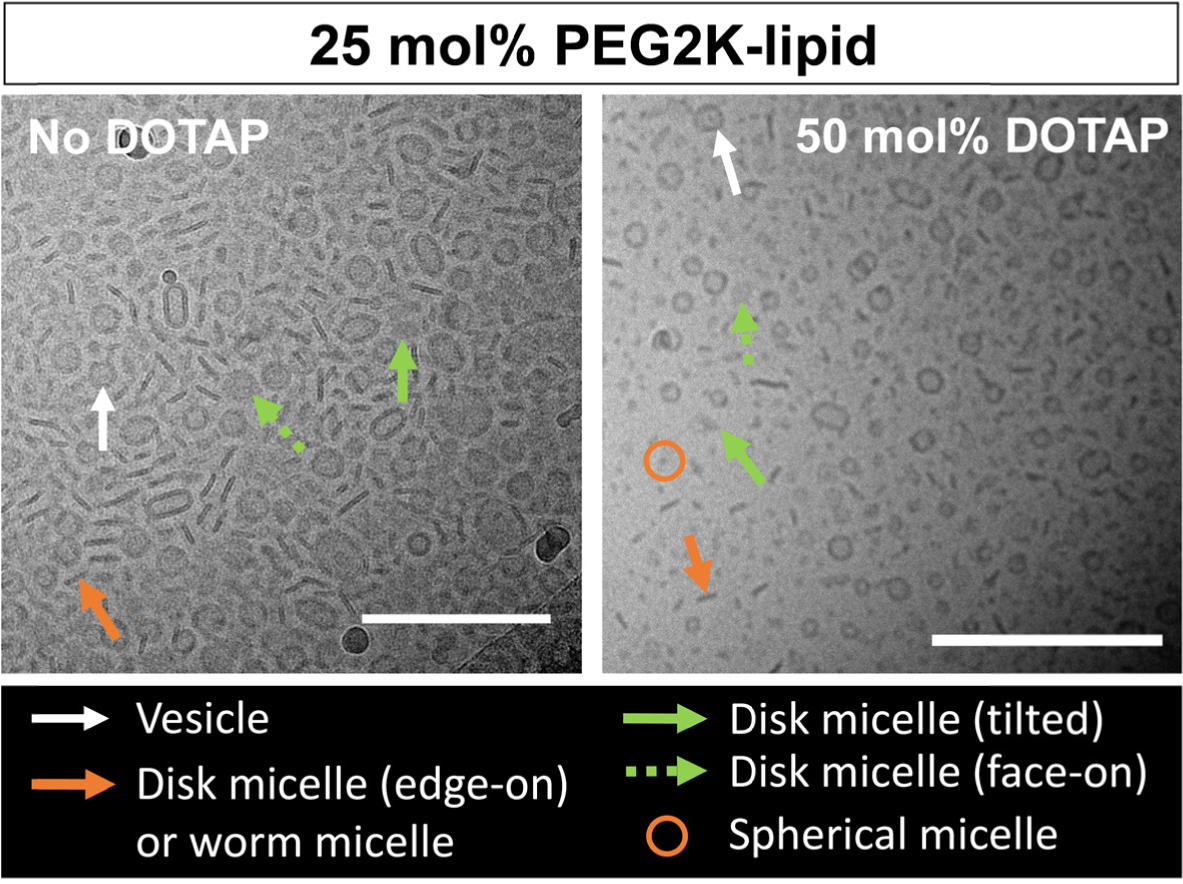
Cryo-TEM images demonstrating the effect of added univalent cationic DOTAP on the lipid assembly structures of a formulation with a high content of PEG2K-lipid. The formulations contain DOTAP as specified and 25 mol% PEG2K-lipid, with DOPC the remainder. Arrows and circles identify vesicle and micelle structures according to the legend. Scale bars: 200 nm. For unlabeled and additional micrographs, see Figures S18, S19, S37, and S38.

While the incorporation of both PEG-lipid and univalent lipid leads to discs coexisting with vesicles, an important overall trend is that PEGylated samples show suppressed formation of very large vesicles. Even in low-magnification images, no very large vesicles were observed in the ternary (PEG2K-lipid, DOTAP and DOPC) and binary (DOPC and PEG2K-lipid or PEG5K-lipid) formulations with PEG-lipids, in contrast to binary formulations of DOTAP and DOPC.

## Discussion

Lipid-based carriers are of great interest for the delivery of hydrophobic drugs, and their size and shape can strongly affect efficacy. For example, recent cell-based studies have shown that the small size and shape anisotropy of micellar discs enhance cell uptake of these vectors^34–36,39^ because distinct additional endocytic pathways are available to them: nanodiscs were found to use macropinocytosis and microtubule-mediated endocytosis in addition to the typical clathrin- and caveolae-mediated endocytic pathways used by larger spherical lipid vectors.^39^ Consistent with these *in vitro* results, formulations of PTX-loaded coexisting nanometer-scale fluid lipid discs and vesicles showed tumor penetration and proapoptotic activity *in vivo* in a model of triple-negative breast cancer in immunocompetent mice.^38^

It is important to point out that all lipid assemblies studied in the present work have membranes that were in the fluid state. This contrasts with previous studies that have characterized the structure and properties of *solid* lipid nanodiscs,^35,36,39,59^ where the (charge-neutral or slightly anionic) assemblies of lipids with saturated tails form the chain-ordered “gel” phase^64^ with very low chain mobility. The advantage of the *fluid* membranes investigated in the present study is that they are much more capable of solubilizing hydrophobic drugs such as PTX than the 2D-crystalline membranes of *solid* lipid nanodiscs.

To be able to take full advantage of the efficacy improvements for lipid carriers of varied shapes, it would be desirable to understand the parameters that affect lipid assembly size and shape. While current theoretical approaches are satisfactory for rationalizing the occurrence of certain structures and categorizing lipids after determining the structures they form in aqueous suspension, they are rarely able to predict assembly structure a priori from the molecular structure of the lipid.^65^ In particular, it is unclear whether disc micelles should be considered an equilibrium phase, and difficulty in predicting the extent of phase coexistence (e.g. vesicles mixed with discs) means that there are currently no design principles for achieving pure phases of intermediate-curvature structures such as discs and cylinders. Below, we consider the prevalent theoretical approaches of relating the structures of lipids and lipid assemblies and apply them to our findings for the formulations with multivalent charged lipids and PEG-lipids.

An early, molecular approach used geometric considerations of how lipids of distinct shape would self-assemble in water to minimize exposure of their hydrophobic tails to define the dimensionless “packing parameter”, *P* = *v*/*a*_0_*l*_c_. Here, *v* and *l*_c_ are the volume and maximum effective length of the tails, and *a*_0_ is the optimal headgroup area.^66,67^ The packing parameter thus categorizes lipid shape and the expected assembly structures by the ratio of the areas of the headgroup (*a*_0_) and the hydrophobic tail(s) (i.e., *v*/*l*_c_). Within this molecular packing model, bilayers (e.g. vesicles), micellar cylinders, and spheres are predicted to exist for ½<*P*<1, 1/3<*P*<1/2, and 0<*P*<1/3, respectively. The model does not predict micellar discs, which are observed experimentally but poorly understood theoretically. Nevertheless, one may expect from packing considerations that discs would appear when the packing parameter falls in a range between that of bilayers and micellar cylinders.

The shape of lipids can also be related to the spontaneous curvature (*C*_0_) of a lipid assembly in Helfrich’s formulation of the membrane curvature elastic energy per unit area (*F*/*A*):^68,69^

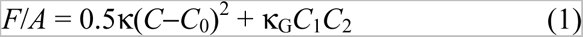

Here, *C*_1_ = 1/*R*_1_ and *C*_2_ = 1/*R*_2_ are the two principal curvatures (with *R*_1_, *R*_2_ the radii) at any point on the surface of the lipid assembly, which may be a bilayer for vesicles or a monolayer for micelles; *C*=*C*_1_+*C*_2_ is the mean curvature and *C*_1_*C*_2_ is the Gaussian curvature; κ and κ_G_ are the bending and Gaussian elastic moduli, respectively. Lipid molecular shape determines *C*_0_: lipids with similar areas of lipid headgroup and hydrophobic tails, i.e., a cylindrical shape, have a spontaneous curvature *C*_0_≈0. Cone-shaped lipids, with a headgroup area that is larger than that of the tails, have *C*_0_˃0, and inverted cone-shaped lipids, with headgroup area smaller than that of the tails, have *C*_0_˂0.

Lipids assemble in such a way to minimize the energetic costs described by equation (1) where the first term describes the elastic energy cost to bend the lipid assembly away from *C*_0_. Thus, *C*_0_ of a lipid formulation usually determines its structure (expressed as the curvature *C*). If their free energies are similar, the coexistence of structures with distinct curvatures may be observed.

The observation of discs in experiments, with lipid chains in either the solid or fluid state, has typically required at least a two-component system. Therefore, we need to consider the spontaneous curvature of a lipid mixture, *C*_0,mixture_. If lipid mixing is complete, *C*_0,mixture_ is the sum of the spontaneous curvatures of the individual lipids weighted by their mole fractions, *x*_i_: *C*_0,mixture_=Σ*x*_i_*C*_0,i_. Alternatively, lateral phase separation of lipids with differing *C*_0_ may occur, giving rise to structures with connected areas of different curvature such as disc micelles (Figure 1C).

As mentioned earlier, both cationic lipids and PEG-lipids are expected to have positive spontaneous curvature (*C*_0_˃0) with relatively large headgroup areas due to a combination of lateral electrostatic and headgroup steric repulsions. This is confirmed by the structures they form when mixed with cylindrically shaped (*C*_0_=0) DOPC. For example, lipid assemblies with increasing curvature form as the content of MVL5 in the membrane increases: primarily vesicles at low content, disc micelles (with high-curvature edges) at intermediate content, and worm-like (cylindrical) micelles and spheres at high content.

To estimate the spontaneous curvature of MVL5, PEG2K-lipid and PEG5K-lipid, we consider the relative circular areas of the headgroups compared to the tail area of the lipids. Figure 7 displays a schematic side view of the effective diameters of the lipid headgroup (*D*_h_) and tails (*D*_t_), the tail length (*l*_c_), and the vertical distance (*l*’) from the end of the tails to the center of the circle with spontaneous radius *R*_0_ = *l*_c_ + *l*’. Using the geometric relation (*D*_h_/2)/(*l*_c_+*l*’) = (*D*_t_/2)/*l*’, we obtain *l*’=*bl*_c_ with *b*=*D*_t_/(*D*_h_–*D*_t_). Thus, we get

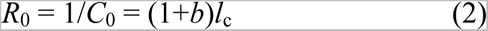

**Figure 7.**
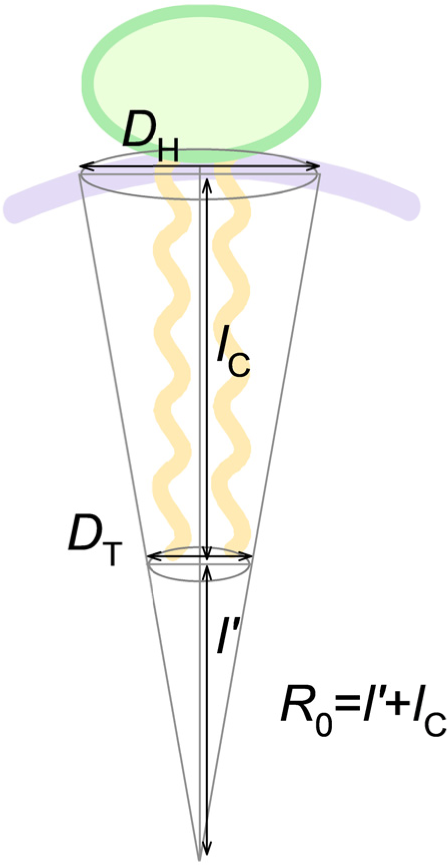
Schematic of a lipid with positive spontaneous curvature, labeled with the parameters used to estimate the radius of the spontaneous curvature from the lipid’s dimensions.

For cylindrically shaped DOPC with dioleoyl chains, *l*_c_ =19.5 Å,^70^ and the tail area π(*D*_t_/2)^2^ approximately equals the area per headgroup of ≈72 Å^2,58^ resulting in *D*_h_ = *D*_t_≈9.6 Å. For MVL5, the area per headgroup (π(*D*_h_(MVL5)/2)^2^) was previously measured to be ≈2.3 times larger than the area per headgroup of DOPC.^58^ Thus, we obtain *D*_h_(MVL5)=14.5 Å and *b*(MVL5)=1.94, resulting in the spontaneous curvature *C*_0_(MVL5) = 1/*R_0_*(MVL5) ≈1/(2.95*l*_c_)≈0.175 nm^−1^.

To obtain an approximation of *D*_h_ for the PEG2K-lipid, we estimate the relevant area per headgroup of the PEG2K-lipid as defined by the concentration just beyond the polymer chain overlap concentration in the brush regime. For PEG2K-lipids, this chain overlap and the resulting mushroom–brush transition occurs at ≈7 mol% in mixtures with DOPC.^71^ At this concentration, each PEG2K-lipid is mixed with ≈13.3 DOPC lipids on average, so that π(*D*_h_(PEG2K-lipid)/2)^2^ ≈ (1+13.3)π(*D*_t_/2)^2^ (i.e., each PEG chain attached to the PEG2K-lipid tails provides an umbrella covering ≈13.3 DOPC lipids). This results in *D*_h_(PEG2K-lipid) ≈ (1+13.3)^1/2^*D*_t_ ≈ 3.78*D*_t_ ≈36.2 Å ≈ 1.03*R*_G_(PEG2K), using an experimentally determined value (35 Å) for the radius of gyration of a PEG chain with molecular weight 2000 g/mol.^62,72,73^ With the corresponding value of *b*=0.36, we obtain the spontaneous curvature as *C*_0_(PEG2K-lipid) = 1/*R*_0_(PEG2K-lipid) ≈ 1/(1.36*l*_c_)≈0.377 nm^−1^.

For the PEG5K-lipid, we expect that *D*_h_(PEG5K) = (*R* ^5K^/*R* ^2K^)*D* (PEG2K). Using the experimentally determined value of *R*_G_(PEG5K) (62 Å),^62,72,73^ this gives *D*_h_(PEG5K-lipid) = 1.77*D*_h_(PEG2K-lipid) = 64.1 Å, resulting in *b*=0.175 and a spontaneous curvature *C*_0_(PEG5K-lipid) = 1/*R*_0_(PEG5K-lipid) ≈ 1/(1.175*l*_c_) ≈0.436 nm^−1^. (We note that the measured radius of gyration of PEG in water agrees well with the Flory equation for a polymer in a good solvent *R*_G_=3.6*n*^0.6^ Å where *n* is the degree of polymerization (*n*≈113 and 45 for the PEG5K- and PEG2K-lipid, respectively).^62,72,73^ Using this relation, we obtain *D*_h_(PEG5K-lipid) = 1.74*D*_h_(PEG2K-lipid) = 62.9 Å, corresponding to *b*=0.180, and a spontaneous curvature *C*_0_(PEG5K-lipid) = 1/*R*_0_(PEG5K-lipid) ≈ 1/(1.180*l*_c_) ≈0.435 nm^−1^.)

Consistent with our estimates, the cryo-TEM images of the samples with 10 mol% cone-shaped lipids show that the spontaneous curvature of the lipids increases as *C*_0_(MVL5) < *C*_0_(PEG2K-lipid) < *C*_0_(PEG5K-lipid) because the observed lipid assembly structures are vesicles for 10 mol% MVL5 and vesicles with increasing fraction of disc micelles going from 10 mol% PEG2K-lipid to 10 mol% PEG5K-lipid. In addition, the data confirms the expectation that *C*_0_(DOTAP) < *C*_0_(MVL5), based on the binary mixtures at 50 mol% cationic lipid as well as the ternary mixtures containing 10 mol% PEG2K-lipid.

The same order of spontaneous curvatures is also consistent with the trend toward structures with increasing average curvature (vesicles<discs<worms<spheres) as the content of cone-shaped lipid increases. Consider, for example, the structures observed in the formulations containing 10 mol% MVL5; 10 mol% MVL5 and 10 mol% PEG2K-lipid; 50 mol% MVL5; 50 mol% MVL5 and 10 mol% PEG2K-lipid; and 75 mol% MVL5 and 10 mol% PEG2K-lipid. They are vesicles, vesicles and discs, discs only, and worms and spheres with an increase in the abundance of spheres, respectively (see Figures 3 and 5, as well as Figures S2–S7 and S24–S30). This is consistent with the theoretical expectation that *C*_0,mixture_=Σ*x*_i_*C*_0,i_.

However, our experimental findings make it clear that there are additional, lipid(-type)-specific parameters to be considered. For example, the ternary formulations containing univalent lipid show very small vesicles, very small discs, and spheres in addition to large vesicles, much like the respective binary formulations containing only cationic lipid. The main effect of the additional 10 or even 25 mol% PEG2K-lipid is the elimination of large and very large vesicles and an increase in the size of the discs, which are larger in binary formulations containing PEG2K-lipid than in those with charged lipids. In other words, these structures are more distinct from the binary formulations containing PEG2K-lipid than from those containing DOTAP, despite the very high *C*_0_ of the PEG-lipids. The data thus suggest that the charge of the lipid assembly itself is a factor promoting the formation of very small assemblies (both vesicles and micelles). The likely explanation for this effect is that the electrostatic self-energy of charged closed objects, such as micellar discs or rods, drives a decrease of the size of the lipid assembly which, in turn, is resisted by the enhanced costs in curvature energy for the decreased particle size. In addition, the data suggests that PEG-lipid is more effective at preventing fusion of small assemblies into larger vesicles than charge.

The most striking finding in this study is the essentially exclusive formation of disc micelles in the formulation containing MVL5. The multivalent headgroup allows access to charge densities of the membrane well beyond what may be achieved with univalent lipids. As the imaging shows, this gives rise to phases (such as the pure phase of discs) and structures (such as the worm-like micelles) unobtainable with univalent lipids, even when the univalent lipids are combined with PEG-lipids. A possible explanation for the preference of MVL5 for discs is that the high spontaneous curvature of MVL5 strongly disfavors its presence in the inner monolayer of vesicles, especially very small ones. Furthermore, in binary mixtures with DOPC, phase separation would also favor discs with MVL5 lowering the elastic cost of high curvature edges. This argument is also valid for the PEG-lipids with their high spontaneous curvature, though the long-range electrostatic repulsion of the MVL5 headgroups may result in contributions beyond next neighbors.

Control of the size of lipid discs is of great interest for possible applications. Summarizing the trends observed in this work, it appears that both increasing *C*_0_ and increasing charge decreases the size of fluid disc micelles. For example, compare the average size of the discs formed at 25 mol% PEG2K-lipid and PEG5K-lipid (Figure 4), or the drastic effect of including DOTAP on the size of the micelles at both 10 and 25 mol% PEG2K-lipid (Figures 6 and S32– S38). Exploiting these trends, possibly by using novel lipids with specifically designed headgroups, promises increased control over the important parameter of disc size.

A relatively recent single-component model^65^ based on membrane curvature elasticity (Eq. (1)) predicts a transition between bilayer structures (such as vesicles) and monolayer micelles (discs or rods) as a function of *C*_0_ and the tail length *l*_c_. The model predicts that bilayers are favored when *l*_c_×*C*_0_ < ¼, while *l*_c_×*C*_0_ > ¼ favors monolayer micelles.^65^ Furthermore, in this model, discs are preferred for large membrane bending modulus κ (compared to k_B_*T*), while rods are favored at small κ. This could be a possible explanation for the strong shift toward flexible worm-like and spherical micelles that incorporation of PEG-lipids has on formulations with higher content of MVL5.

Using our estimates for the spontaneous curvatures of MVL5 and the PEG-lipids, we can compare the predictions of the model^65^ to the data regarding transitions from bilayer membranes (vesicles) to micellar structures. For PEG2K-lipid mixed with DOPC, a phase of coexisting small bilayer vesicles and disc micelles is observed in cryo-TEM at 10 mol% PEG2K-lipid (Figure 2). For membranes with uniformly mixed lipid components, this would result in an average spontaneous curvature *C*_0_(10% PEG2K-lipid) = 0.1*C*_0_(PEG2K-lipid) ≈ 1/13.6*l*_c_, which is much less than the spontaneous curvature of 1/4*l*_c_ predicted by the model as the condition for micelle formation. This curvature would be achieved by a uniformly mixed formulation with a molar fraction of PEG2K-lipid of ≈0.34. For MVL5, a phase of nearly pure disc micelles is seen in cryo-TEM images in two-component membranes consisting of 50 mol% of MVL5 with DOPC (Figure 3). Here, we would estimate the average spontaneous curvature of a uniformly mixed phase *C*_0_(50%MVL5) = 0.5*C*_0_(MVL5) ≈ 1/5.9*l*_c_, which again is <1/4*l*_c_. A *C*_0_ of 1/4*l*_c_ would only be reached at a MVL5 molar fraction of 0.74, which is close to the composition (75 mol%) where cryo-EM showed spherical and worm-like micelles in our experiments (Figure 3). Finally, for the formulation containing 25 mol% PEG5K-lipid, which largely consisted of disc micelles (Figure 4), *C*_0_(25% PEG5K-lipid) = 0.25*C*_0_(PEG5K-lipid) ≈ 1/(4 × 1.175)*l*_c_ ≈0.21/*l*_c_, very close to the proposed condition^65^ for micelle formation. For this lipid, application of the model predicts micelle formation at a molar fraction of 0.29.

With the above comparisons, it is important to remember that the model assumes a single component system. Our formulations, in contrast, consist of at least two lipid components (as do other known disc-forming formulations). This means that lipid–lipid phase separation may occur, segregating high-curvature lipids into the curved edges of discs (Figure 1) where their local content could be high enough to satisfy the condition for micelle formation.

Sonication is required to form the smaller and micellar structures we observed for at least some of the formulations investigated. Without sonication, only vesicles of a variety of sizes (from 17 nm to > 1 μm in diameter, most of them large) are observed at 50 mol% DOTAP (see Figures S39, S40). When the formulation contains an additional 10 mol% PEG2K-lipid, the observed structures also include spherical micelles and occasional discs (24–41 nm diameter) (see Figures S41, S42).

## Conclusion

Lipid assemblies with fluid membranes are promising carriers of hydrophobic drugs. We have shown that the inclusion of cone-shaped lipids (*C*_0_ ˃ 0) – charged lipids as well as PEG-lipids – in fluid-phase membranes with DOPC effects structural phase transitions from spherical vesicles to mixtures of vesicles and micelles with anisotropic (disc, worm-like rods of varying length) and spherical shapes. In most formulations, different assembly structures coexist. Remarkably, however, higher contents of the multivalent lipid MVL5 form pure and mixed micellar phases with minimal vesicles. With a size of order ≈10–30 nm, these fluid lipid nanodiscs are significantly smaller than liposomal carriers of hydrophobic drugs (which typically range from ≈50–150 nm in size). The observed trends regarding formation and size of the nanodiscs will inform ongoing efforts to develop novel lipid-based delivery systems for small hydrophobic drugs.

Continuing work will focus on identifying additional lipid compositions that exhibit pure phases to build a library of micellar building blocks. Observations about structural phase transitions as a function of lipid charge and shape from this and other studies will also direct the design of new synthetic lipids that promote stable lipid micelles. For example, synthesis of charged PEG-lipids for use in place of separate charged lipid and PEG-lipid components might eliminate the phase separation observed for the PEGylated lipid assemblies containing MVL5, where spherical and cylindrical micelles coexist, and thereby promote the formation of pure micellar phases. With regard to applications, it is also important to note that all cryo-TEM samples were prepared in deionized water. Depending on the application, lipid assemblies are likely to be used in solutions with salt. Salt ions screen electrostatic interactions between charged particles, which decreases the lateral repulsion between charged headgroups such as those of DOTAP and MVL5. Thus, these cationic lipids will have a reduced *C*_0_ in salt solution. The *C*_0_ of PEG-lipids should be less affected by ionic strength. Therefore, in salt-containing solution, a shift toward the structures observed for assemblies without charged lipids would be expected for the PEGylated assemblies containing charged lipids. Future work will thus investigate how the structures of the lipid assemblies, especially of those containing charged lipids, change in the presence of salt.

We expect that our experimental findings will stimulate new curvature elasticity models of lipid assemblies to more accurately predict transitions from bilayer vesicles to anisotropic (disc- or rod-shaped) micelles. In particular, for a direct comparison to our experimental system, future models will need to take into account lipid charge and consider membranes consisting of multiple components, thus allowing for phase separation within the membrane. In addition, it is possible that multivalent charged lipids, which challenge the validity of mean-field theoretical approaches, will require novel theories.

## Materials and Methods

### Materials

Lipid stock solutions of DOPC, DOTAP, and 1,2-dioleoyl-sn-glycero-3-phospho-ethanolamine-N-[methoxy(polyethylene glycol)-5000] (ammonium salt) (PEG5K-lipid) in chloroform were purchased from Avanti Polar Lipids. PTX was purchased from Acros Organics and dissolved in chloroform at 10.0 mM concentration. MVL5^52,77,78^ and 1-(3,4-dioleyloxybenzoyl)-ω-methoxy-poly(ethylene glycol) 2000 (PEG2K-lipid)^78^ were synthesized as previously reported and dissolved in chloroform to a concentration of 10 mM.

### Liposome preparation

Solutions of lipid and PTX were prepared in chloroform:methanol (3:1, v/v) in small glass vials at a total molar concentration of 20 mM. Individual stock solutions of each component were combined according to the final desired concentration and molar composition. The organic solvent was evaporated by a stream of nitrogen for 10 min and dried further in a vacuum (rotary vane pump) for 16 h. The resulting films were hydrated with high-resistivity water (18.2 MΩ cm) to yield a 20 mM aqueous solution. Immediately thereafter, suspensions were agitated with a tip sonicator (Sonics and Materials Inc. Vibra Cell, set to 30 Watt output) for 7 min to form sonicated liposomes.

### Cryogenic Transmission Electron Microscopy

All samples were vitrified using a manual plunger on carbon lacey substrates (300 mesh copper grids) prepared in house.^79^ Grids were plasma cleaned using O_2_ and H_2_ for 30 s using a Solarus plasma cleaner (Gatan) immediately prior to sample preparation. A total of 3 μL of sample was applied to the grid and manually blotted from the back with filter paper for 5 s, followed immediately by plunging into liquid ethane. Images were acquired using Leginon^80^ on a Tecnai TF20 or Tecnai T12 equipped with 4K TVIPS CMOS camera, operated at 200 KeV or 120 KeV, respectively. Images were collected at nominal magnifications of 62k× (TF20) and 68k× (T12), corresponding to pixel sizes of 3.0 Å/pixel and 2.46 Å/pixel, respectively. A summary of the sample compositions (mol%) with image pixel size is provided in Table S1 in the Supporting Information.

## Acknowledgments

This research was supported by the National Institutes of Health under award R01GM130769 (mechanistic studies on developing lipid nanoparticles for drug delivery). Partial support was provided by the National Science Foundation under award DMR-1807327 (membrane phase behavior of lipid nanoparticles). Cryo-TEM experiments were conducted at the Simons Electron Microscopy Center and National Resource for Automated Molecular Microscopy located at the New York Structural Biology Center, supported by grants from the Simons Foundation (SF349247) and the NIH National Institute of General Medical Sciences (GM103310). The authors acknowledge useful discussions of the Cryo-TEM data with Bridget Carragher and Clinton Potter. V.M.S. was supported by the National Science Foundation Graduate Research Fellowship Program under Grant No. DGE 1144085. C.R.S. acknowledges sabbatical leave for Spring quarters 2022 and 2023, which allowed for scholarly activities related to the research reported here. He is also grateful for discussions with Robijn Bruinsma on structural transitions from bilayer vesicles to monolayer micelles.

