## Supporting Information for "The Influence of Multivalent Charge and PEGylation on Shape Transitions in Fluid Lipid Assemblies: From Vesicles to Discs, Rods, and Spheres"

for

### Contents

Chemical Structures of Lipids Used S2

Unlabeled and additional cryogenic TEM images S3–S43

Table of Sample Compositions S44

### Lipid Structures

DOPC

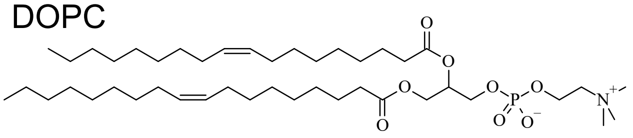

DOTAP

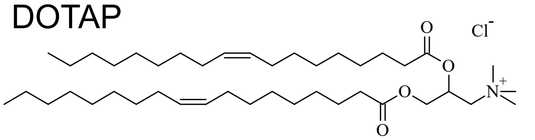

MVL5

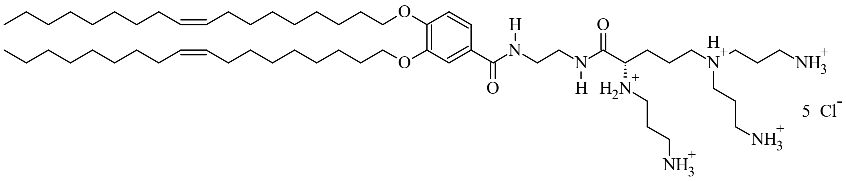

PEG2K-lipid

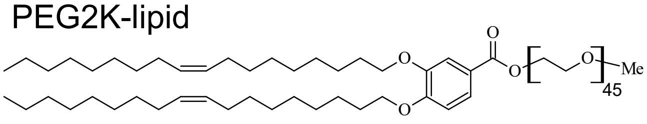

PEG5K-lipid

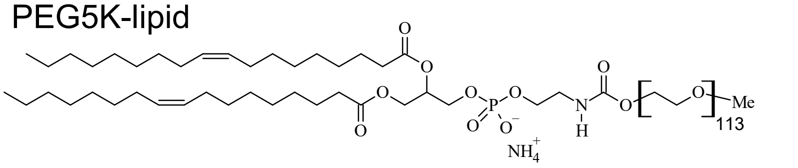

**Figure S1.** Chemical structures of the lipids used in this study.

### 10 mol% MVL5

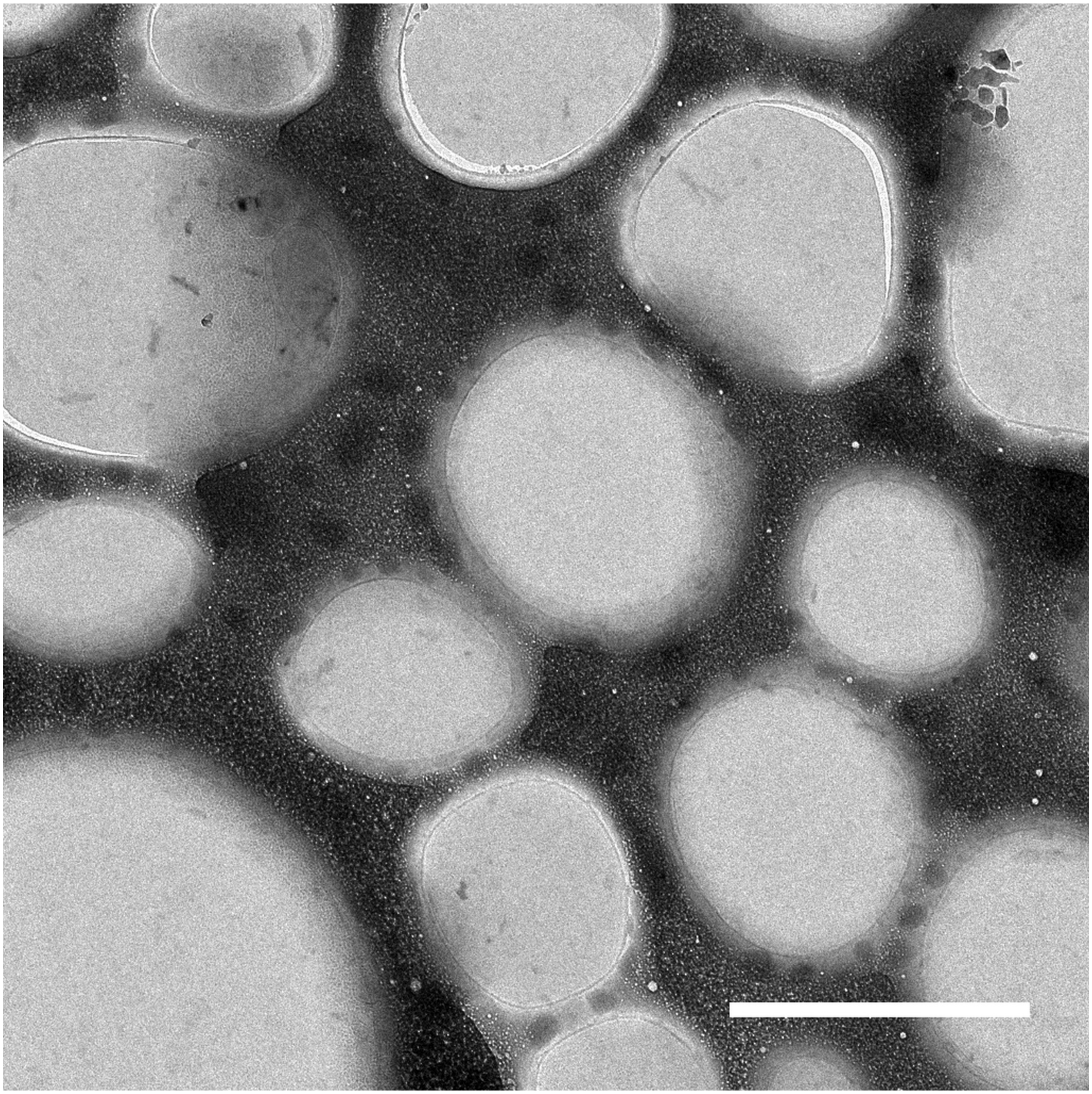

**Figure S2.** Lower magnification cryo-EM image of a sample containing 10 mol% MVL5, 3 mol% PTX, and the remainder DOPC. Scale bar: 2 μm.

### 10 mol% MVL5

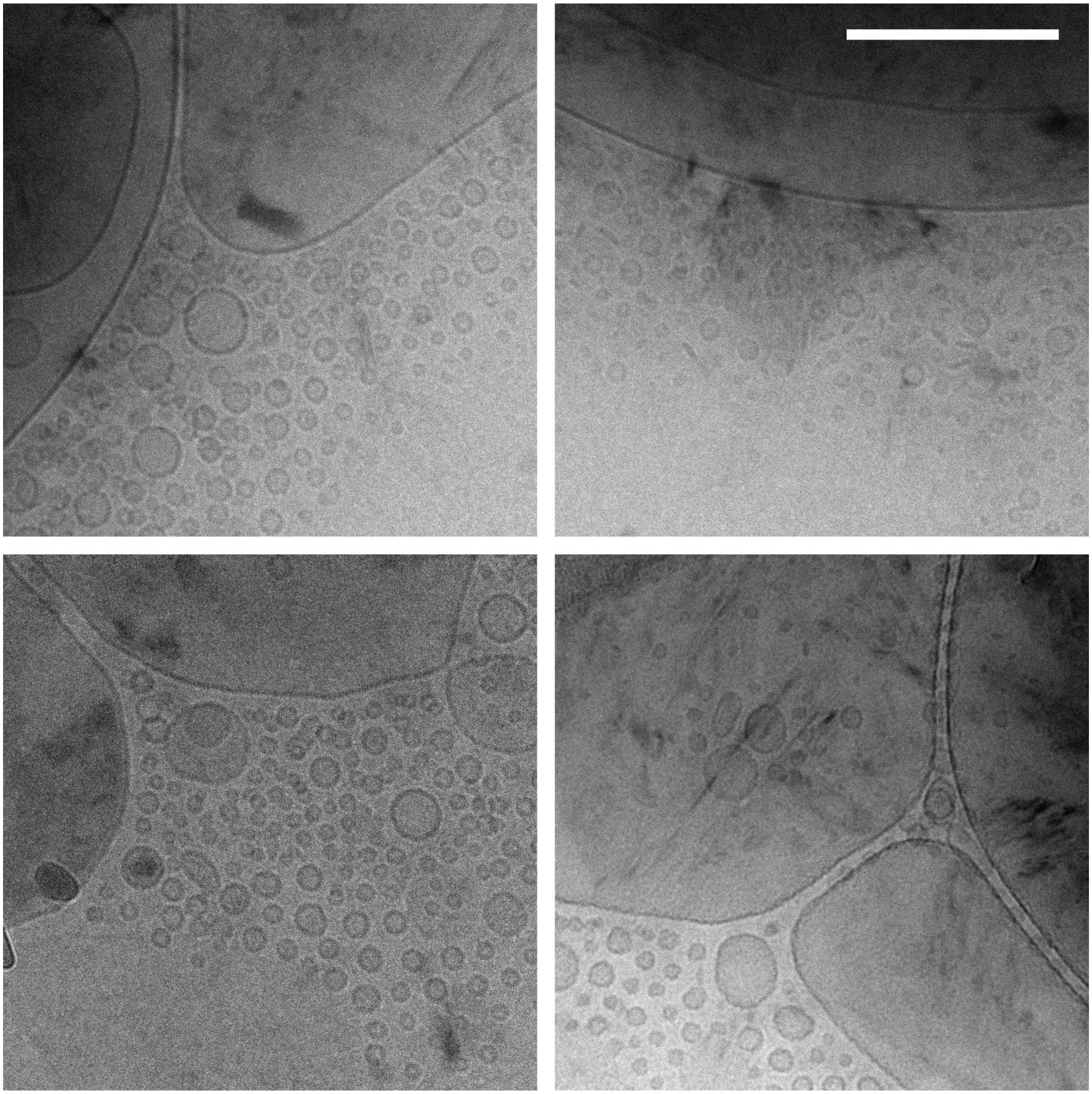

**Figure S3.** Cryo-EM images of a sample containing 10 mol% MVL5, 3 mol% PTX, and the remainder DOPC. Scale bar: 200 nm.

### 50 mol% MVL5

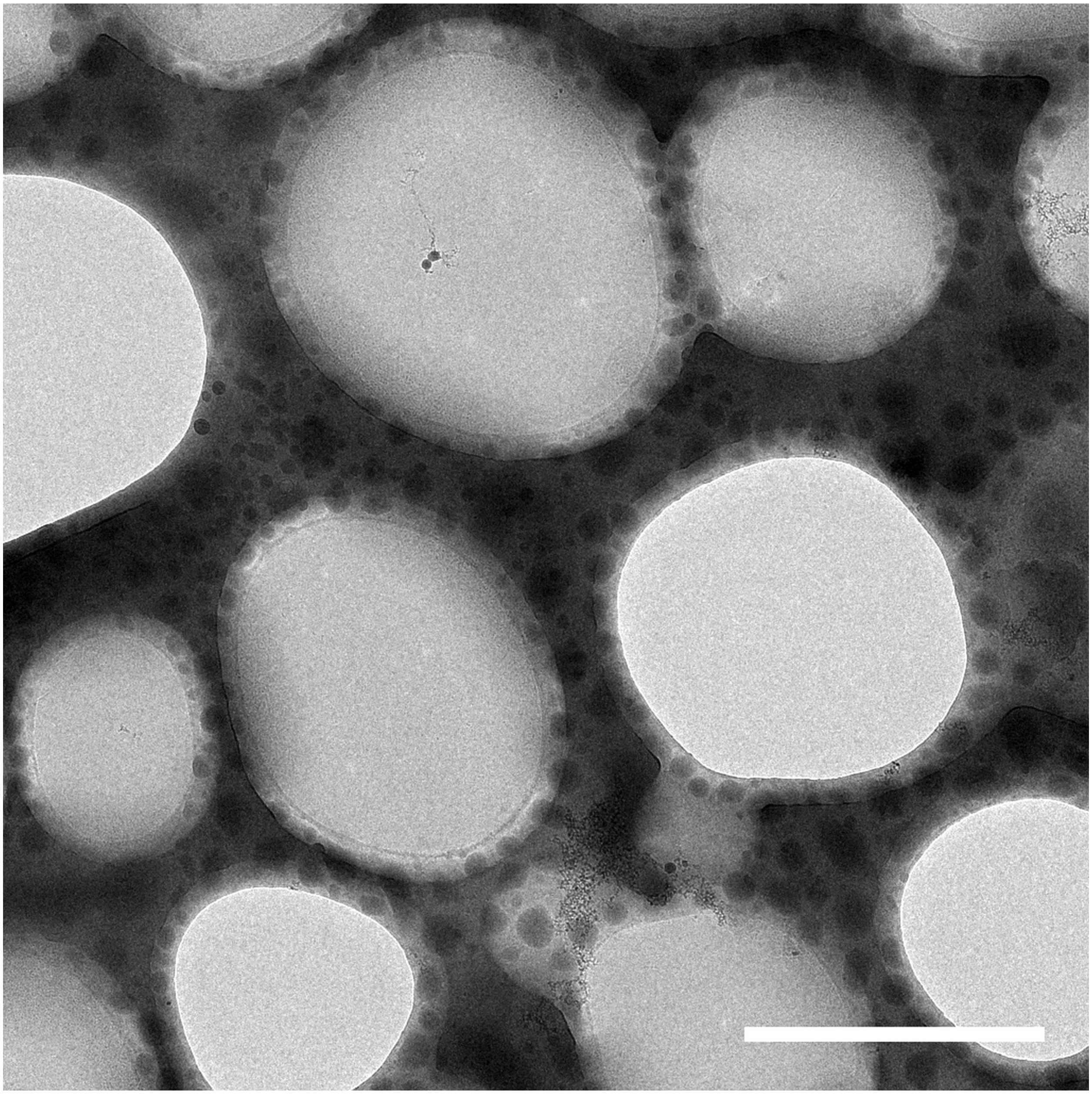

**Figure S4.** Lower magnification cryo-EM image of a sample containing 50 mol% MVL5, 3 mol% PTX, and the remainder DOPC. Scale bar: 2 μm.

### 50 mol% MVL5

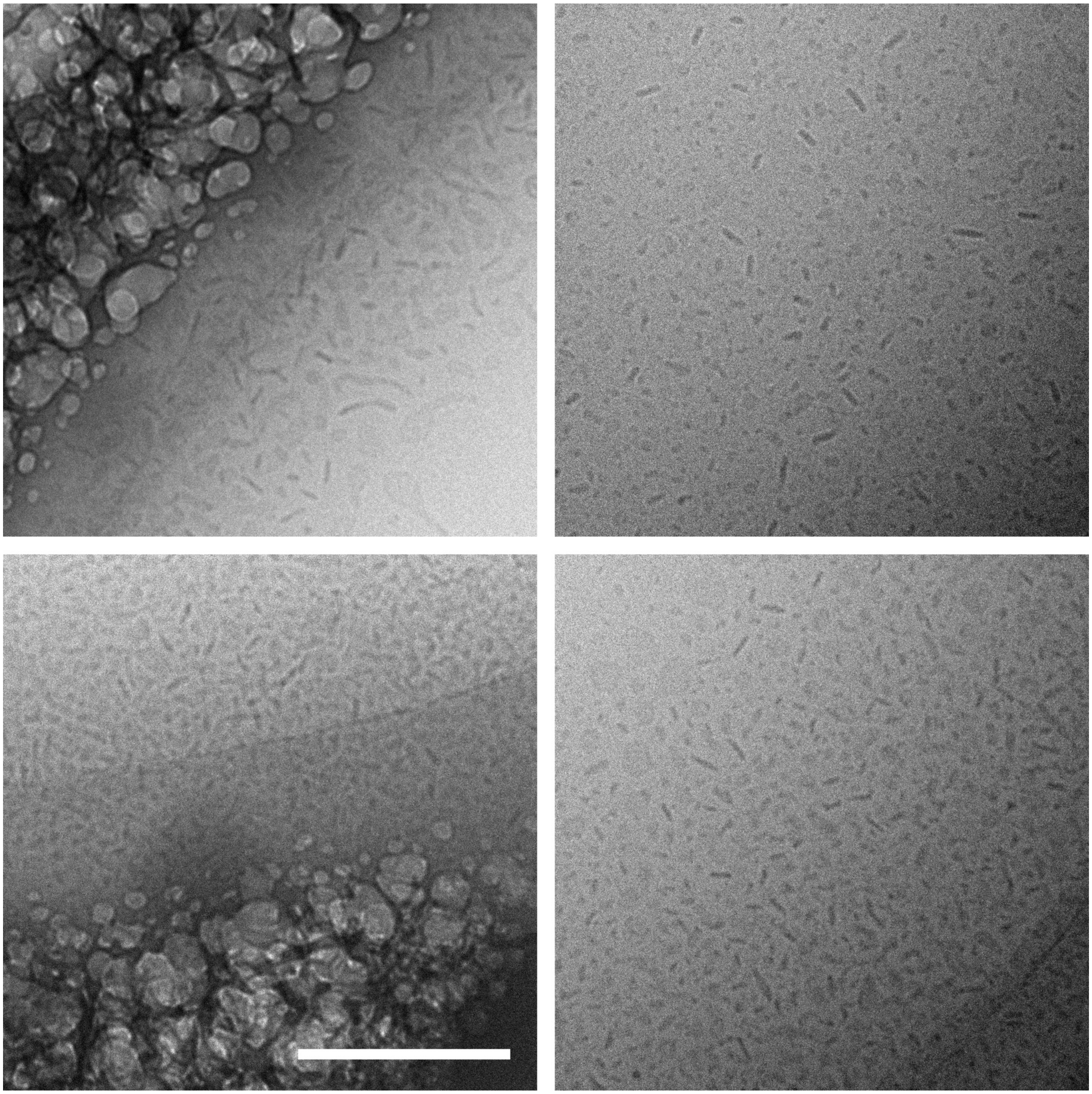

**Figure S5.** Cryo-EM images of a sample containing 50 mol% MVL5, 3 mol% PTX, and the remainder DOPC. Scale bar: 200 nm.

### 75 mol% MVL5

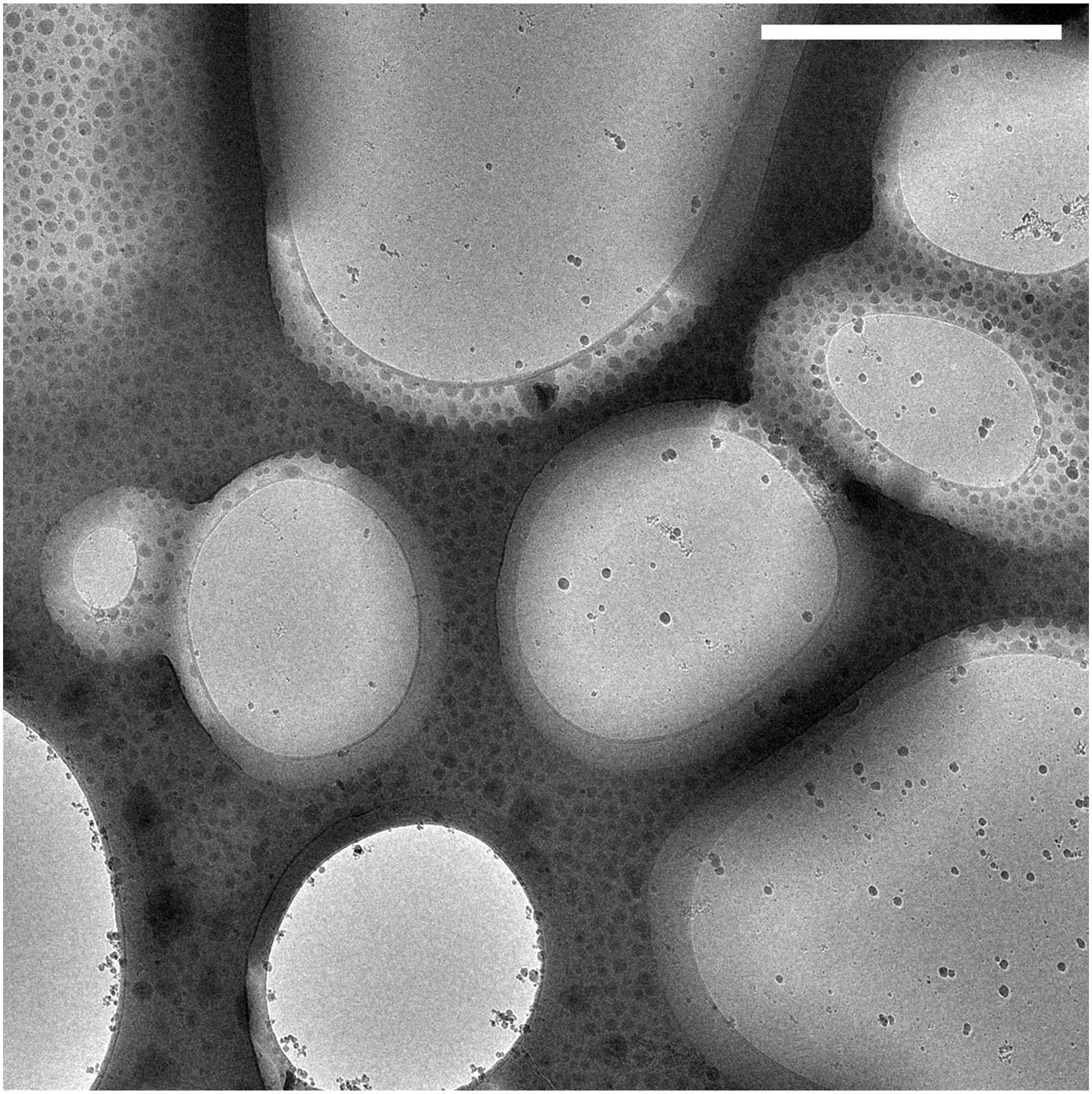

**Figure S6.** Lower magnification cryo-EM image of a sample containing 75 mol% MVL5, 3 mol% PTX, and the remainder DOPC. Scale bar: 2 μm.

### 75 mol% MVL5

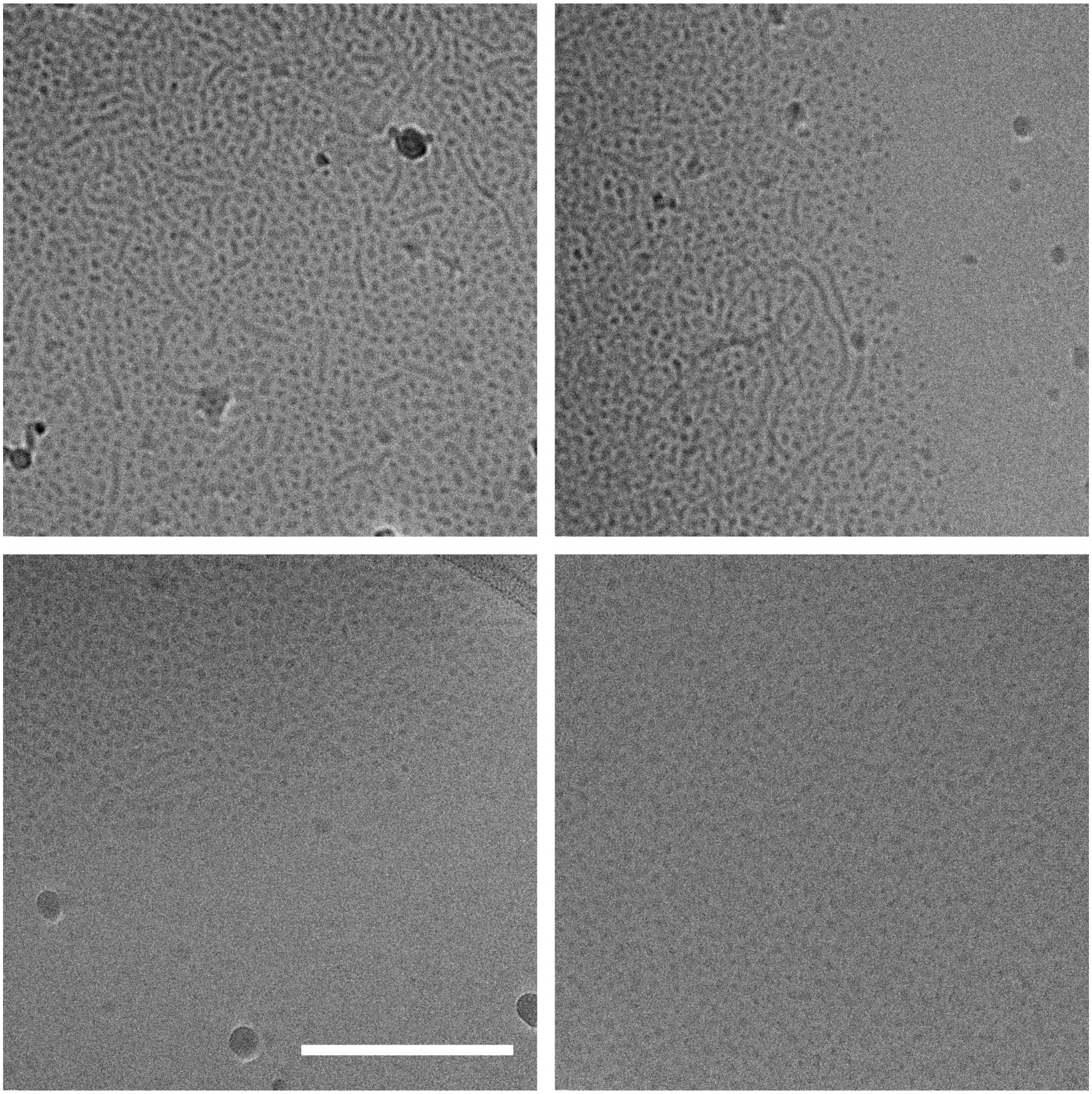

**Figure S7.** Cryo-EM images of a sample containing 75 mol% MVL5, 3 mol% PTX, and the remainder DOPC. Scale bar: 200 nm.

### DOPC only

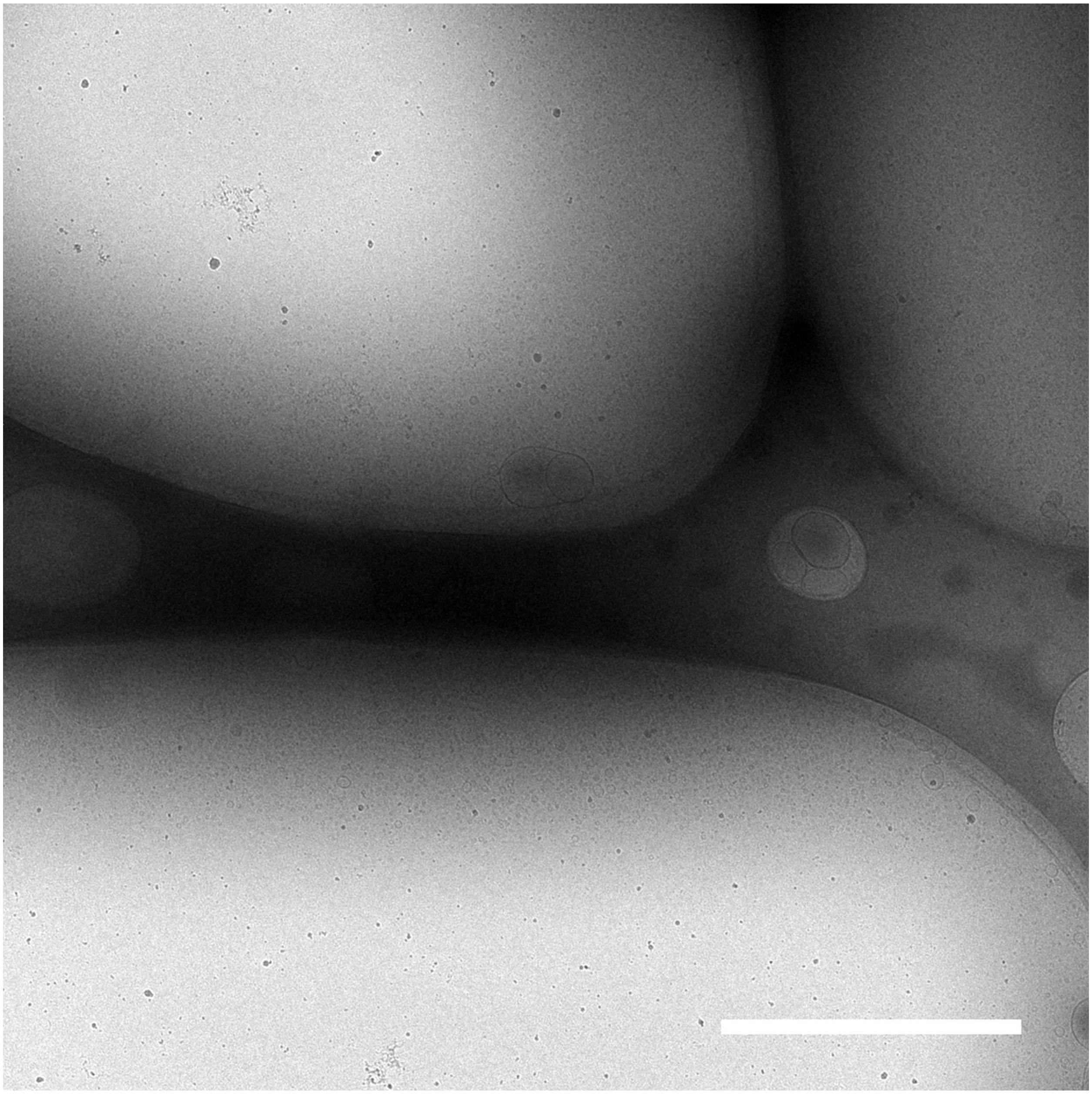

**Figure S8.** Lower magnification cryo-EM image of a sample containing 97 mol% DOPC,and 3 mol% PTX. Scale bar: 2 μm.

### DOPC only

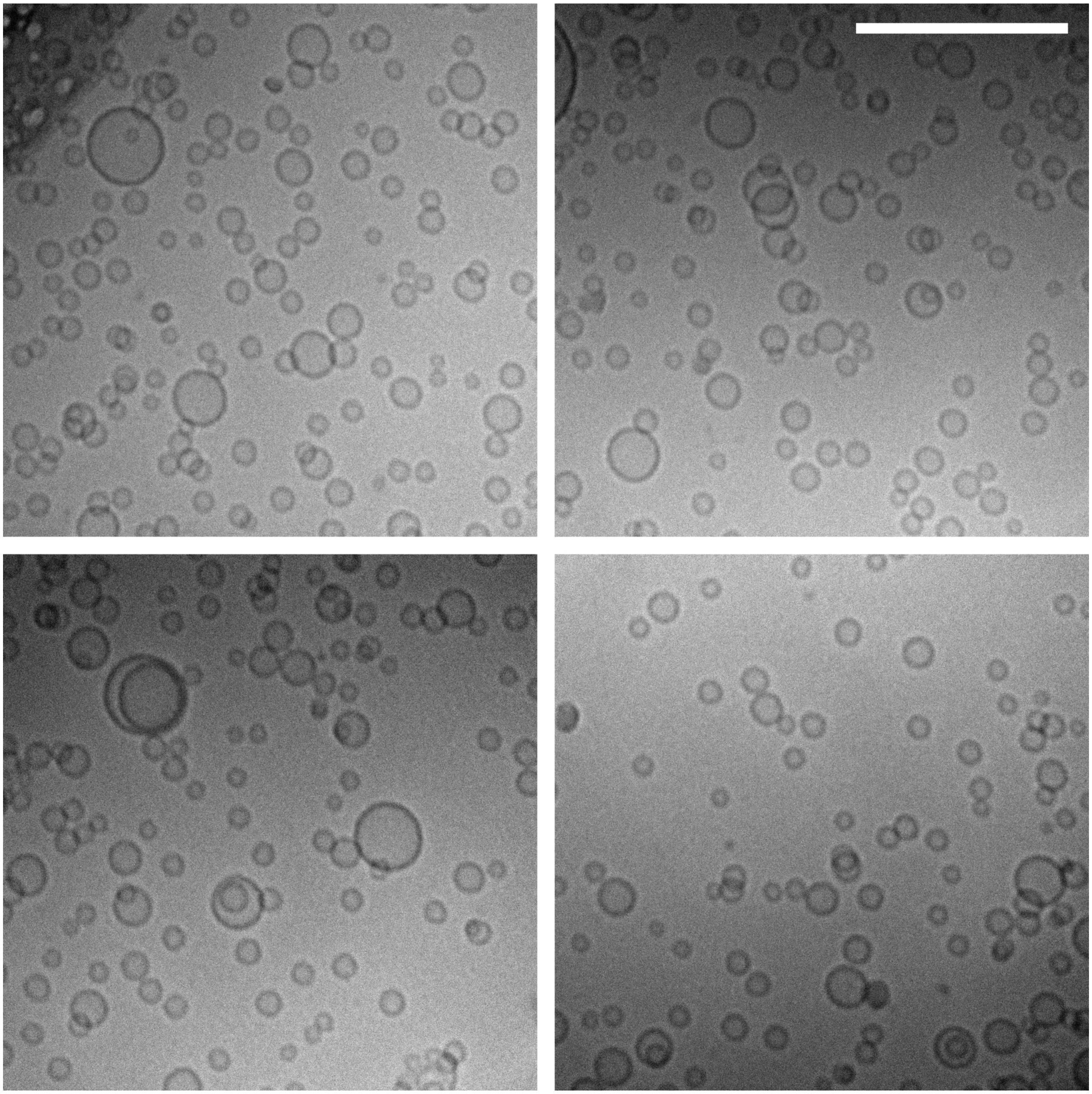

**Figure S9.** Cryo-EM images of a sample containing 97 mol% DOPC,and 3 mol% PTX. Scale bar: 200 nm.

### 2 mol% PEG2K-lipid

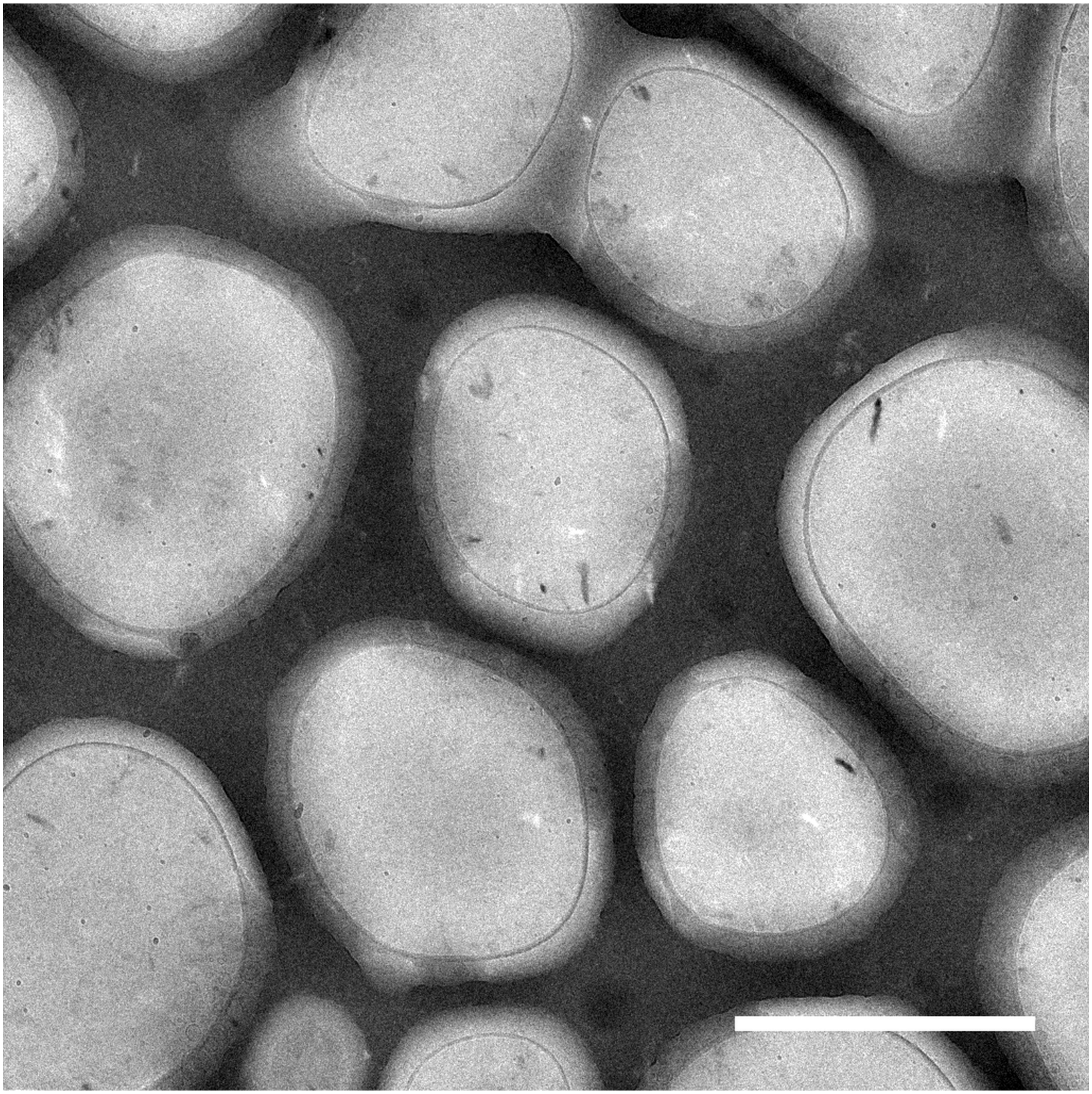

**Figure S10.** Lower magnification cryo-EM image of a sample containing 2 mol% PEG2K-lipid, 3 mol% PTX, and the remainder DOPC. Scale bar: 2 μm.

### 2 mol% PEG2K-lipid

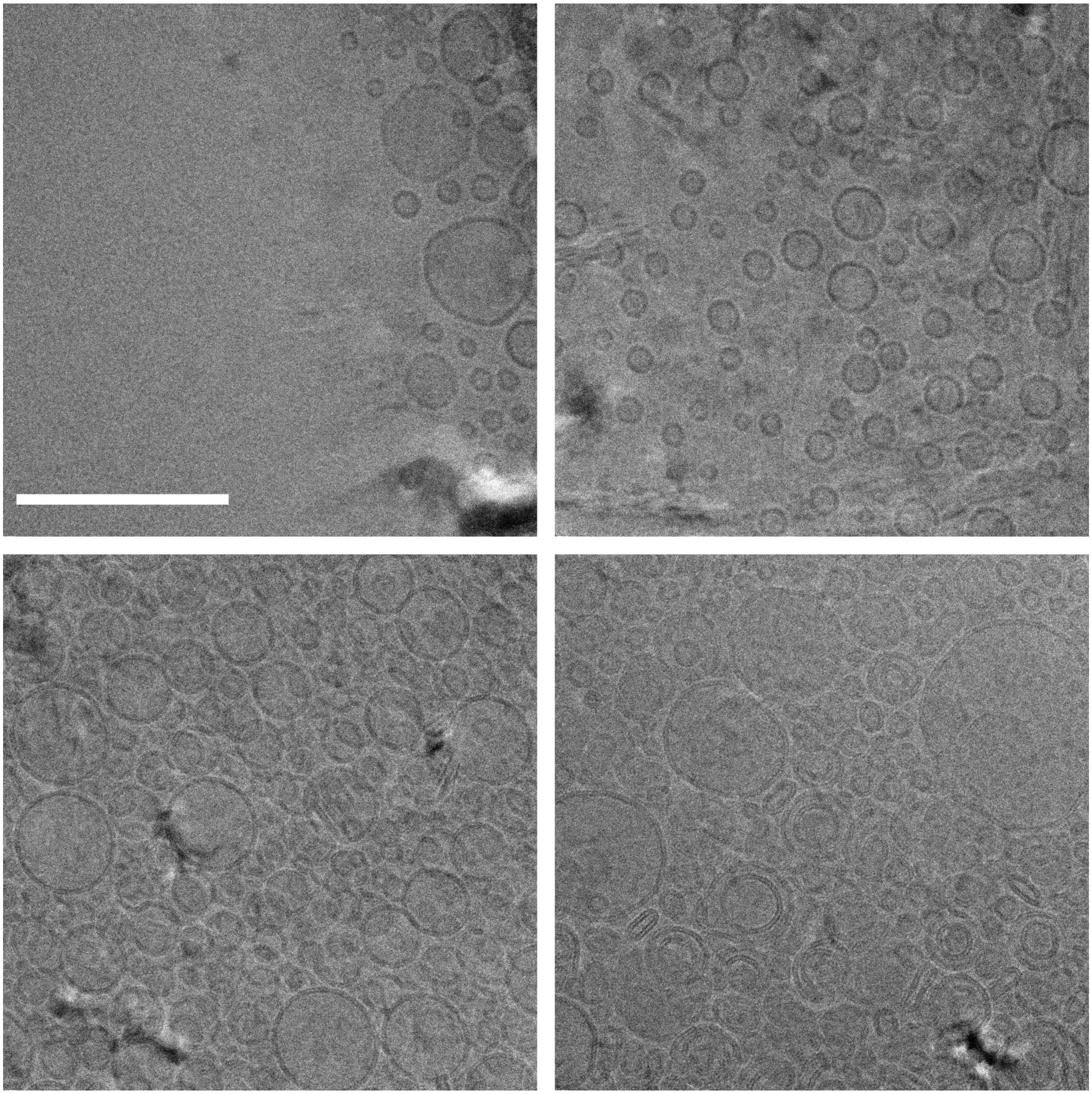

**Figure S11.** Cryo-EM images of a sample containing 2 mol% PEG2K-lipid, 3 mol% PTX, and the remainder DOPC. Scale bar: 200 nm.

### 10 mol% PEG2K-lipid

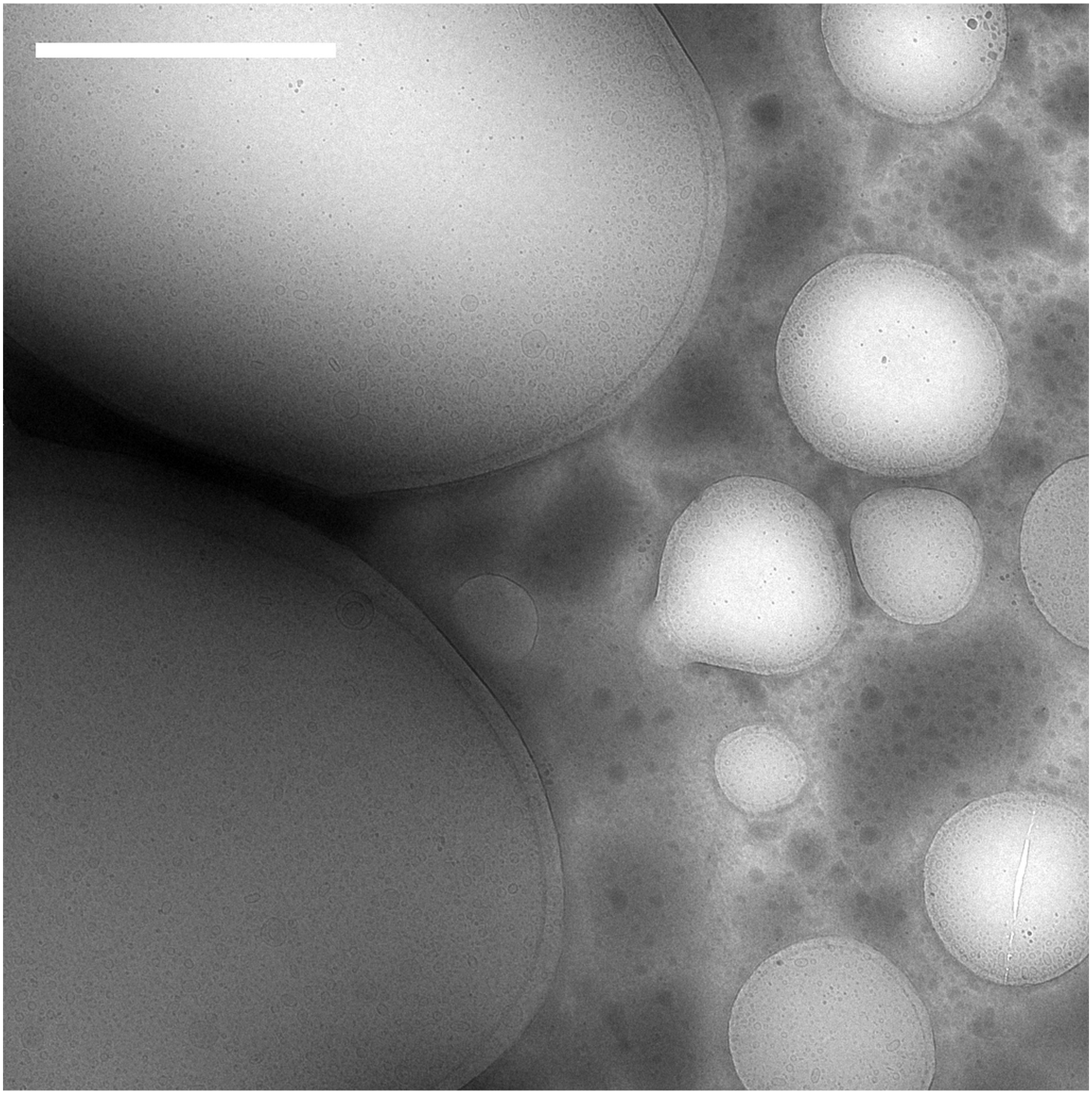

**Figure S12.** Lower magnification cryo-EM image of a sample containing 10 mol% PEG2K-lipid, 3 mol% PTX, and the remainder DOPC. Scale bar: 2 μm.

### 10 mol% PEG2K-lipid

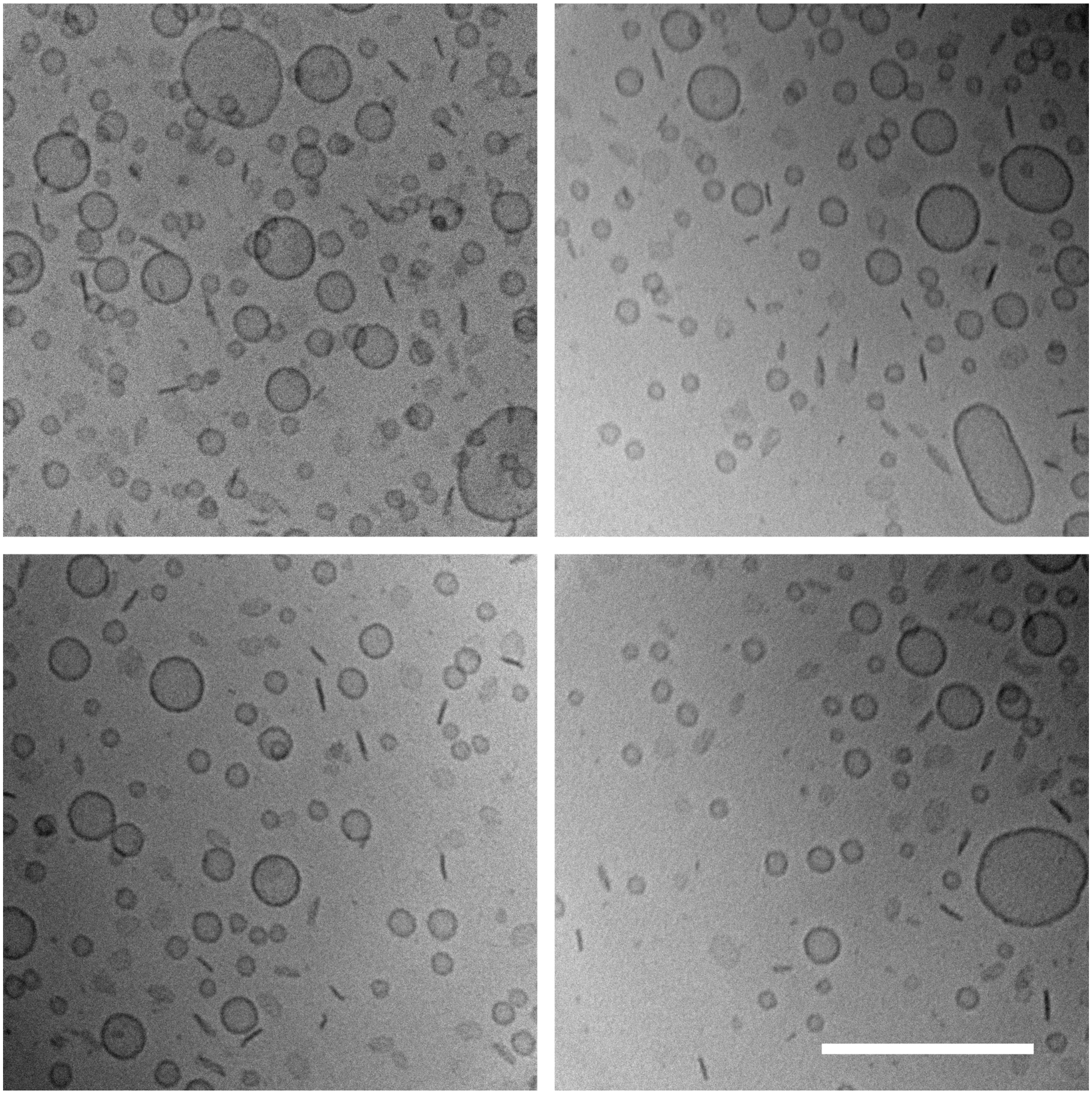

**Figure S13.** Cryo-EM images of a sample containing 10 mol% PEG2K-lipid, 3 mol% PTX, and the remainder DOPC. Scale bar: 200 nm.

### 50 mol% DOTAP

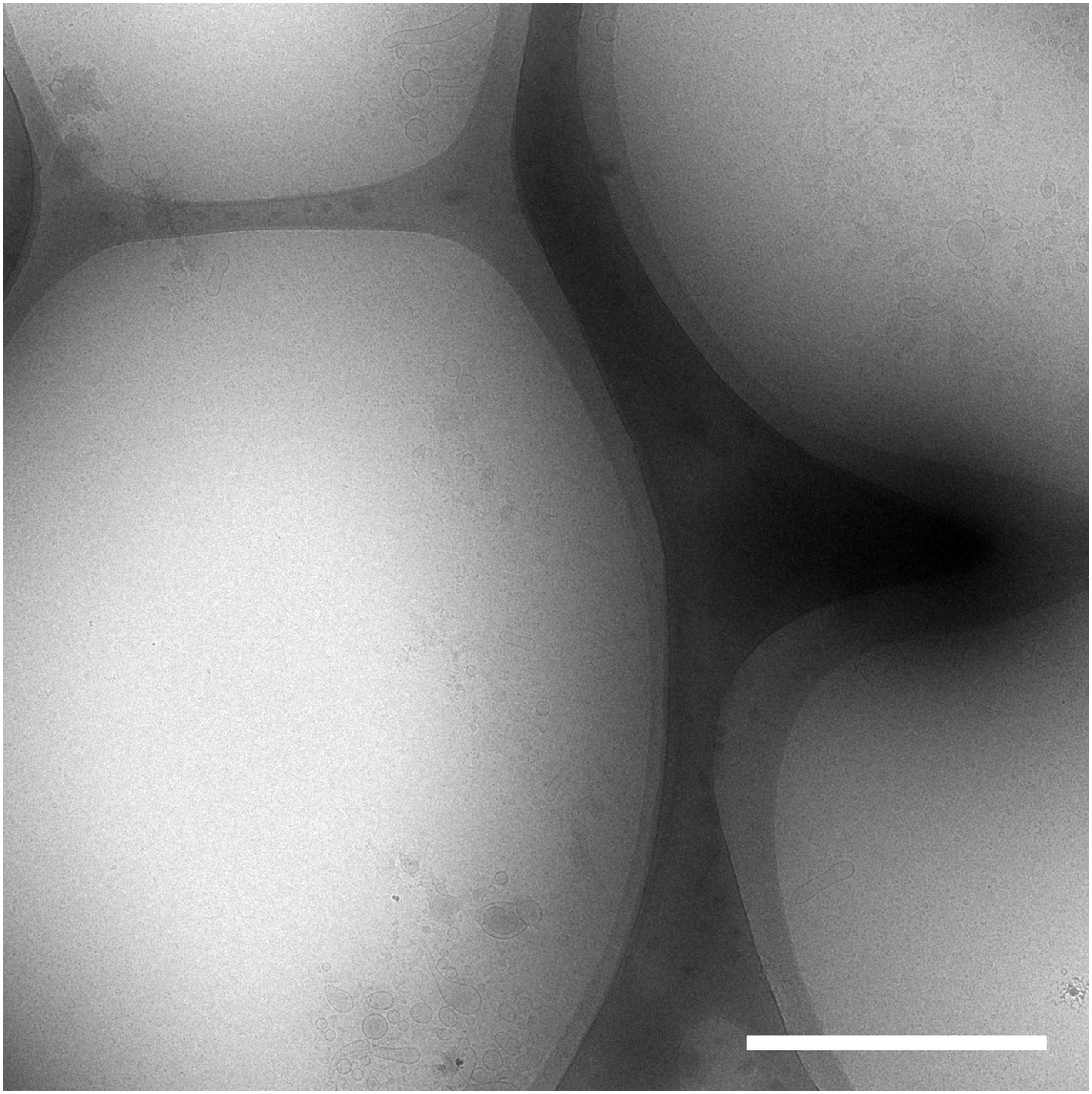

**Figure S14.** Lower magnification cryo-EM image of a sample containing 50 mol% DOTAP, 3 mol% PTX, and the remainder DOPC. Scale bar: 2 μm.

### 50 mol% DOTAP

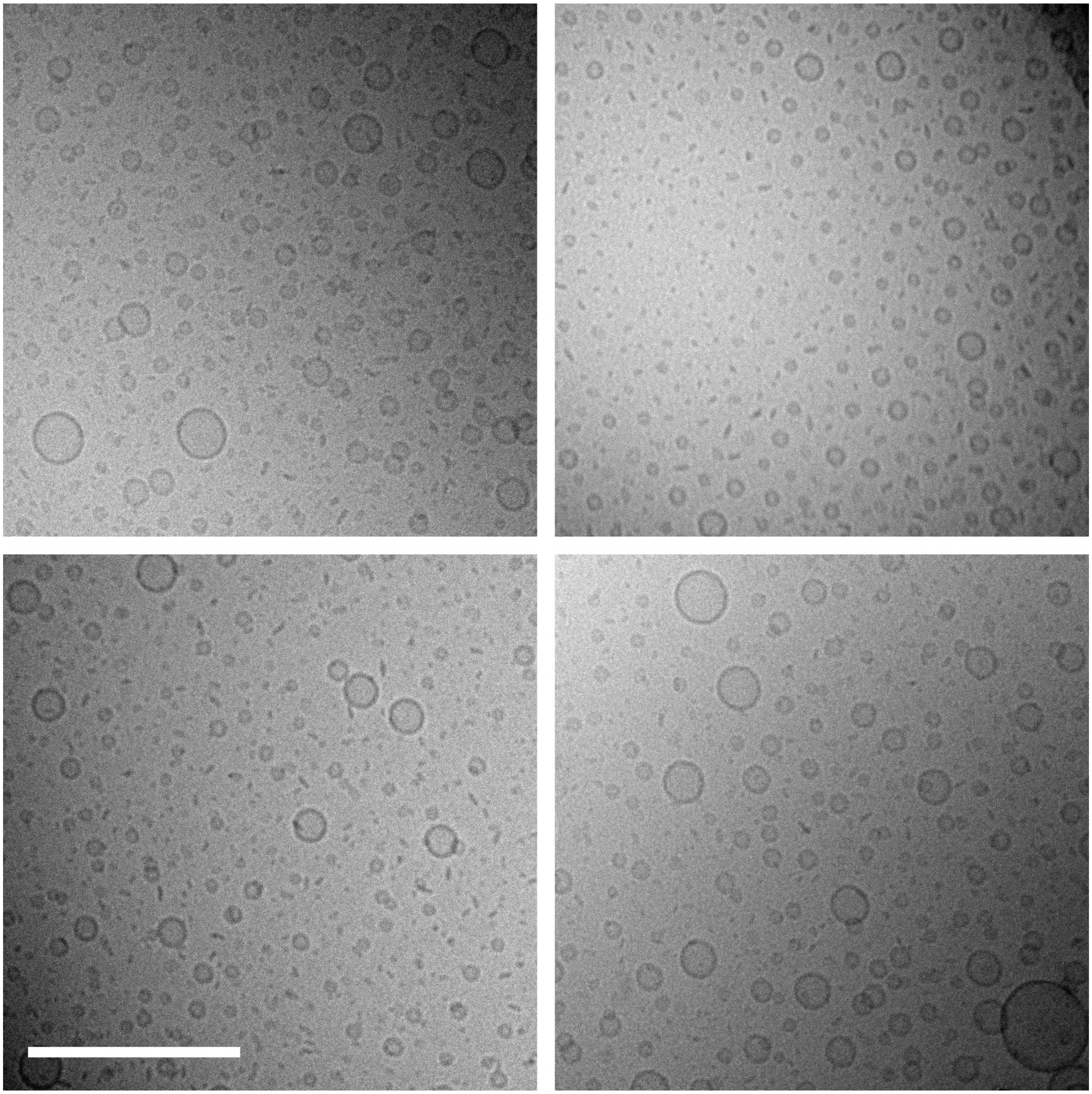

**Figure S15.** Cryo-EM images of a sample containing 50 mol% DOTAP, 3 mol% PTX, and the remainder DOPC. Scale bar: 200 nm.

### 80 mol% DOTAP

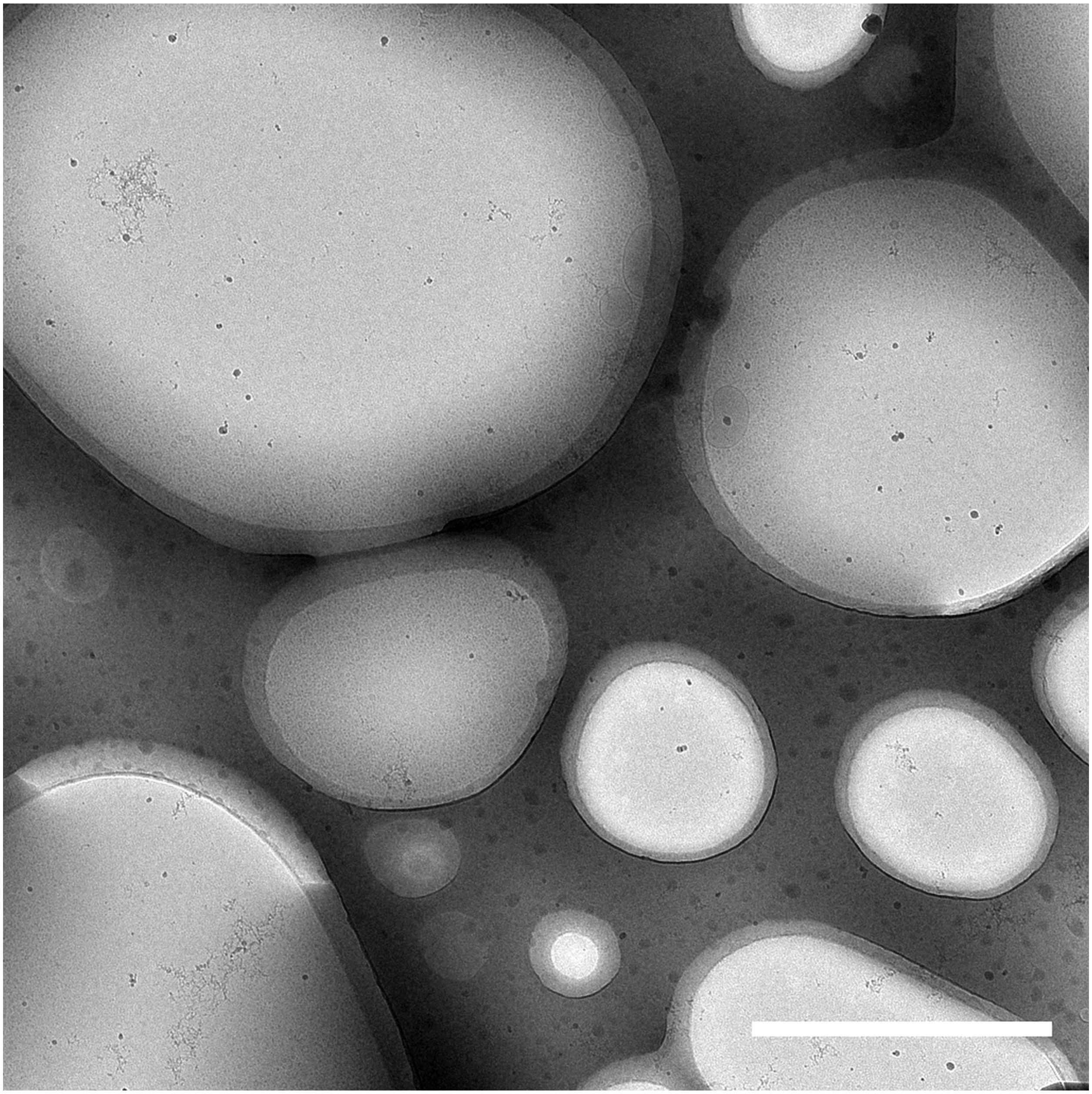

**Figure S16.** Lower magnification cryo-EM image of a sample containing 80 mol% DOTAP, 3 mol% PTX, and the remainder DOPC. Scale bar: 2 μm.

### 80 mol% DOTAP

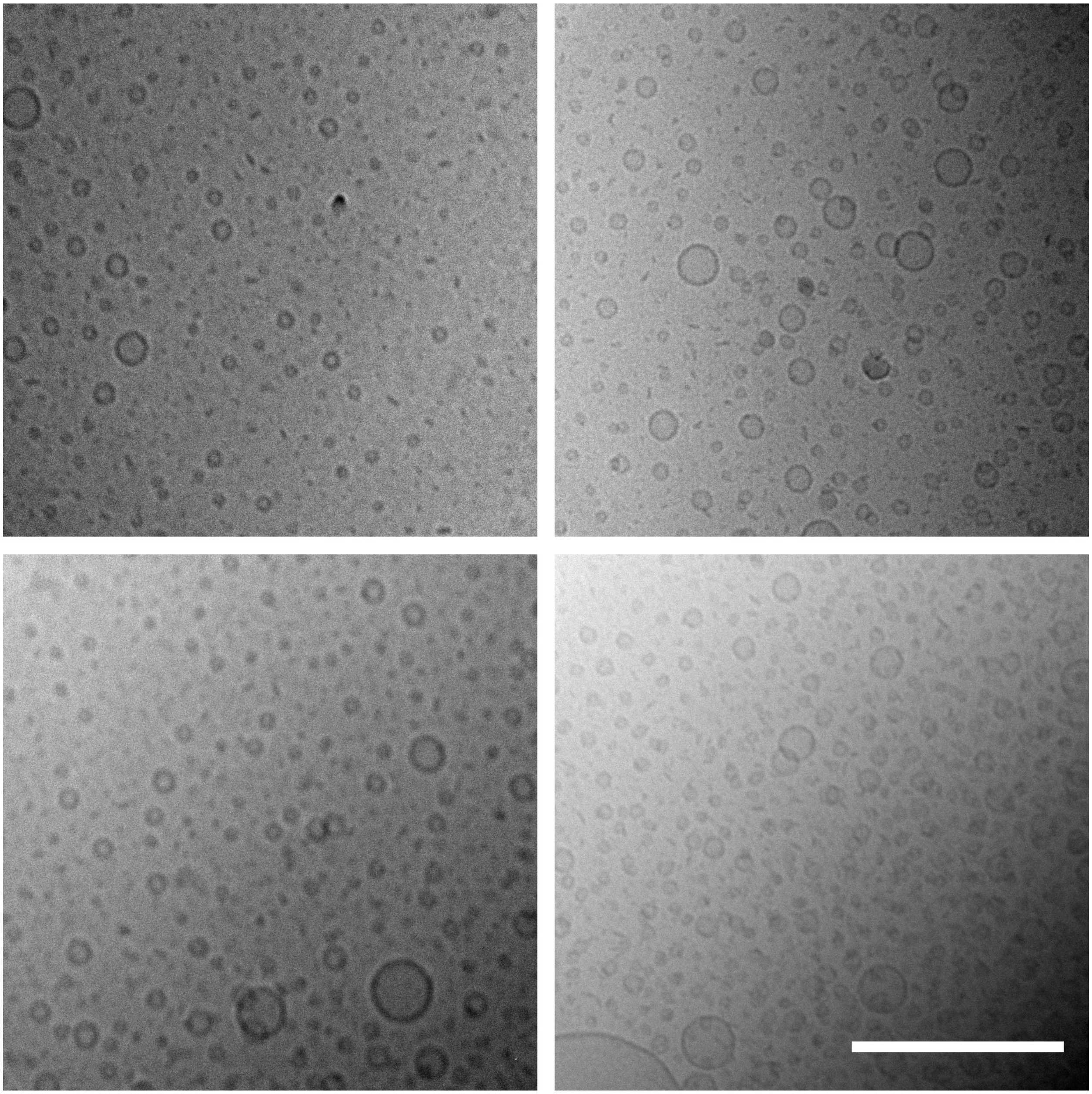

**Figure S17.** Cryo-EM images of a sample containing 80 mol% DOTAP, 3 mol% PTX, and the remainder DOPC. Scale bar: 200 nm.

### 25 mol% PEG2K-lipid

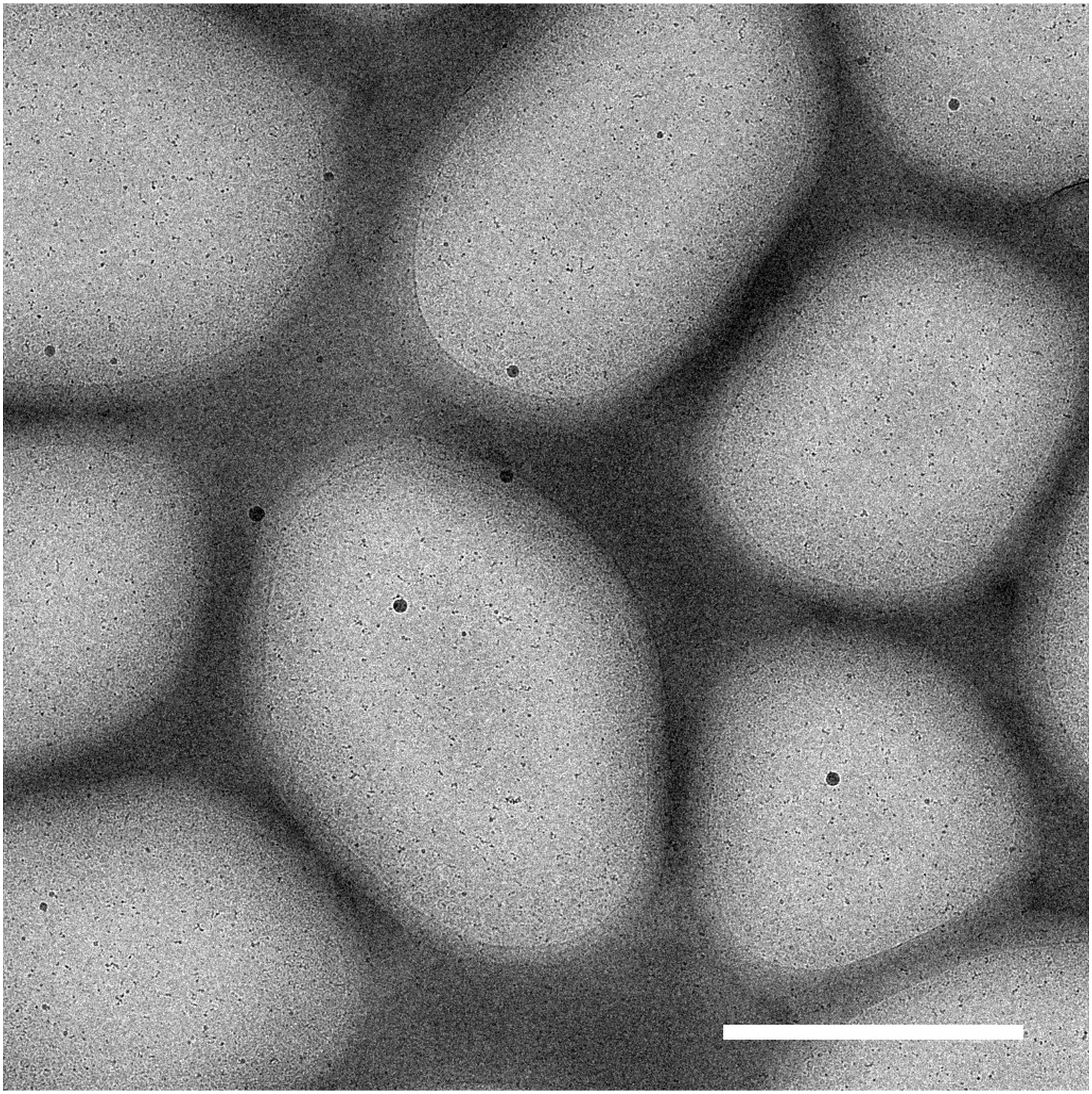

**Figure S18.** Lower magnification cryo-EM image of a sample containing 25 mol% PEG2K-lipid, 3 mol% PTX, and the remainder DOPC. Scale bar: 2 μm.

### 25 mol% PEG2K-lipid

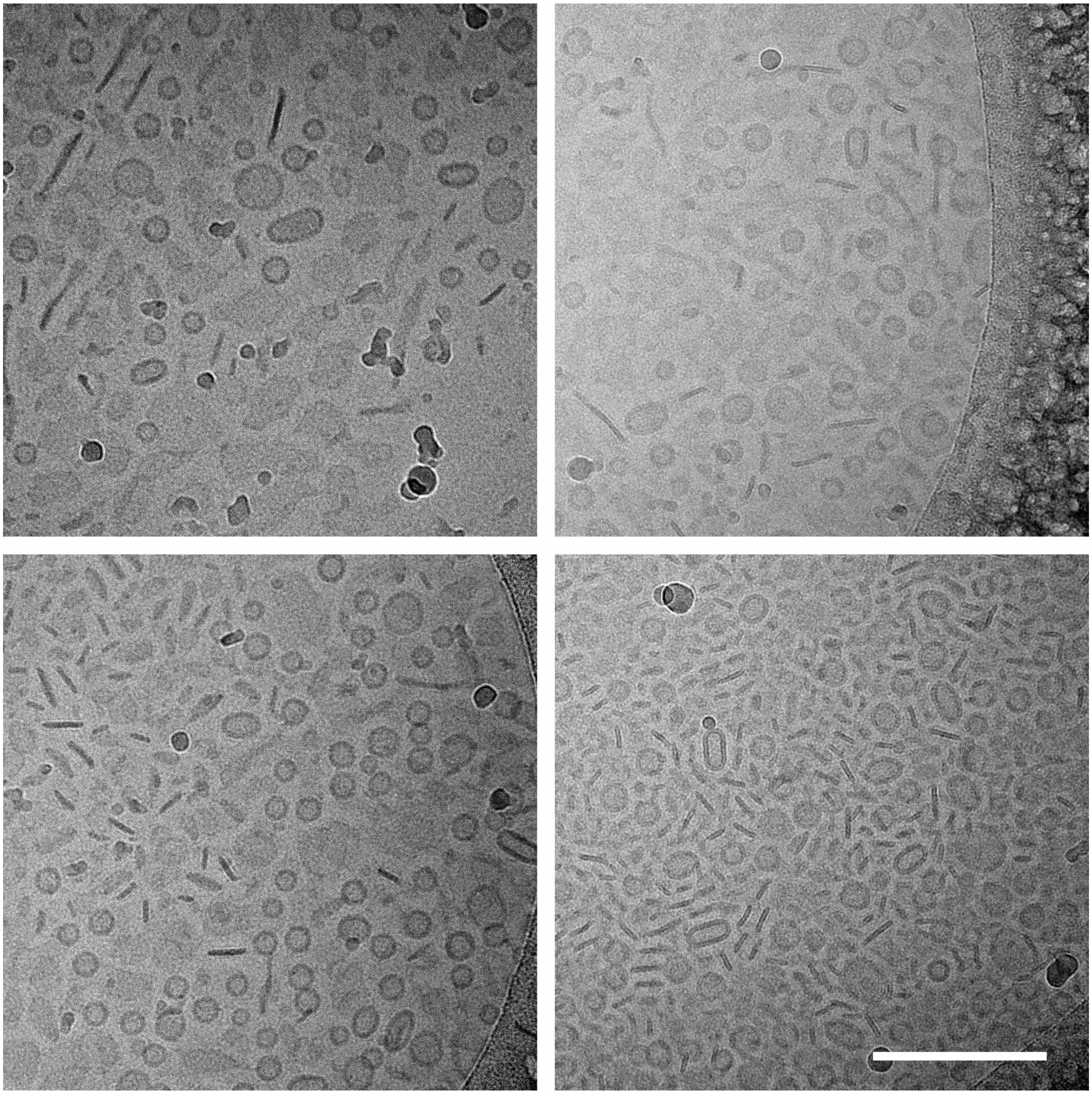

**Figure S19.** Cryo-EM images of a sample containing 25 mol% PEG2K-lipid, 3 mol% PTX, and the remainder DOPC. Scale bar: 200 nm.

### 2 mol% PEG5K-lipid

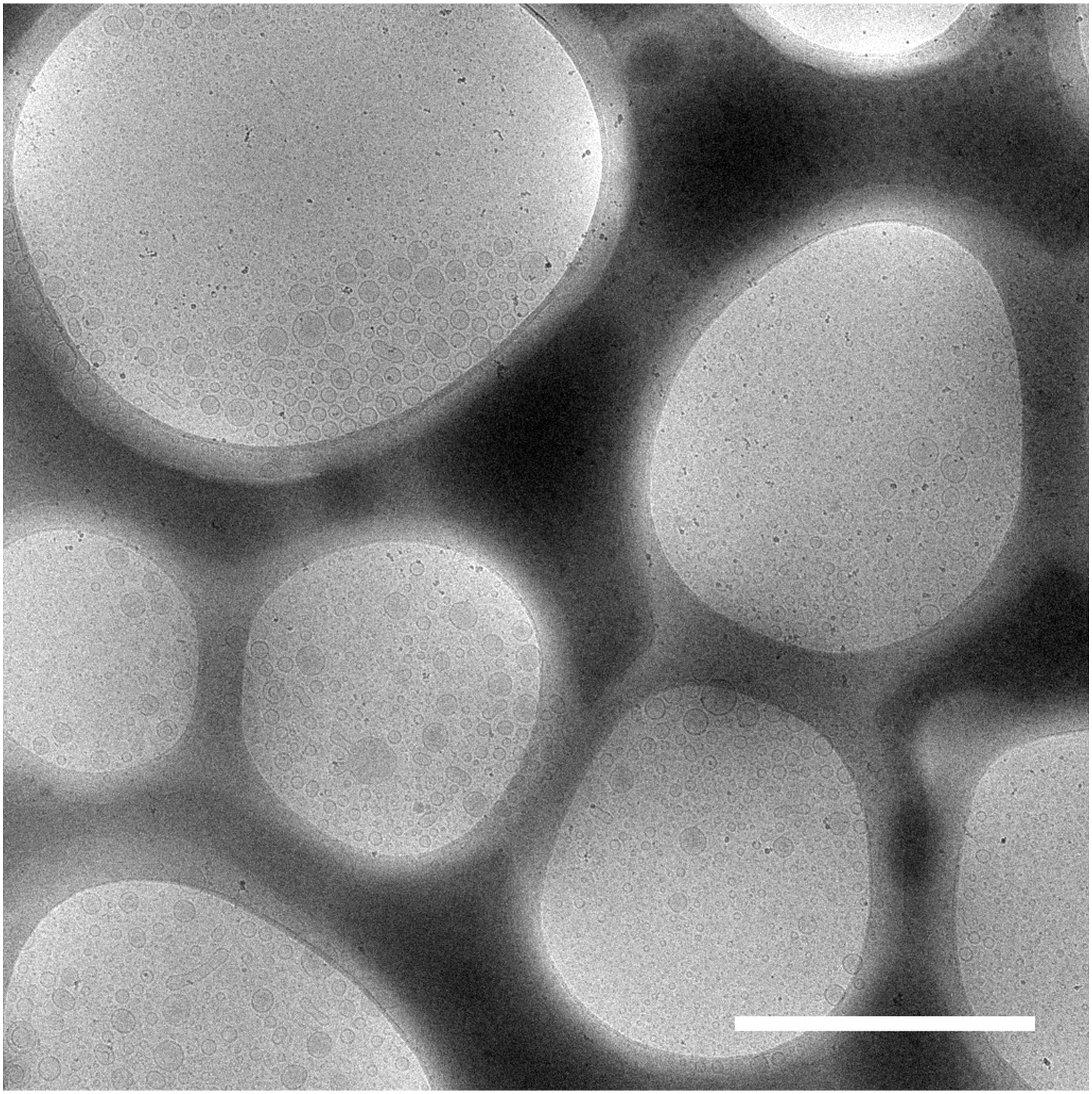

**Figure S20.** Lower magnification cryo-EM image of a sample containing 2 mol% PEG5K-lipid, 3 mol% PTX, and the remainder DOPC. Scale bar: 2 μm.

### 2 mol% PEG5K-lipid

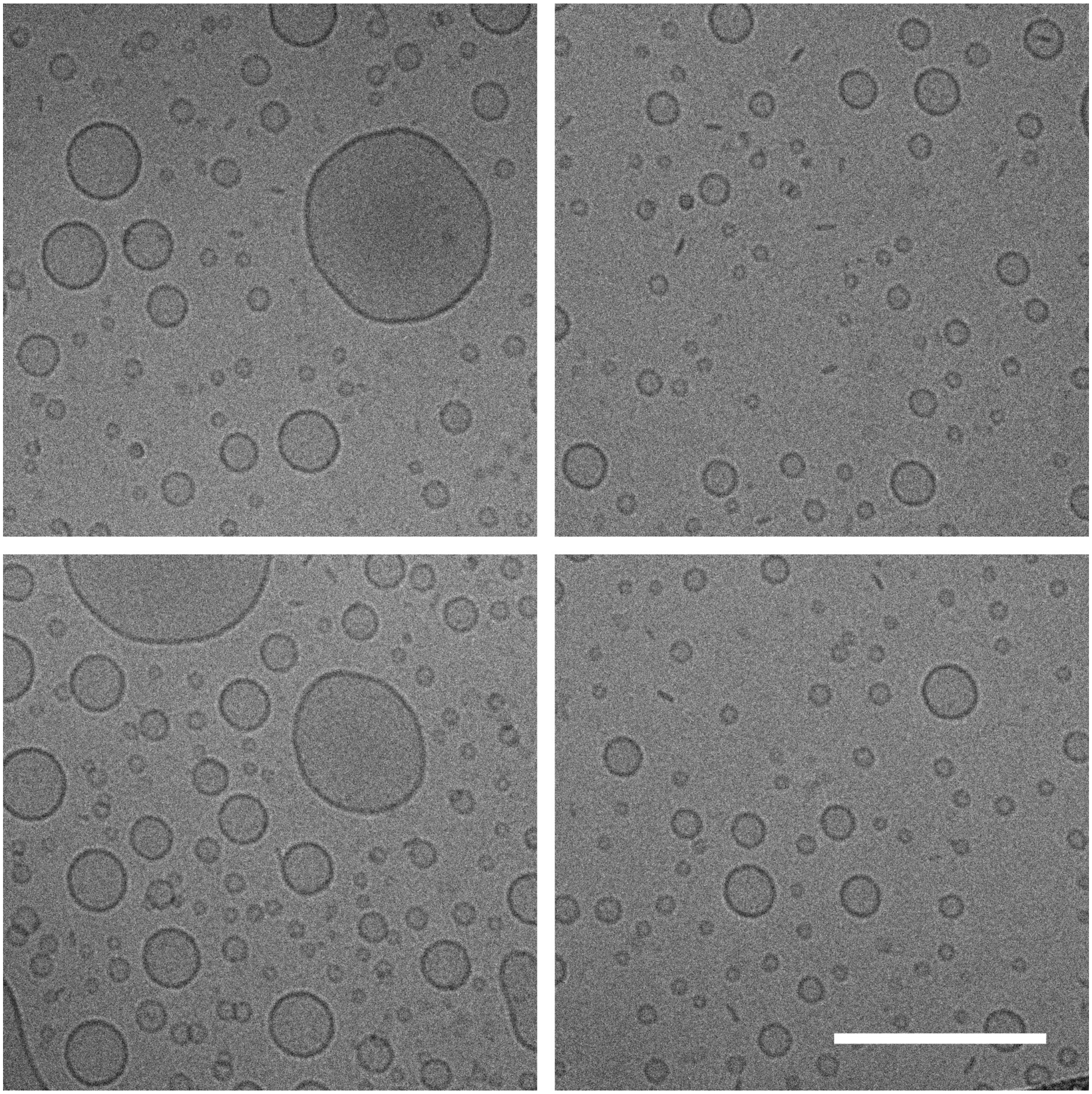

**Figure S21.** Cryo-EM images of a sample containing 2 mol% PEG5K-lipid, 3 mol% PTX, and the remainder DOPC. Scale bar: 200 nm.

### 25 mol% PEG5K-lipid

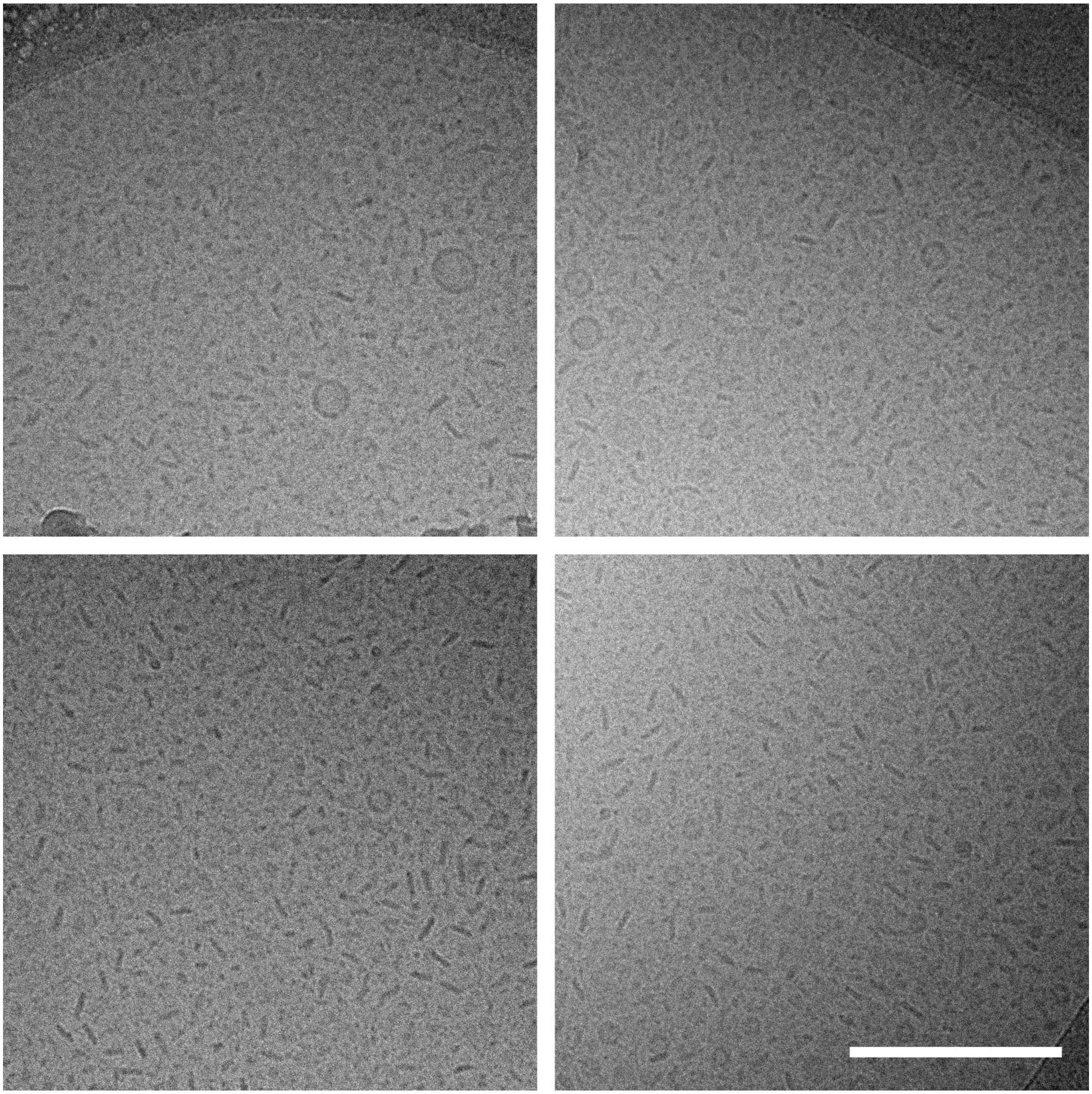

**Figure S22.** Cryo-EM images of a sample containing 25 mol% PEG5K-lipid, 3 mol% PTX, and the remainder DOPC. Scale bar: 200 nm.

### 25 mol% PEG5K-lipid

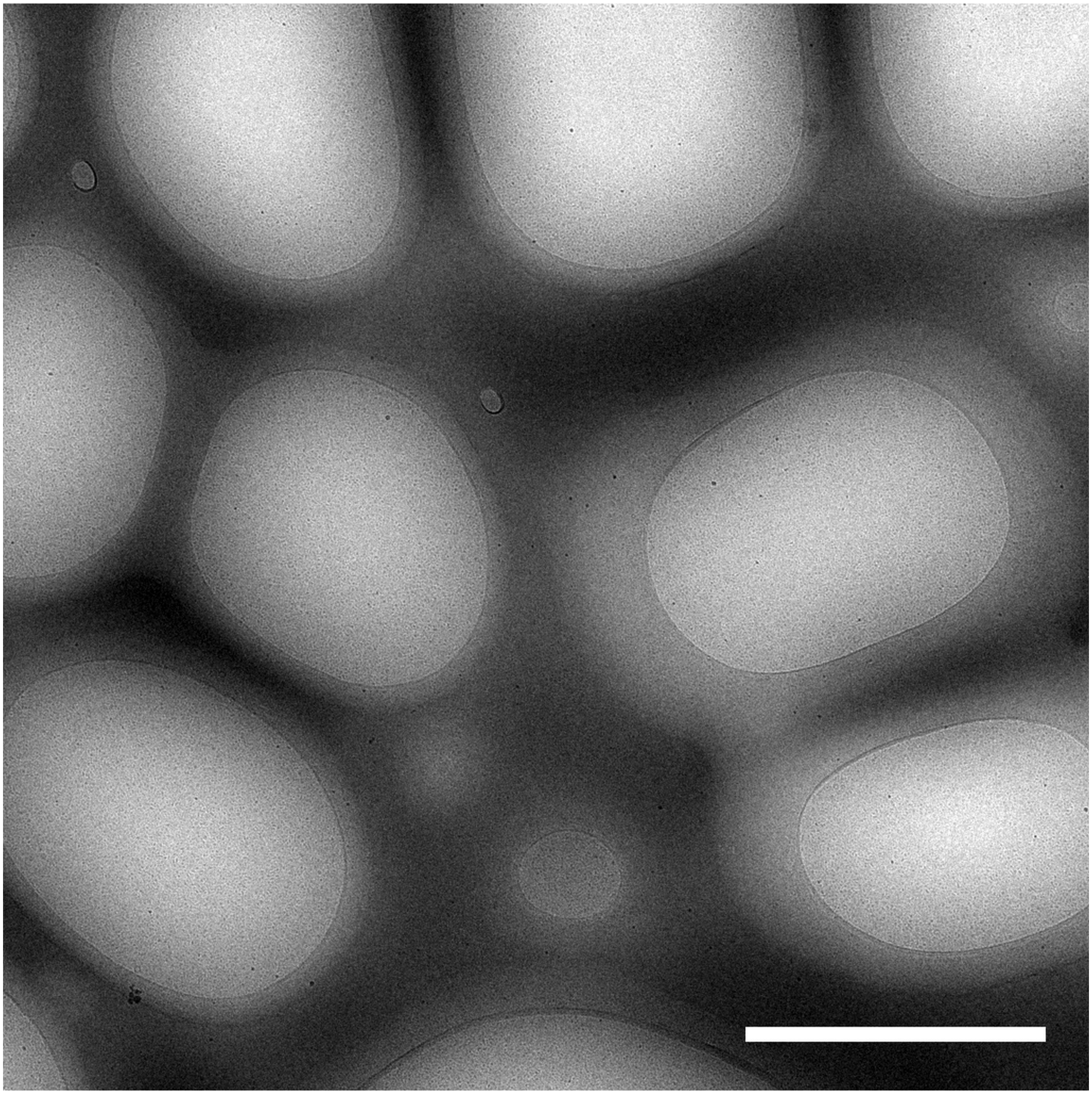

**Figure S23.** Lower magnification cryo-EM image of a sample containing 25 mol% PEG5K-lipid, 3 mol% PTX, and the remainder DOPC. Scale bar: 2 μm.

### 10 mol% MVL5, 10 mol% PEG2K-lipid

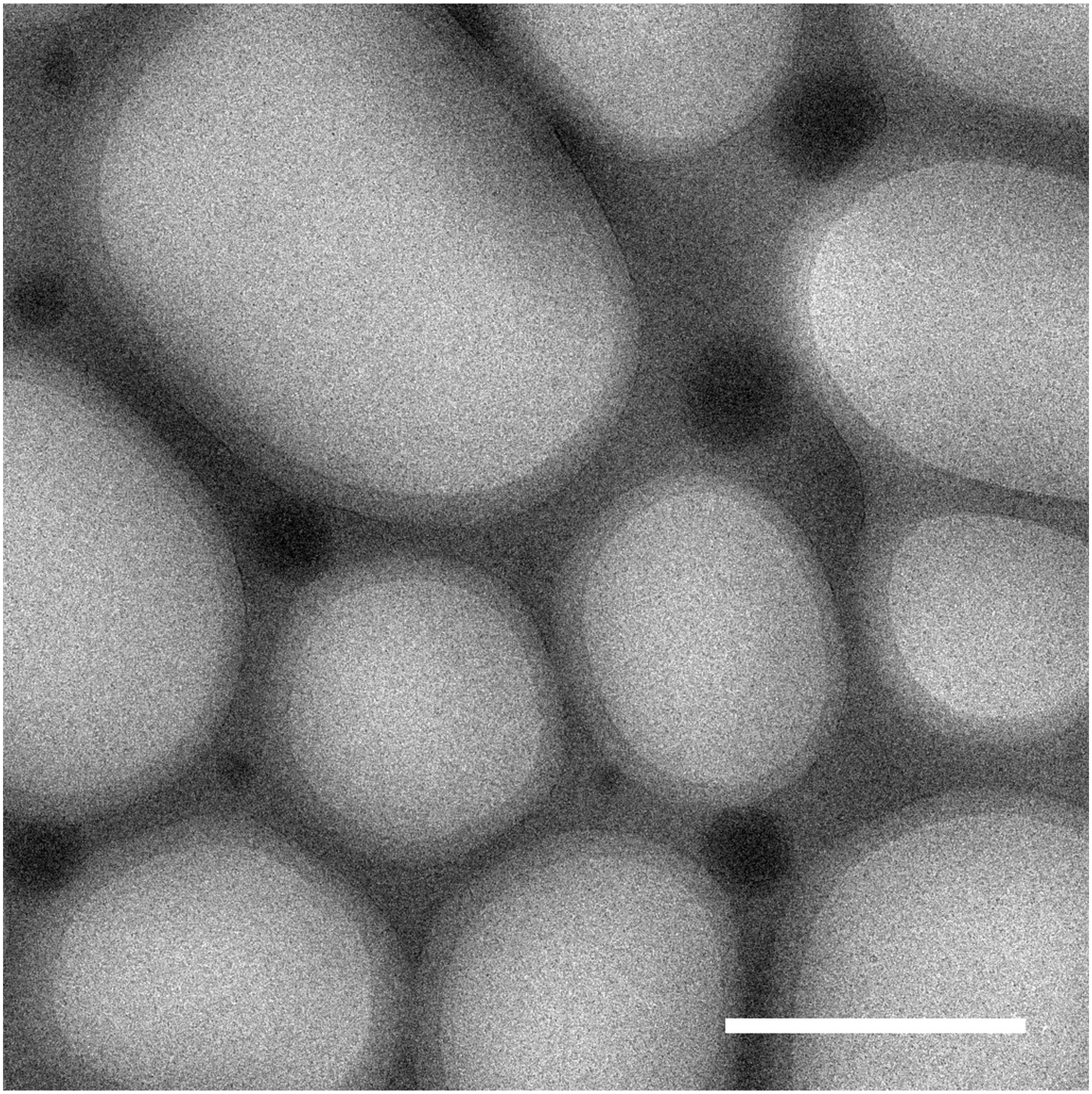

**Figure S24.** Lower magnification cryo-EM image of a sample containing 10 mol% MVL5, 10 mol% PEG2K-lipid, 3 mol% PTX, and the remainder DOPC. Scale bar: 2 μm.

### 10 mol% MVL5, 10 mol% PEG2K-lipid

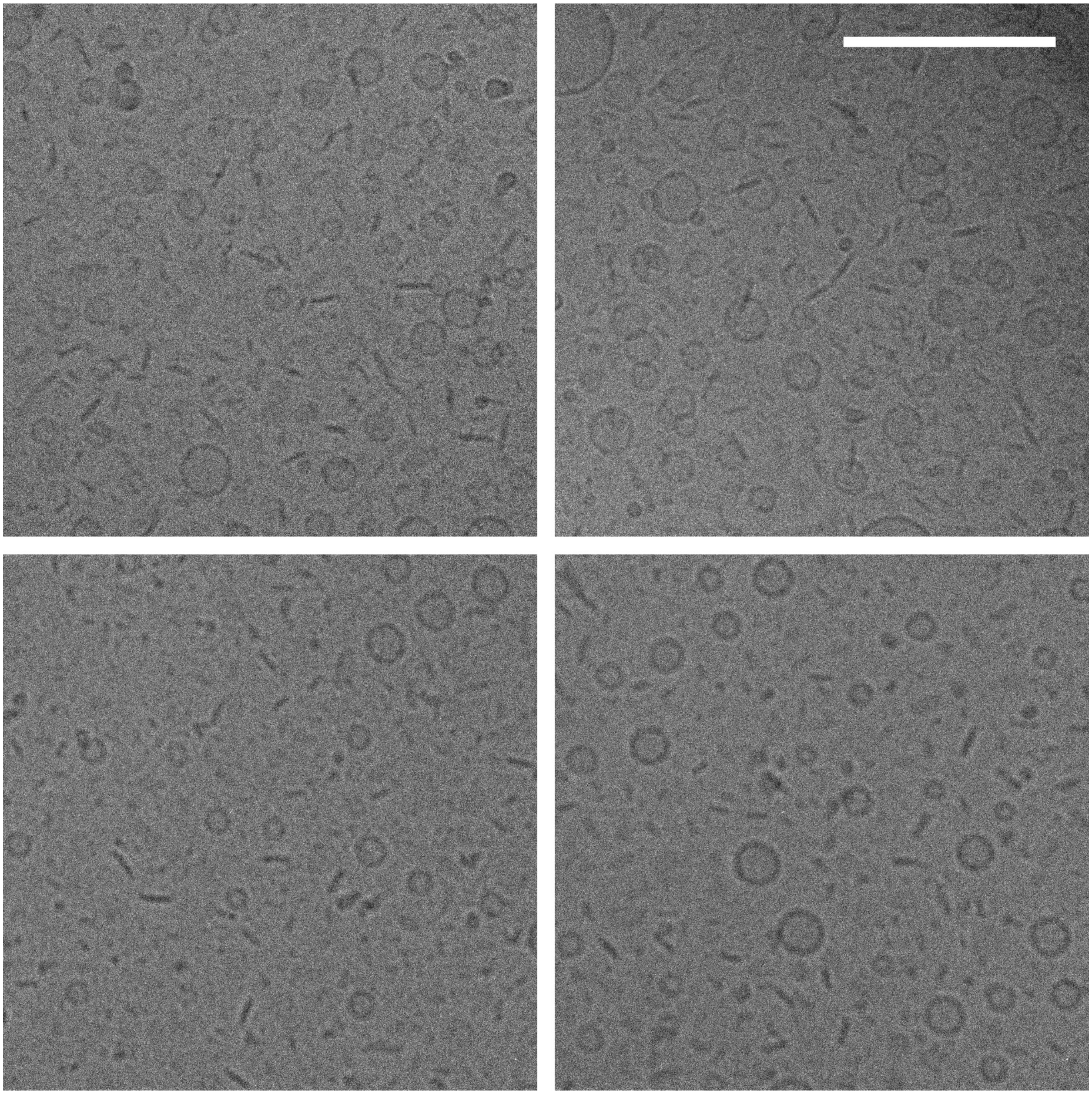

**Figure S25.** Cryo-EM images of a sample containing 10 mol% MVL5, 10 mol% PEG2K-lipid, 3 mol% PTX, and the remainder DOPC. Scale bar: 200 nm.

### 50 mol% MVL5, 10 mol% PEG2K-lipid

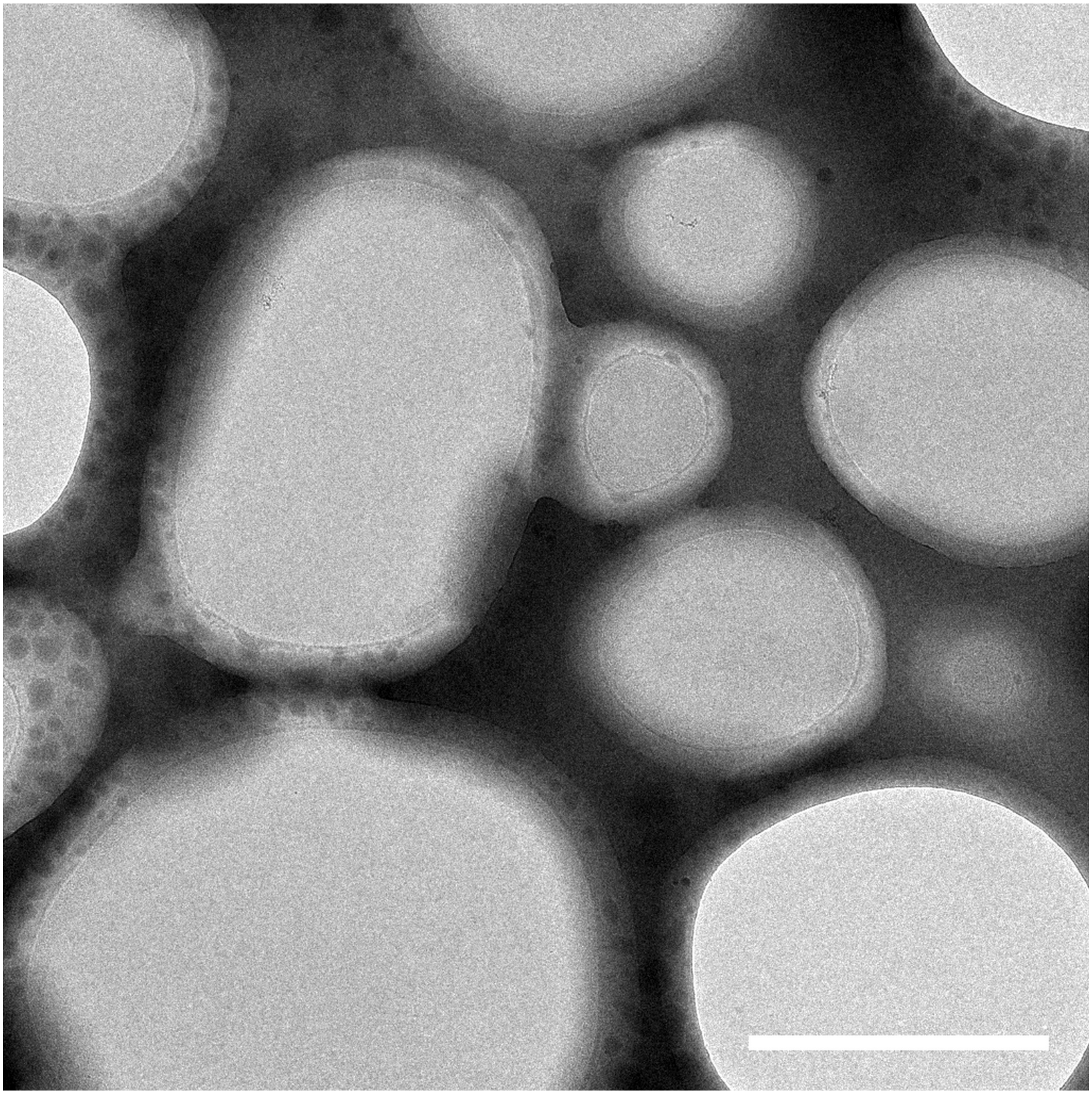

**Figure S26.** Lower magnification cryo-EM image of a sample containing 50 mol% MVL5, 10 mol% PEG2K-lipid, 3 mol% PTX, and the remainder DOPC. Scale bar: 2 μm.

### 50 mol% MVL5, 10 mol% PEG2K-lipid

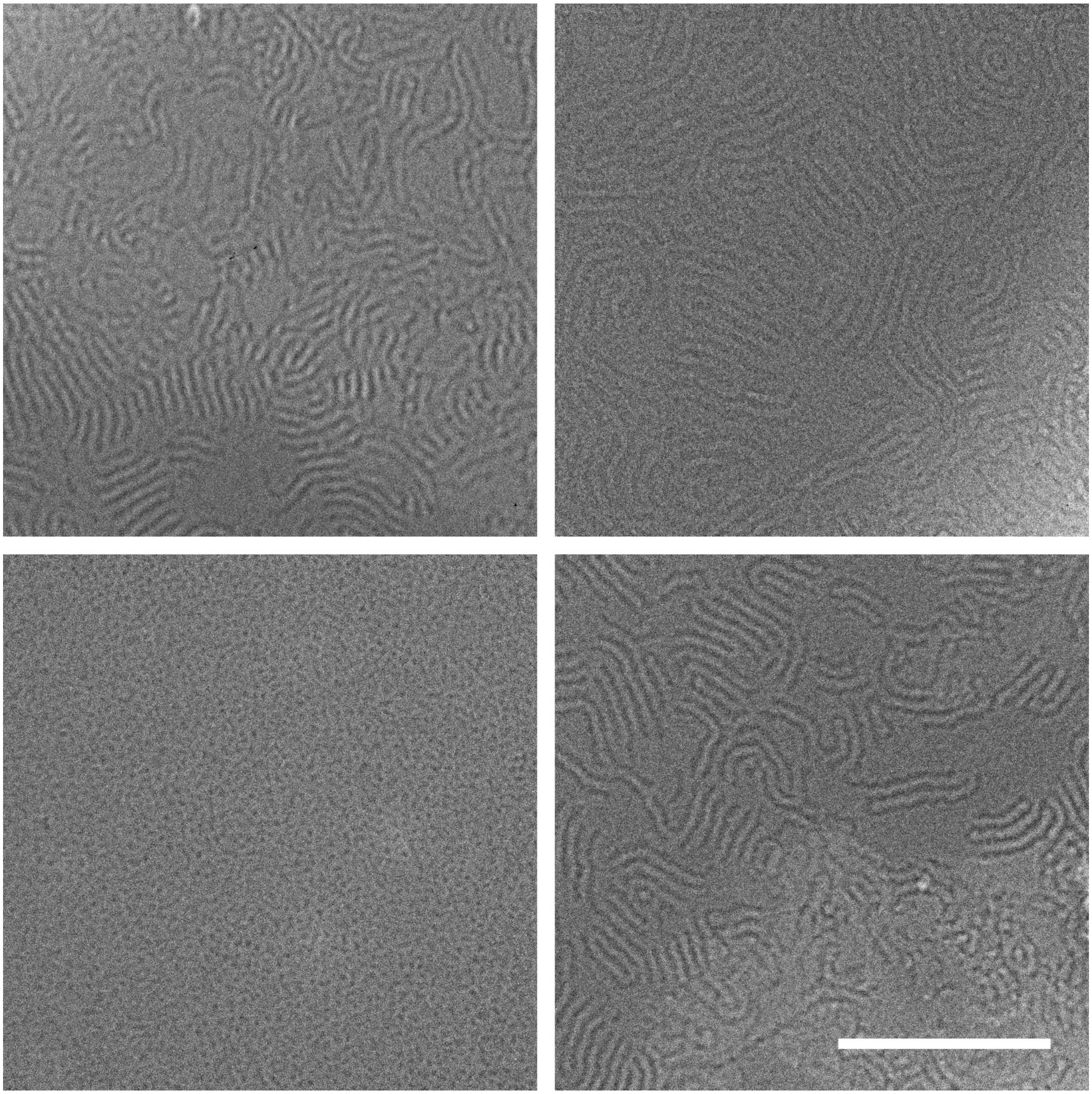

**Figure S27.** Cryo-EM images of a sample containing 50 mol% MVL5, 10 mol% PEG2K-lipid, 3 mol% PTX, and the remainder DOPC. Scale bar: 200 nm.

### 75 mol% MVL5, 10 mol% PEG2K-lipid

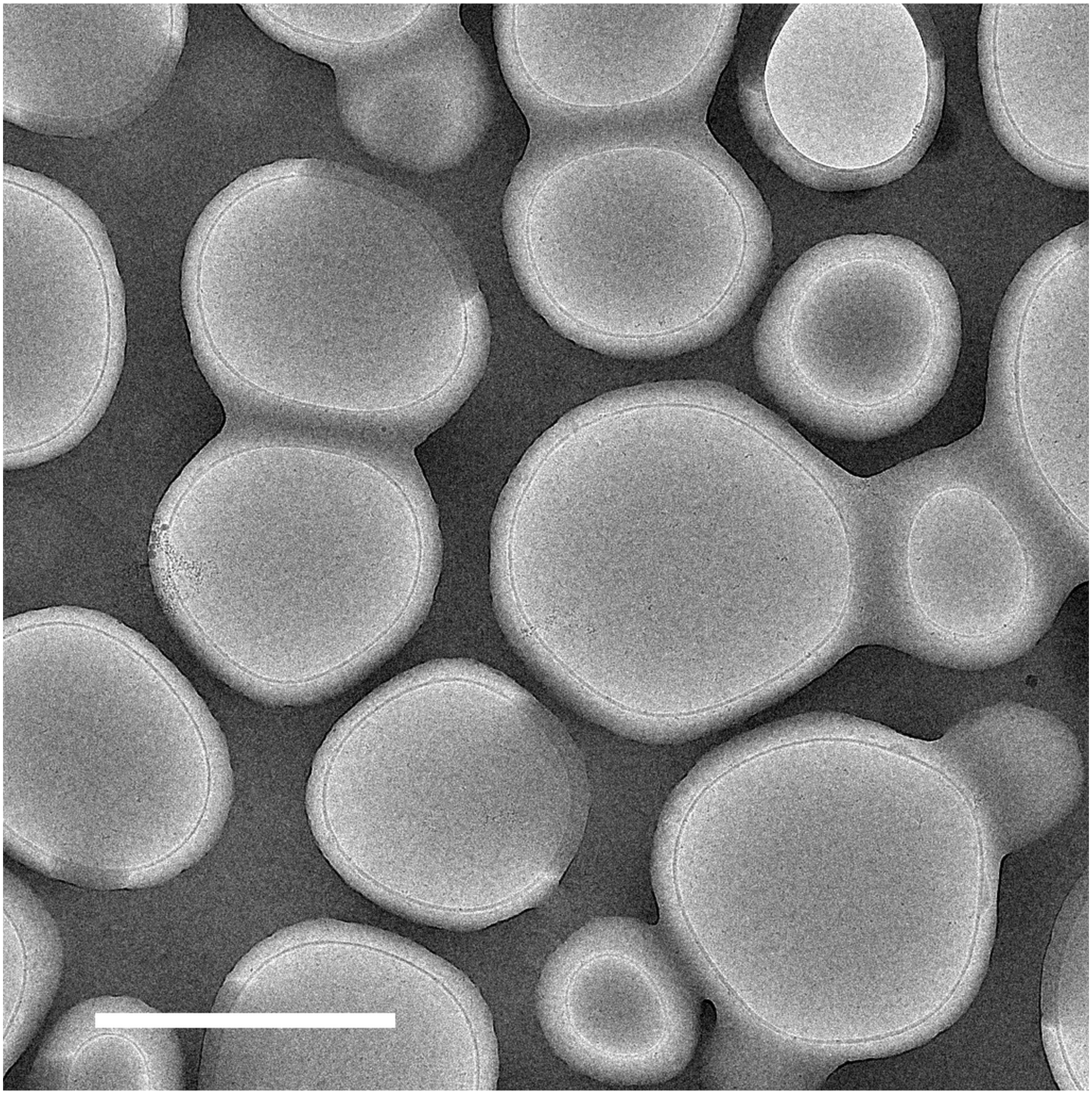

**Figure S28.** Lower magnification cryo-EM image of a sample containing 75 mol% MVL5, 10 mol% PEG2K-lipid, 3 mol% PTX, and the remainder DOPC. Scale bar: 2 μm.

### 75 mol% MVL5, 10 mol% PEG2K-lipid (Worms)

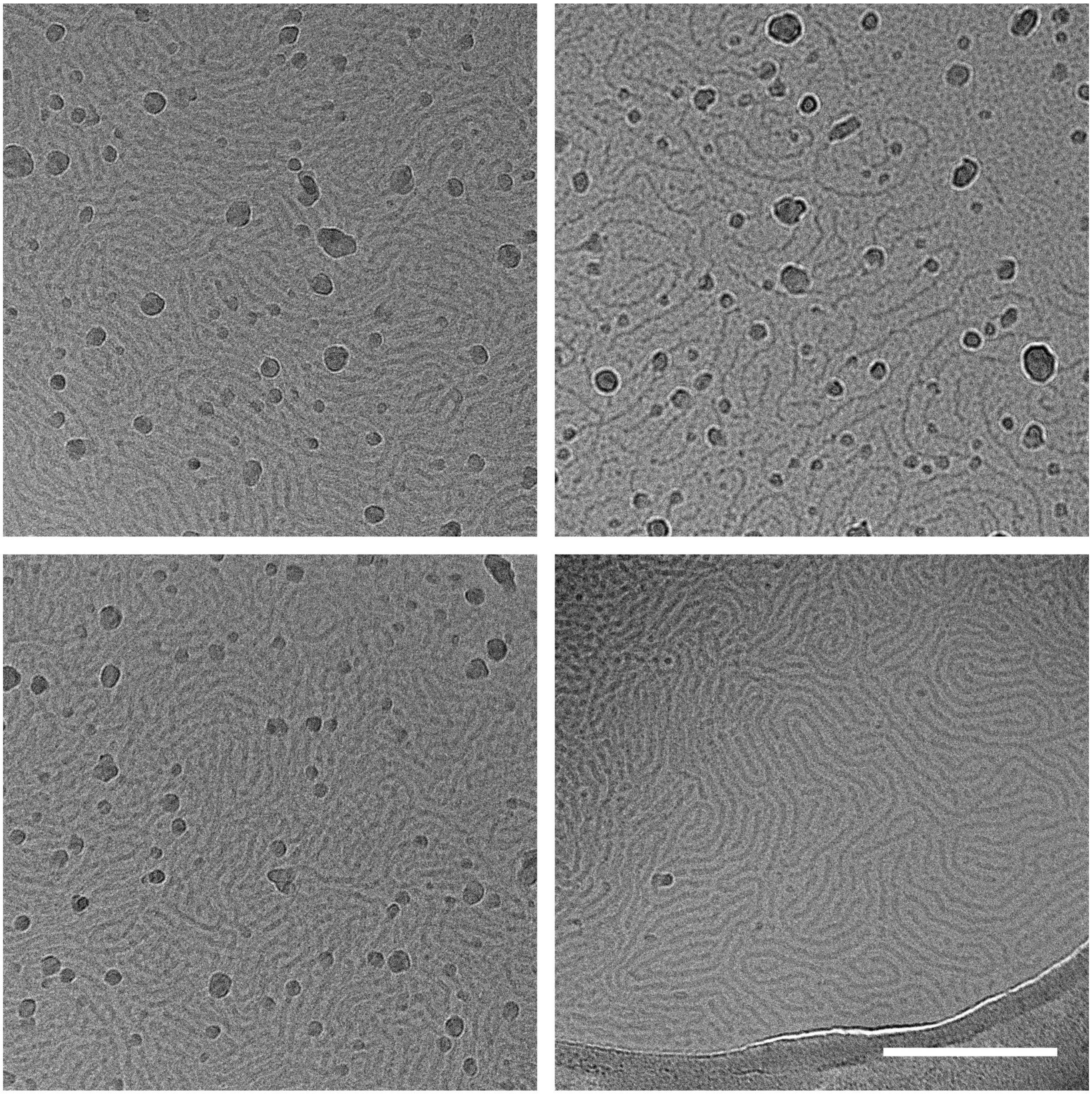

**Figure S29.** Cryo-EM images of a sample containing 75 mol% MVL5, 10 mol% PEG2K-lipid, 3 mol% PTX, and the remainder DOPC. Scale bar: 200 nm.

### 75 mol% MVL5, 10 mol% PEG2K-lipid (Spheres)

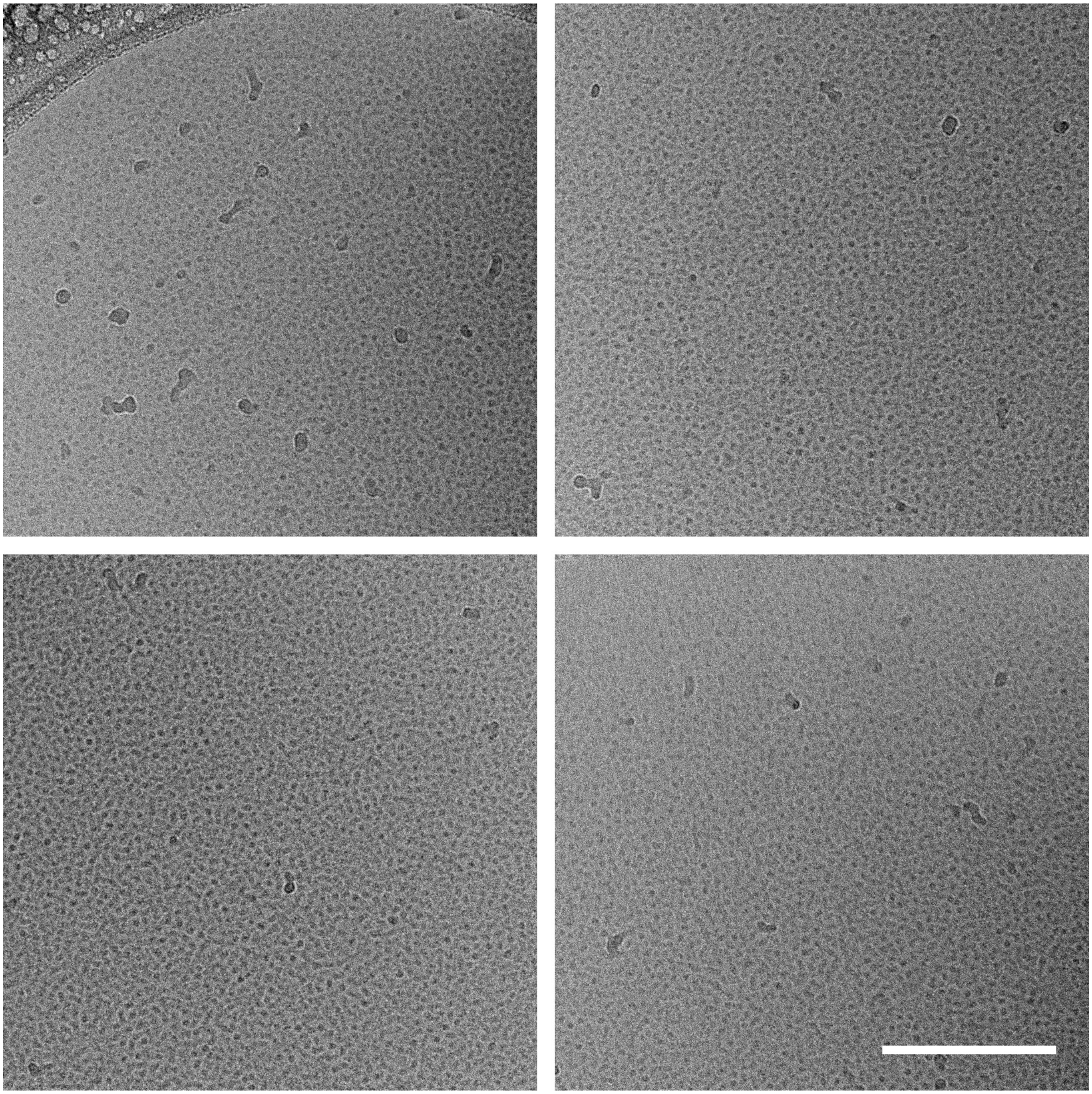

**Figure S30.** Cryo-EM images of a sample containing 75 mol% MVL5, 10 mol% PEG2K-lipid, 3 mol% PTX, and the remainder DOPC. Scale bar: 200 nm.

### Possible Size Sorting by Ice Thickness Gradients (50 mol% DOTAP)

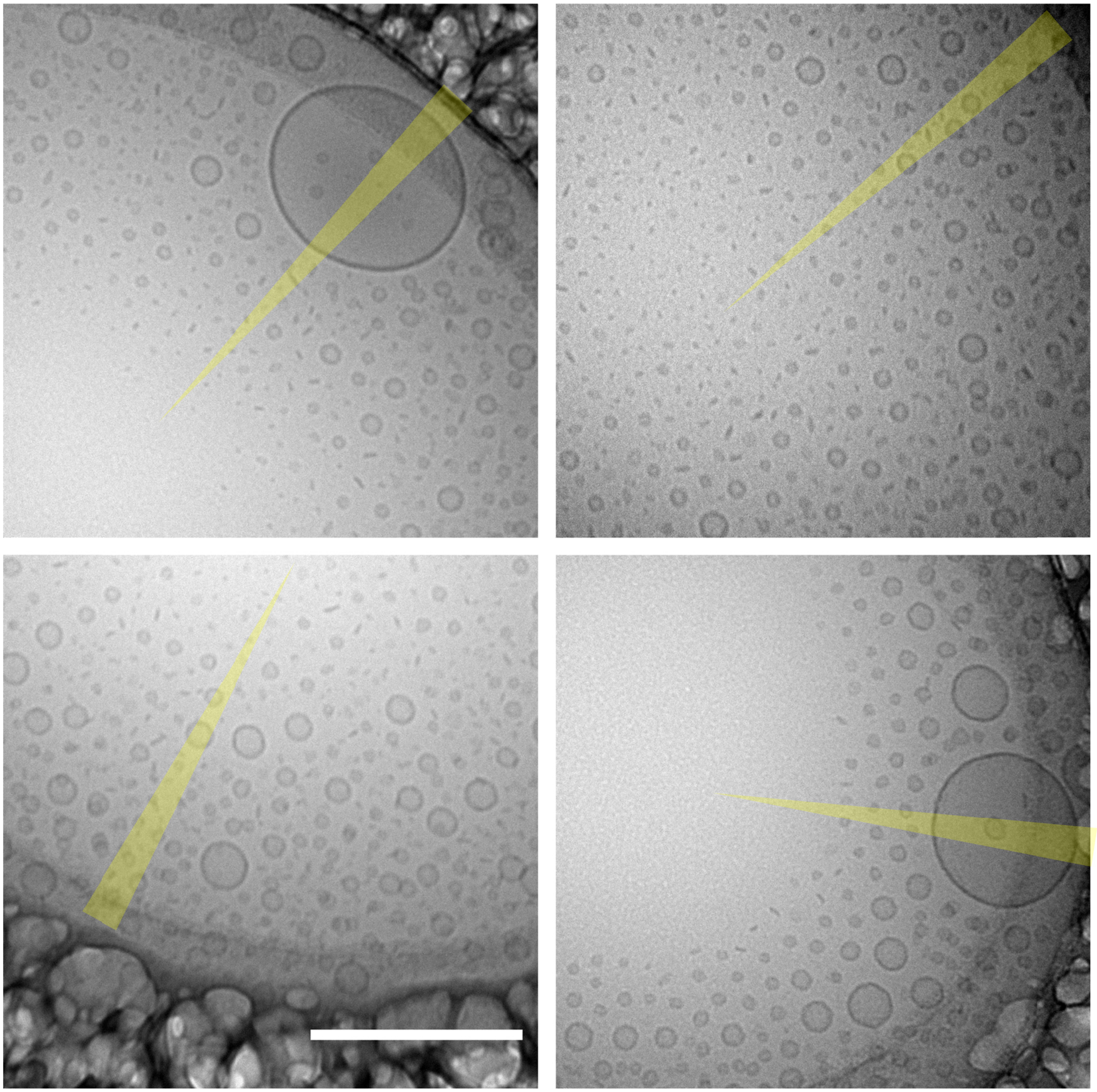

**Figure S31.** Cryo-EM images exhibiting possible sorting of objects by a gradient in ice thickness (indicated by yellow wedges). Sample composition: 50 mol% DOTAP, 3 mol% PTX, and the remainder DOPC. Scale bar: 200 nm.

### Effect of added DOTAP at 10 mol% PEG2K-lipid

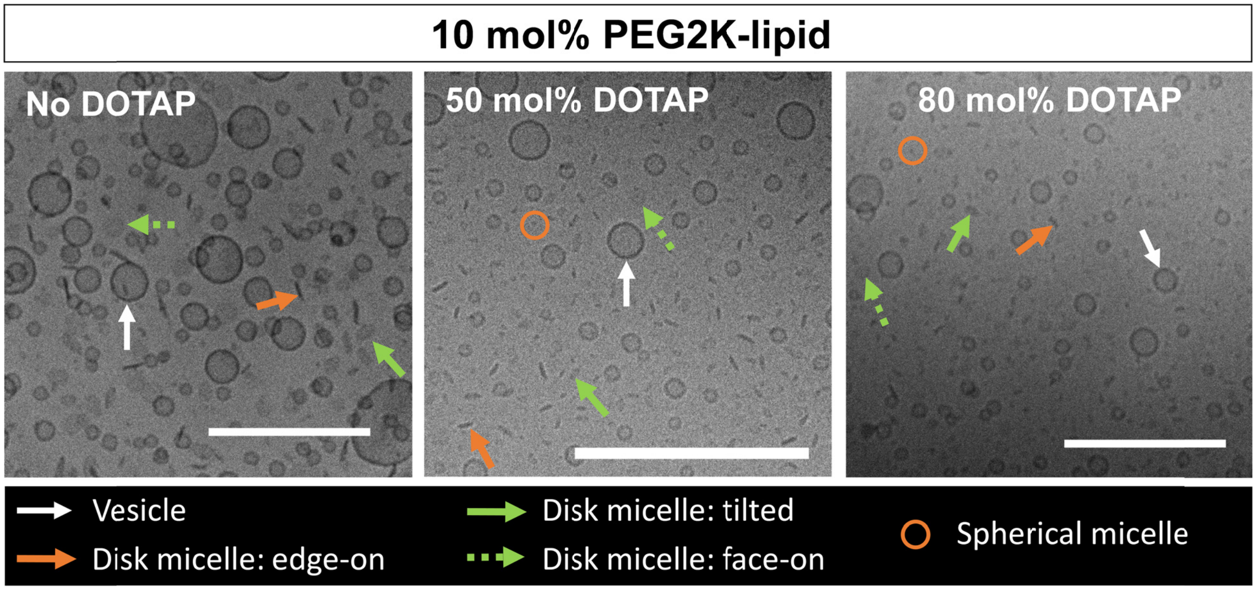


**Figure S32.** Cryo-TEM images demonstrating the effect of added univalent cationic DOTAP on the lipid assembly structures of a formulation with 10 mol% PEG2K-lipid. The formulations contain DOTAP as specified and 10 mol% PEG2K-lipid, with DOPC the remainder. Arrows and circles identify vesicle and micelle structures according to the legend. Scale bars: 200 nm. For unlabeled and additional micrographs, see Figures S12, S13, S33–S36.

### 50 mol% DOTAP, 10 mol% PEG2K-lipid

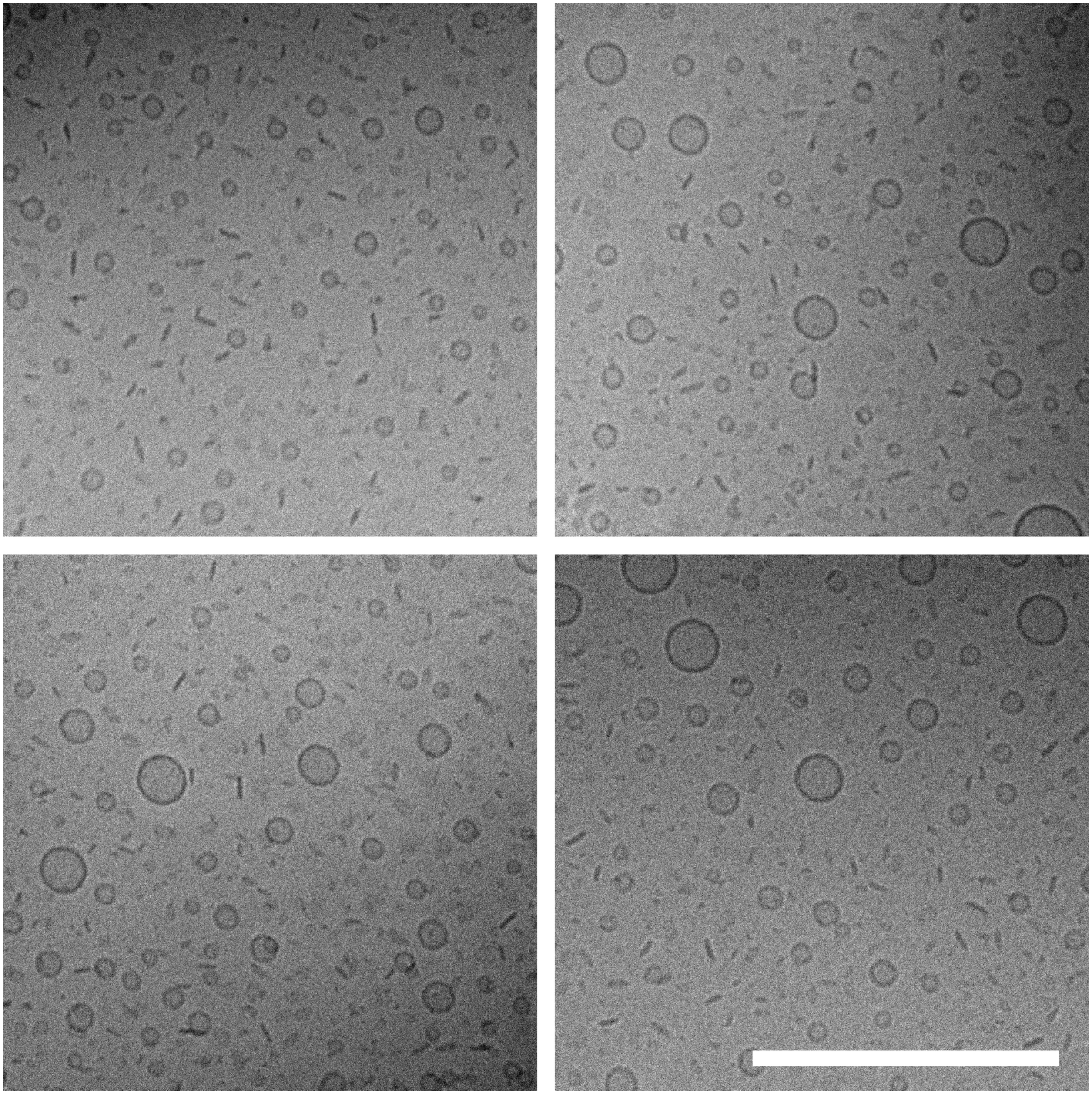

**Figure S33.** Cryo-EM images of a sample containing 50 mol% DOTAP, 10 mol% PEG2K-lipid, 3 mol% PTX, and the remainder DOPC. Scale bar: 200 nm.

### 50 mol% DOTAP, 10 mol% PEG2K-lipid

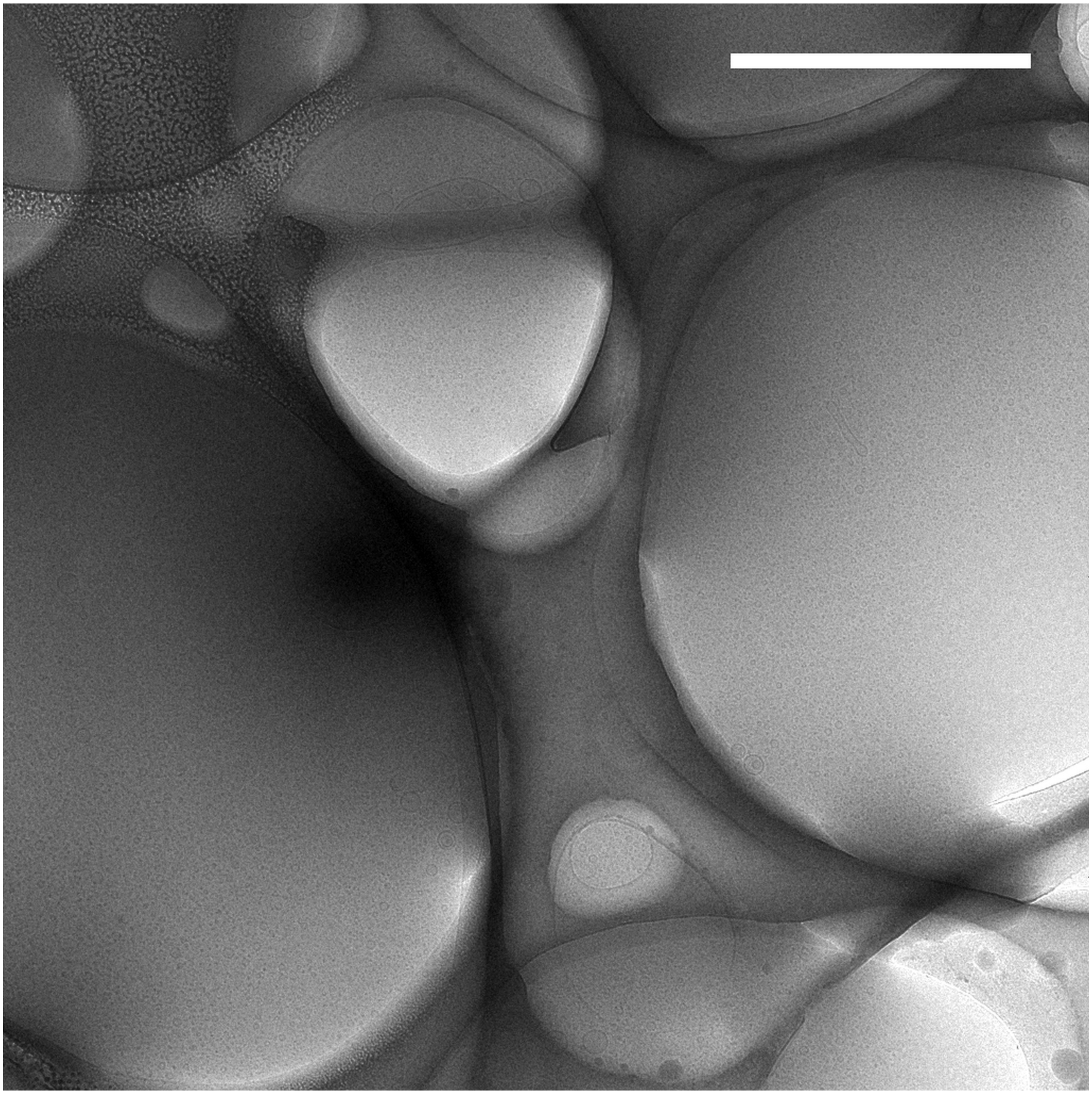

**Figure S34.** Lower magnification cryo-EM image of a sample containing 50 mol% DOTAP, 10 mol% PEG2K-lipid, 3 mol% PTX, and the remainder DOPC. Scale bar: 2 μm.

### 80 mol% DOTAP, 10 mol% PEG2K-lipid

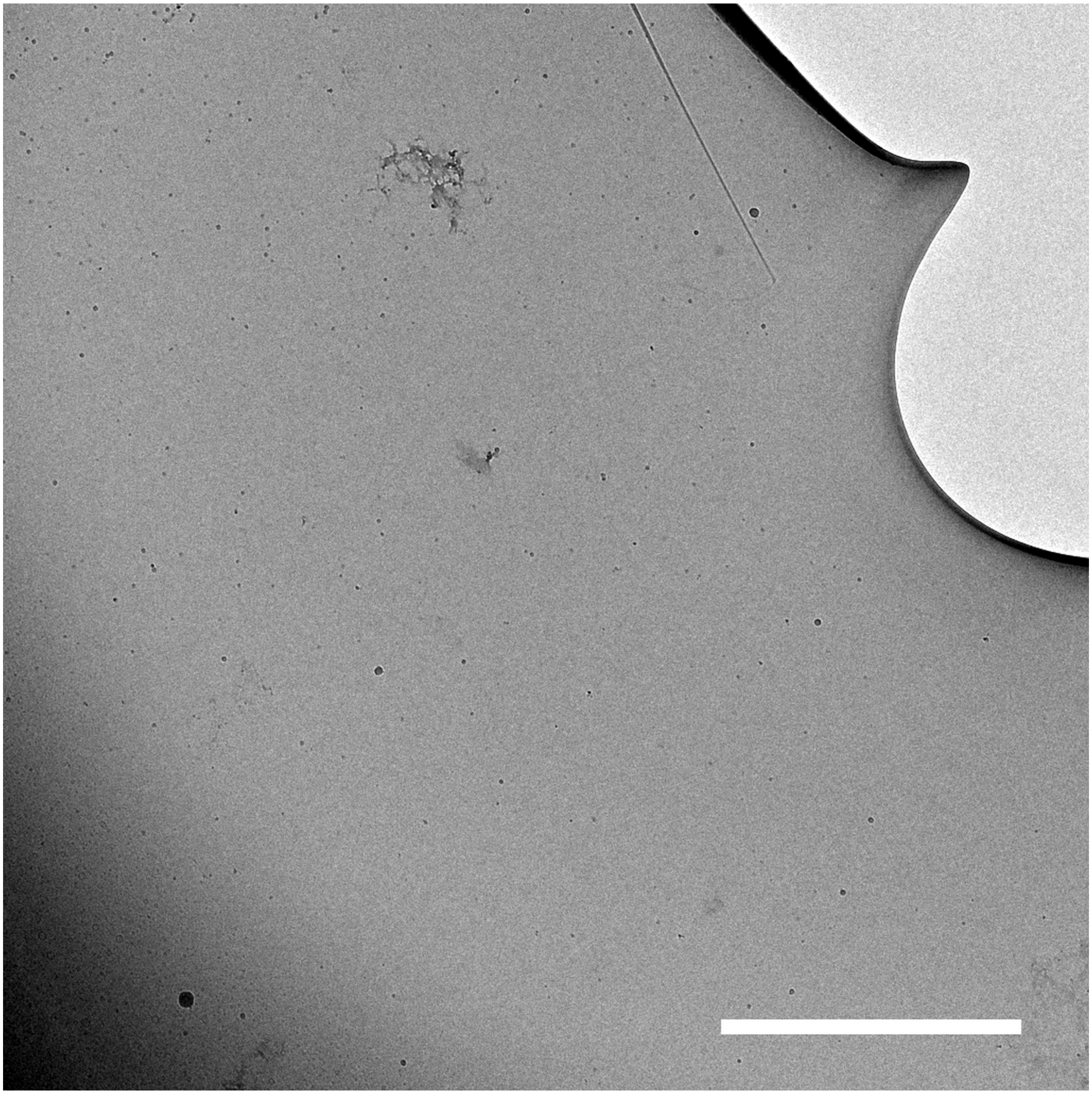

**Figure S35.** Lower magnification cryo-EM image of a sample containing 80 mol% DOTAP, 10 mol% PEG2K-lipid, 3 mol% PTX, and the remainder DOPC. Scale bar: 2 μm.

### 80 mol% DOTAP, 10 mol% PEG2K-lipid

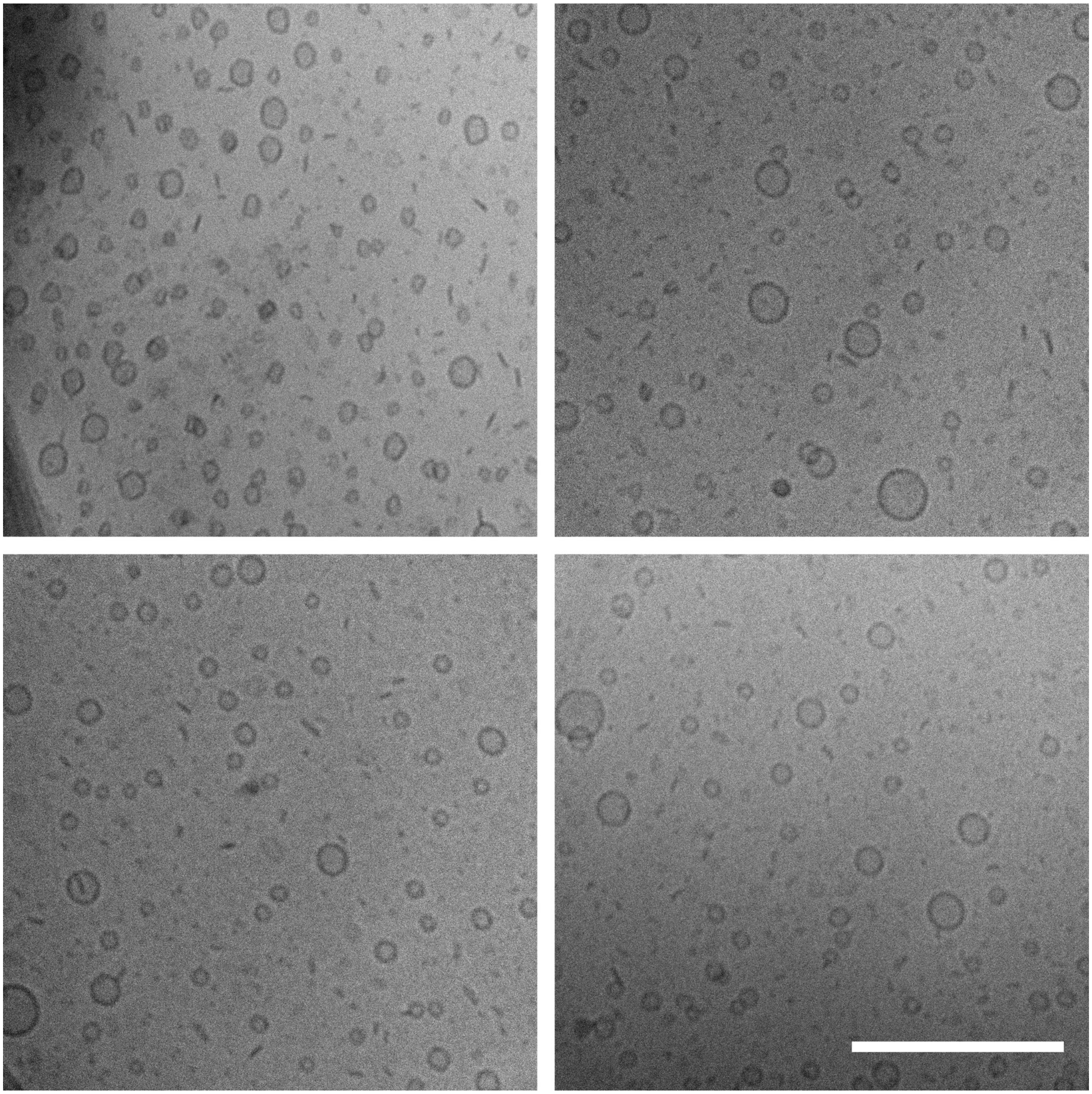

**Figure S36.** Cryo-EM images of a sample containing 80 mol% DOTAP, 10 mol% PEG2K-lipid, 3 mol% PTX, and the remainder DOPC. Scale bar: 200 nm.

### 50 mol% DOTAP, 25 mol% PEG2K-lipid

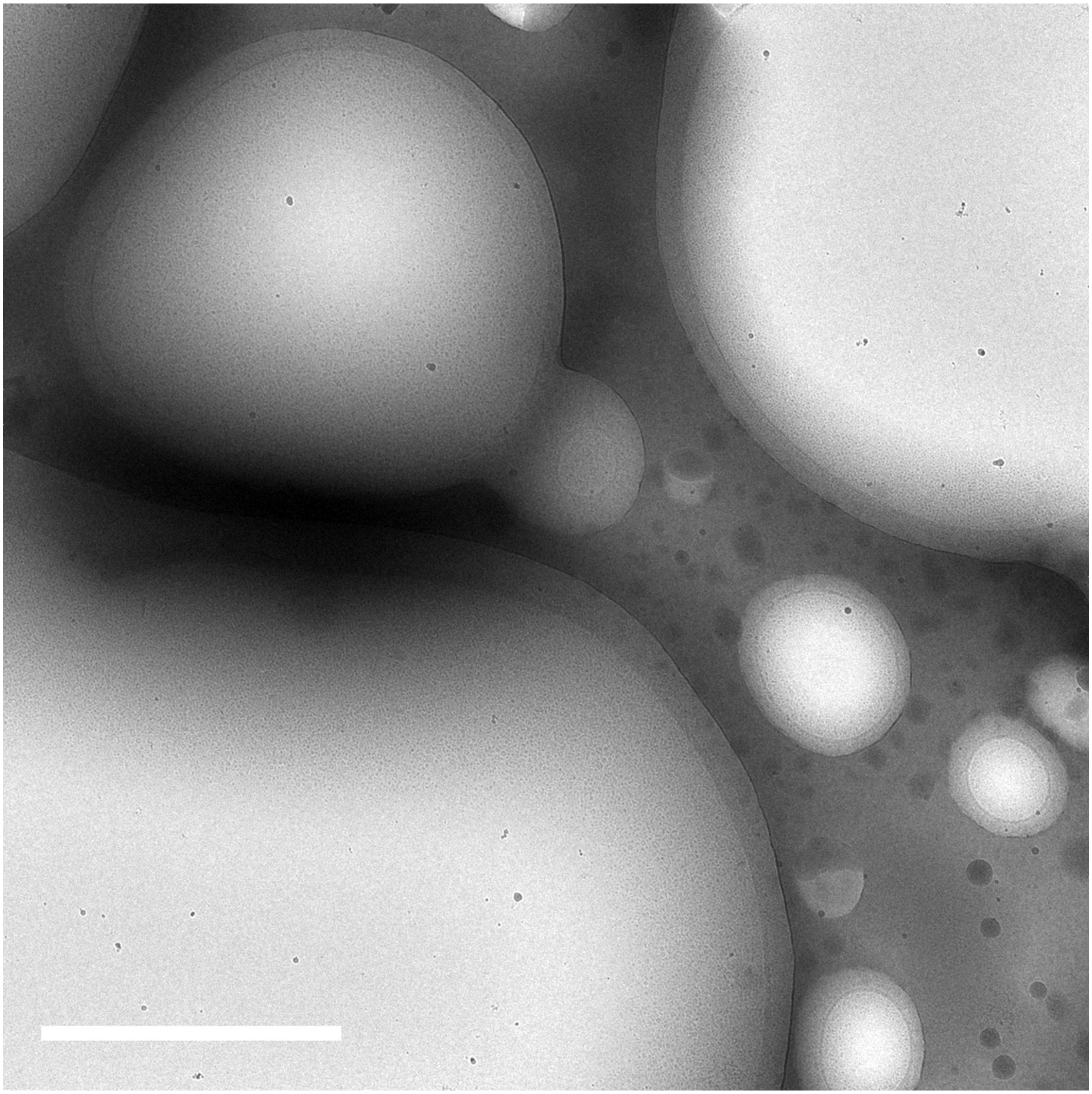

**Figure S37.** Lower magnification cryo-EM image of a sample containing 50 mol% DOTAP, 25 mol% PEG2K-lipid, 3 mol% PTX, and the remainder DOPC. Scale bar: 2 μm.

### 50 mol% DOTAP, 25 mol% PEG2K-lipid

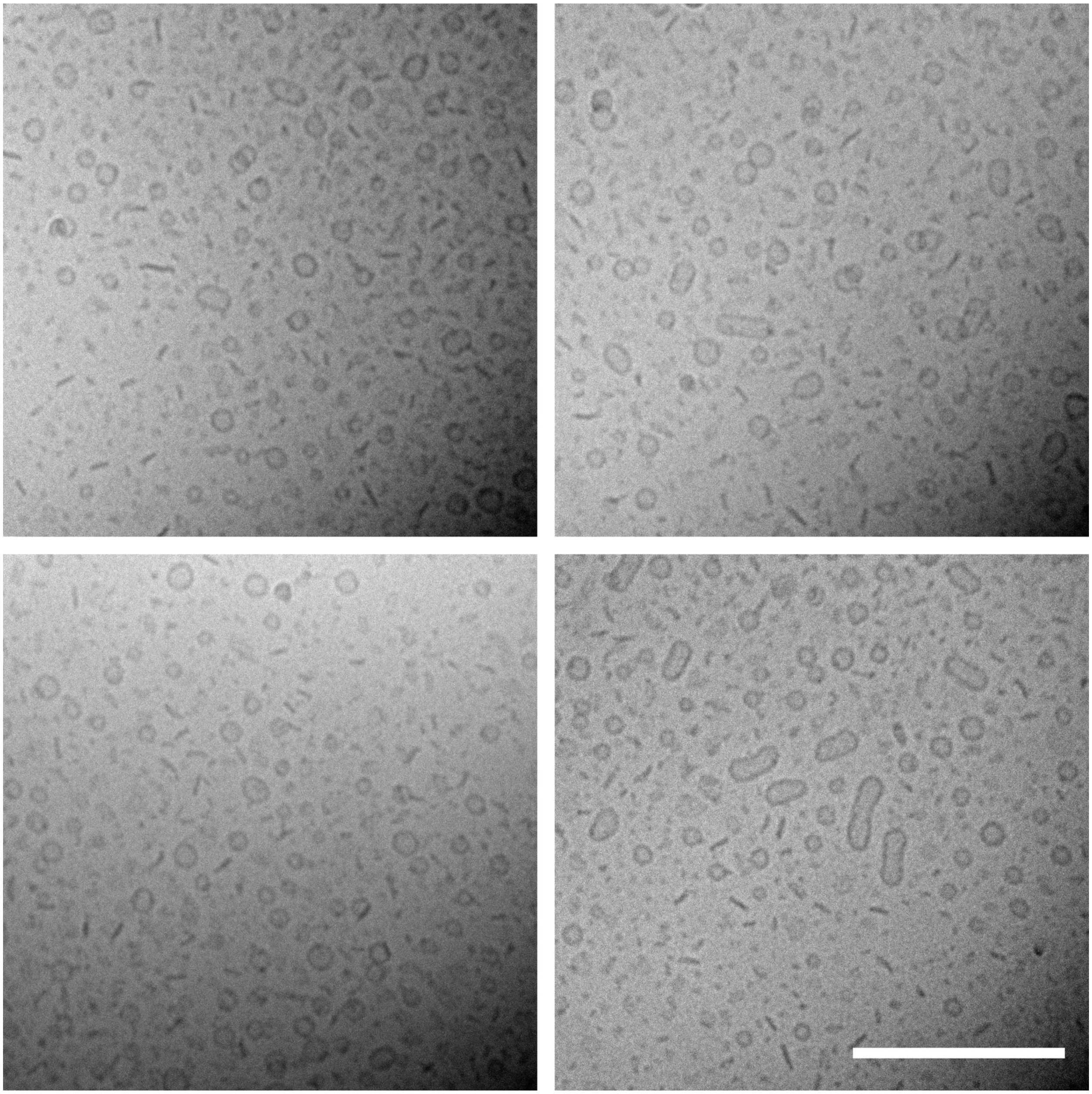

**Figure S38.** Cryo-EM images of a sample containing 50 mol% DOTAP, 25 mol% PEG2K-lipid, 3 mol% PTX, and the remainder DOPC. Scale bar: 200 nm.

### 50 mol% DOTAP (unsonicated)

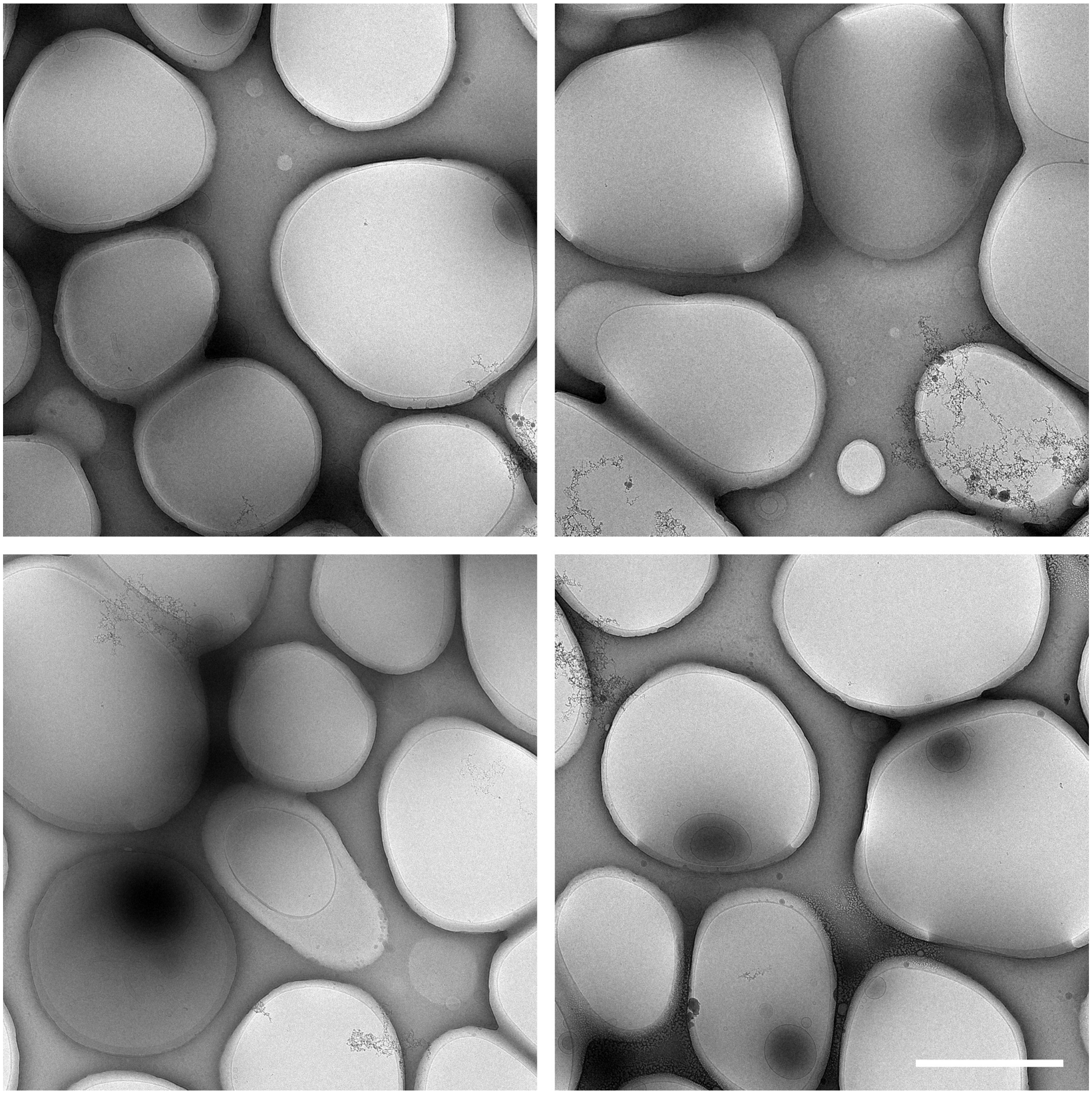

**Figure S39.** Lower magnification cryo-EM images of an unsonicated sample containing 50 mol% DOTAP, 3 mol% PTX, and the remainder DOPC. Scale bar: 2 μm.

### 50 mol% DOTAP (unsonicated)

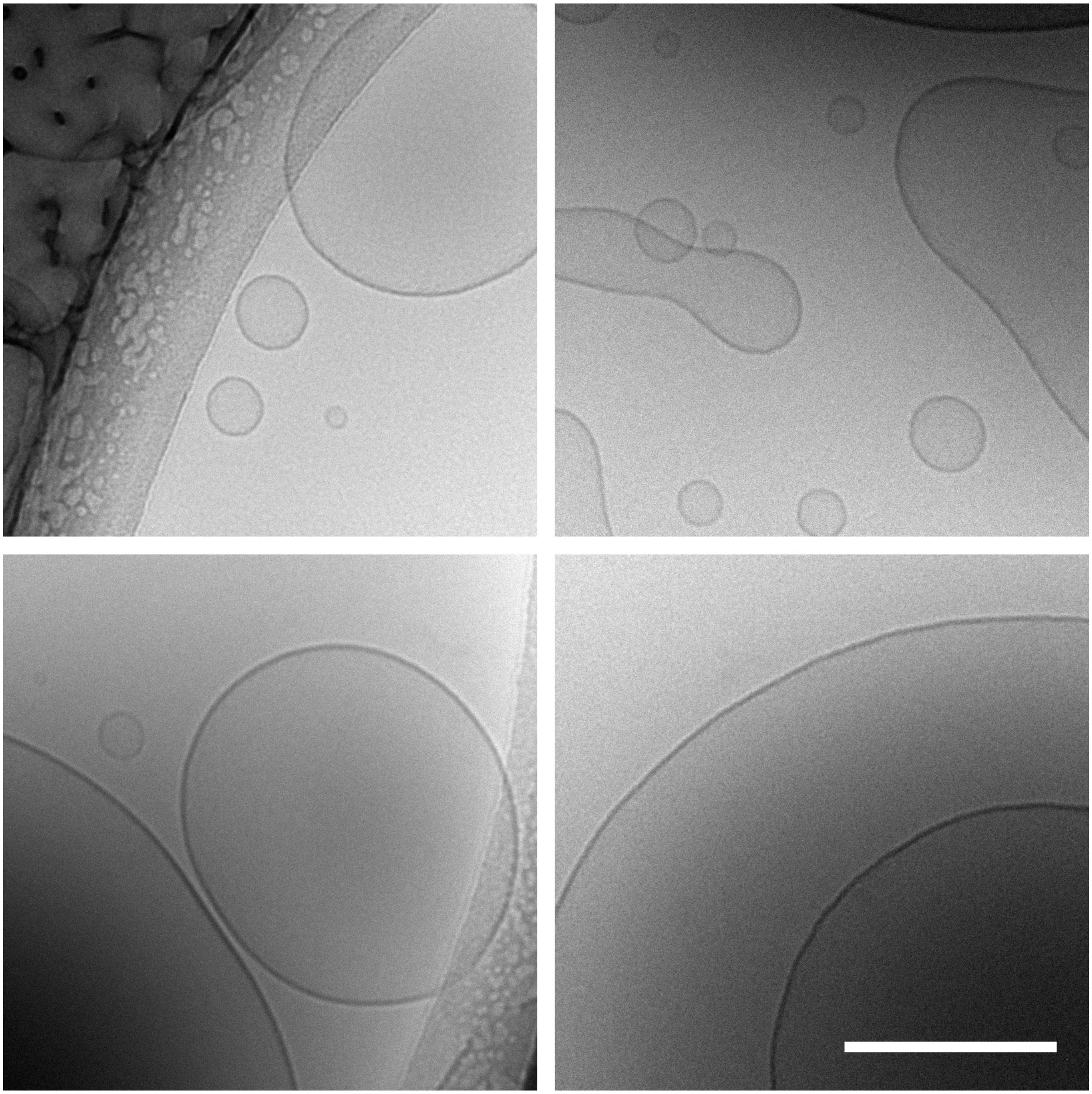

**Figure S40.** Cryo-EM images of an unsonicated sample containing 50 mol% DOTAP, 3 mol% PTX, and the remainder DOPC. Scale bar: 200 nm.

### 50 mol% DOTAP, 10 mol% PEG2K-lipid (unsonicated)

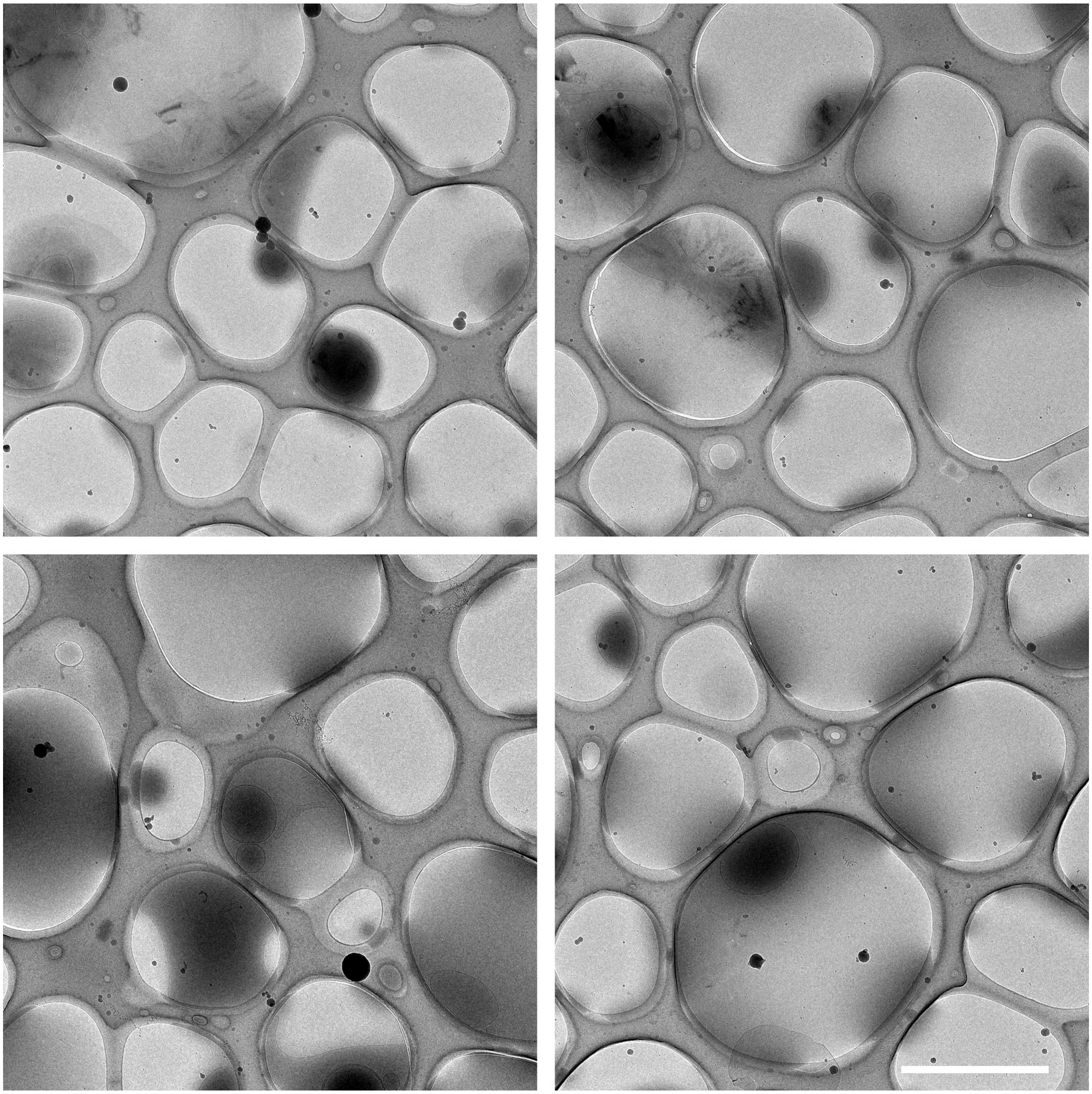

**Figure S41.** Lower magnification cryo-EM images of an unsonicated sample containing 50 mol% DOTAP, 10 mol% PEG2K-lipid, 3 mol% PTX, and the remainder DOPC. Scale bar: 2 μm.

### 50 mol% DOTAP, 10 mol% PEG2K-lipid (unsonicated)

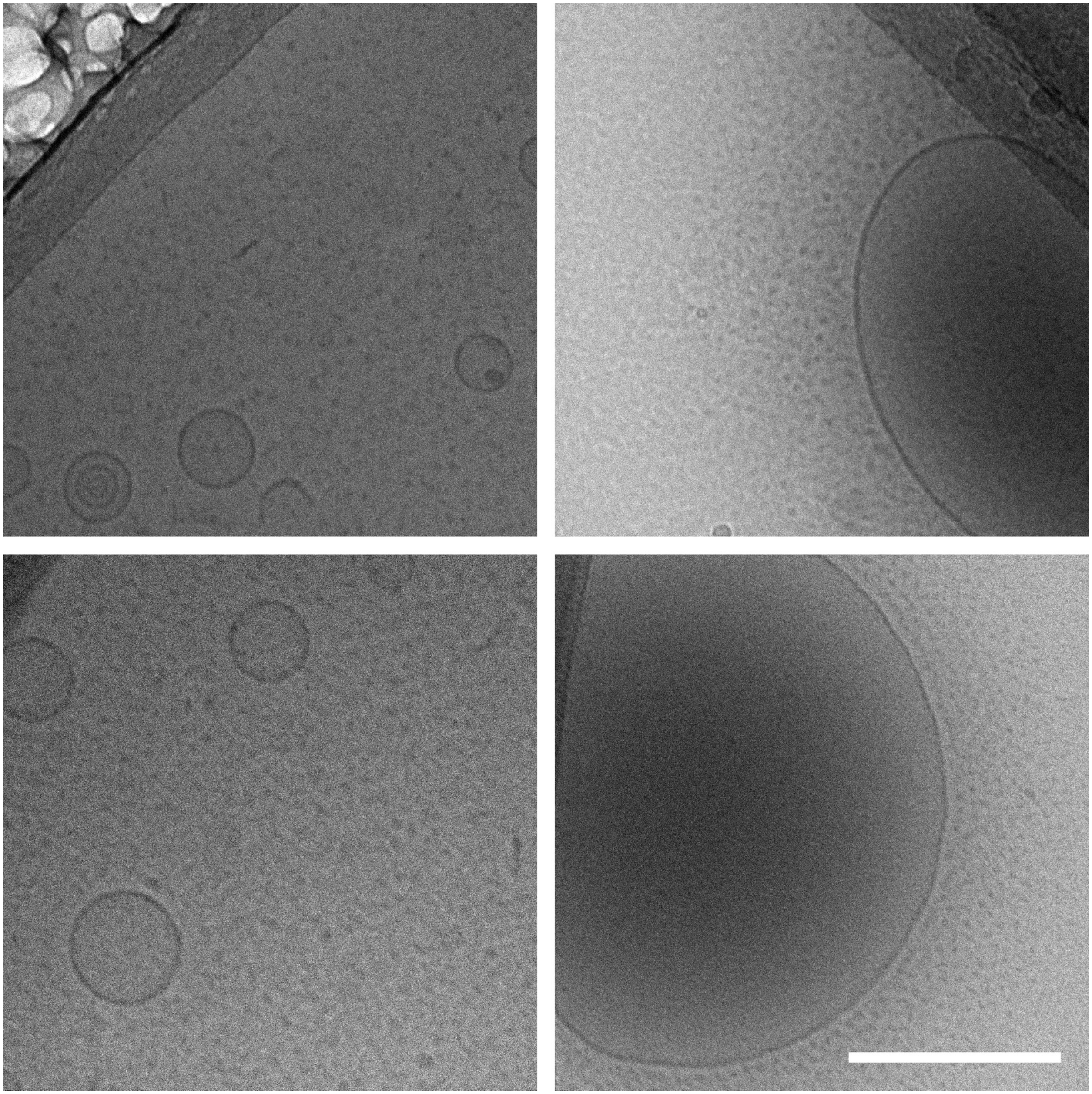

**Figure S42.** Cryo-EM images of an unsonicated sample containing 50 mol% DOTAP, 10 mol% PEG2K-lipid, 3 mol% PTX, and the remainder DOPC. Scale bar: 200 nm.

**Table S1.** Summary of sample compositions (given as molar ratios). PTX: paclitaxel.

| Å/pixel (original images) | Figures | DOTAP | MVL5 | PEG2K-lipid | PEG5K-lipid | DOPC | DOPG | PTX |
|---|---|---|---|---|---|---|---|---|
| 1.23 | 2, S8, S9 | | | | | 97 | | 3 |
| 2.46 | 2, S14, S15, S31 | 50 | | | | 47 | | 3 |
| 1.23 | 2, 5, S12, S13, S32 | | | 10 | | 87 | | 3 |
| 1.23 | S16, S17 | 80 | | | | 17 | | 3 |
| 2.46 | 3, S2, S3 | | 10 | | | 90 | | |
| 2.46 | 3, S4, S5 | | 50 | | | 50 | | |
| 2.46 | 3, S6, S7 | | 75 | | | 25 | | |
| 2.46 | 4, S10, S11 | | | 2 | | 98 | | |
| 3 | 4, 6, S18, S19 | | | 25 | | 75 | | |
| 2.46 | 4, S20, S21 | | | | 2 | 98 | | |
| 2.46 | 4, S22, S23 | | | | 25 | 75 | | |
| 2.46 | 5, S24, S25 | | 10 | 10 | | 80 | | |
| 2.46 | 5, S26, S27 | | 50 | 10 | | 40 | | |
| 3 | 5, S28–S30 | | 75 | 10 | | 15 | | |
| 1.7 | S32–S34 | 50 | | 10 | | 37 | | 3 |
| 1.23 | S32, S35, S36 | 80 | | 10 | | 7 | | 3 |
| 1.23 | 6, S37, S38 | 50 | | 25 | | 22 | | 3 |
| 2.46 | S39, S40 | 50 | | | | 47 | | 3 |
| 2.46 | S41, S42 | 50 | | 10 | | 37 | | 3 |
